# Polar Marine Microbial Communities as Reservoirs of Polyester Degrading Enzymes

**DOI:** 10.64898/2026.01.21.700866

**Authors:** Aransa Griñen, Jerónimo Cifuentes-Anticevic, Marianne Buscaglia, Pablo Vergara-Barros, Amparo Núñez Verdejo, Rodrigo Pérez-Lara, Dennys Worden-Aracena, Felipe Gatica Vargas, Felipe Engelberger, César A. Ramírez-Sarmiento, Beatriz Díez

## Abstract

**Background:** Polyethylene terephthalate (PET) is one of the most widely used plastics and a major contributor to marine pollution. While the diversity of PET hydrolases (PETases), which degrade PET into mono(2-hydroxyethyl) terephthalate (MHET), terephthalate (TPA) and ethylene glycol, has been documented in temperate and tropical waters, their potential presence in polar oceans remains unascertained.

**Results:** Here, we systematically screened polar and non-polar marine metagenomes using Hidden Markov models (HMM) generated using experimentally validated PETases. We identified >680 putative PETase-like sequences, with Antarctic and Arctic candidates enriched in high-fidelity motifs associated with PETase-like activity. Phylogenetic and structural analyses defined a high-confidence PETase-like clade comprising both Type I and Type II enzymes, differing in thermostability-related features and PET-binding motifs. Experimental assays confirmed polyesterase activity in 5/9 candidates from this clade, including polar-derived variants active at 14-25°C. Downstream enzymes for PET consumption were also widespread, detecting 209 putative MHET hydrolases and 442 TPA-catabolyzing enzymes. Further, we reconstructed 112 metagenome-assembled genomes (MAGs) carrying at least one enzyme of the PET degradation pathway, 89 of them carrying a PETase-like gene. Notably, 15 MAGs encoded multiple PETase-like enzymes, nine of them from polar datasets, and one polar MAG harbored a complete PETase-MHETase-TPA pathway, suggesting the genomic potential for a fully integrated degradation potential in cold-adapted taxa.

**Conclusions:** Together, these results demonstrate that polar oceans act as previously overlooked reservoirs of taxonomically and functionally diverse plastic-degrading enzymes. The enrichment of PETase-like enzymes and downstream pathways in polar microbial communities expands the global biogeography of plastic biodegradation and highlights cold-active enzymes as promising candidates for developing low-temperature plastic bioremediation strategies.

## INTRODUCTION

Plastics have become indispensable in modern society due to their exceptional durability, versatility, and low production cost. Global plastic production reached 413.8 million tons in 2023, and is projected to exceed 736 million tons by 2040 [1,2]. This exponential growth, coupled with inadequate waste management – 52 million tons of plastic were unmanaged in 2020 [3] – has resulted in widespread contamination of our environments, particularly affecting the oceans.

The packaging industry is one of the largest plastic consumers, and alone accounts for the majority of plastic waste in marine environments, with an estimated 4.8 to 12.7 million metric tons of plastic entering the oceans annually [4–7]. This marine plastic debris, which constitutes approximately 90% of oceanic solid waste [8], undergoes progressive fragmentation into microplastics, threatening marine ecosystems and food chains while causing significant economic losses due to poor plastic recovery via recycling [9–11].

Among synthetic plastics, polyethylene terephthalate (PET), a thermoplastic formed by petroleum-derived units of ethylene glycol (EG) and terephthalic acid (TPA), accounts for approximately 20% of global plastic production and is largely used in food and beverage packaging [4,9,12]. Current PET recycling methods rely on mechanical or chemical processes that face significant limitations, including material downgrading into non-recycled materials and energy-intensive processes, thus calling for the development of sustainable and truly circular alternatives [10,13].

As a result, enzymatic PET recycling has emerged as a promising biotechnological solution [9,14]. PET hydrolases (PETases) hydrolyze the ester bonds of PET, yielding mono(2-hydroxyethyl) terephthalate (MHET), terephthalic acid (TPA), and ethylene glycol (EG) as degradation products [9,14]. Most characterized PETases are thermophilic cutinases that function optimally near the glass transition temperature of PET (∼65°C) [15]. At this temperature, the flexibility of the PET chains increases and the enzymes have a higher probability of accessing their ester bonds for hydrolysis [16].

The landmark discovery of a secreted mesophilic PETase in the Gram-negative bacteria *Ideonella sakaiensis* 201-F6, termed *Is*PETase [17], along with a dedicated secreted MHET hydrolase (MHETase) that can further degrade this intermediate product into TPA and EG, and a set of transmembrane transporter and intracellular enzymes required for TPA catabolism [18], opened new avenues for biotechnological applications, such as enzymatic plastic upcycling [19–21]. Mesophilic enzymes offer substantial advantages for industrial applications through reduced energy requirements and operating efficiently at ambient temperatures. Subsequently, several marine-derived PETases active at 20-40°C have been identified, including PE-H from *Pseudomonas aestusnigri* VGXO14*^T^*[22] and Ple628/Ple629 from a marine microbial consortium [23,24]. Despite these advances, most studies have focused primarily on temperate marine environments [25–27], or in extreme environments such as hydrothermal vents [27], largely overlooking an in-depth characterization of high-latitude ecosystems that are increasingly affected by plastic contamination, predominantly polyesters [28–30].

The recent characterization of the cold-active PETase Mors1 from *Moraxella* sp. TA144, an Antarctic marine bacterium, as well as Mors1 homologs in Antarctic metagenomic samples [31], provides compelling evidence that polar microbes harbor unique enzymes capable of degrading synthetic polyesters at low temperatures. Such environmental conditions have been shown to drive the evolution of cold-adapted enzymes with distinct catalytic and structural features, including increased flexibility and activity at low temperatures [32,33]. In fact, increased active site flexibility has been defined as a hallmark for plastic degradation at low temperatures for *Is*PETase based on comparative molecular dynamics against thermophilic counterparts [34].

Cold-adapted enzymes are highly valuable for biotechnology, where low-temperature activity is desirable in processes requiring energy efficiency or the preservation of heat-sensitive substrates [35,36]. At the same time, plastic pollution has been increasingly documented in remote polar regions, from Arctic sea ice to Antarctic surface waters and sediments [28–30,37–39]. This highlights not only the global reach of plastic pollution but also the urgent need to understand how polar microbial communities respond and whether they contribute to plastic turnover *in situ*.

Here, we present a comprehensive metagenomic survey of PETase-like enzymes in polar marine environments, comparing them with those of temperate oceans. Using Hidden Markov Models (HMM) built from experimentally validated PETases, we screened publicly available metagenomic datasets spanning both marine regions. We characterized the diversity, distribution and abundance of putative PETase-like enzymes, classified them based on conserved functional motifs, and experimentally validated the polyesterase activity of selected candidates from a clade containing known PETases and novel PETase-like enzymes. Furthermore, we identified potential MHETases and genes encoding for TPA catabolic enzymes and transporters in 112 bacterial genomes assembled from these metagenomes (MAGs), which are required for downstream processing of PET for carbon usage.

Our findings reveal that polar oceans, long considered enzymatically cold-limited [40,41], harbor a diverse repertoire of putative polyesterases and microorganisms with potentially complete plastic degradation pathways, expanding the known ecological range of microbial plastic degradation and uncovering novel cold-active biocatalysts with direct implications for sustainable plastic recycling and bioremediation strategies.

## METHODS

### Identification of potential PETase-like enzymes in marine metagenomes

A Hidden Markov Model (HMM) profile was built from a multiple sequence alignment (MSA) of 9 known bacterial PETase amino acid sequences (Table 1) using the *hmmbuild* module of HMMER3 [42]. An additional sequence of a known PETase from *Oleispira antarctica* RB-8 (residues 37-310) was not included in building the HMM profile due to its low activity [31], but included for the phylogenetic analysis as a reference sequence.

**Table 1.** Currently known and experimentally validated PETases used as references in this study. Names of experimentally validated PETases used in this work, with the accession entry IDs according to the National Center for Biotechnology Information (NCBI) database and the UniProt database for their amino acid sequences and the PDB ID for their experimental structures. These sequences (except sequence 10) were used to build the HMM for the search of PETase-like enzymes in metagenomes. All sequences in this Table were used as references for the phylogenetic analysis.

| Seq. number | NCBI entry | UniProt ID | PDB ID | Organism | References |
|---|---|---|---|---|---|
| 1 | CAA37220.1 | P19833 | 8SPK | *Moraxella sp.* TA144 | [31] |
| 2 | GAP38373.1 | A0A0K8P6T7 | 6EQE | *Ideonella sakaiensis* 201-F6 | [17,43] |
| 3 | BAK48590.1 | F7IX06 | 6AID | *Thermobifida alba* AHK119 | [44,45] |
| 4 | CAH17553.1 | E5BBQ3 | 4CG3 | *Thermobifida fusca* KW3 | [46,47] |
| 5 | ADV92526.1 | E9LVH8 | 5LUI | *Thermobifida cellulosilytica* | [46,48] |
| 6 | AFA45122.1 | H6WX58 | —— | *Thermobifida halotolerans* | [49] |
| 7 | BAO42836.1 | W0TJ64 | —— | *Saccharomonospora viridis* | [50] |
| 8 | ACY96861.1 | D1A9G5 | 7YKO | *Thermomonospora curvata* | [51] |
| 9 | AEV21261.1 | G9BY57 | 4EB0 | *Unknown prokaryotic organism* | [52,53] |
| 10 | CCK74972.1 | R4YKL9 | —— | *Oleispira antarctica* RB-8 | [31] |

This HMM profile was then used with *hmmsearch* (e-value <1e-3) [42] to screen the Ocean Microbial Reference Gene Catalog (OM-RGC.v1, BioProject PRJEB7988) [54] via the Ocean Gene Atlas website [55], the Polar Marine Reference Gene Catalog (PM-RGC, BioProject PRJNA588686) [56], and the predicted proteins from other Antarctic samples collected in Chile Bay, Western Antarctic Peninsula (BioProject PRJNA805531 [57] and PRJNA1280935 [58]) for potential PETase-like enzymes. This permissive threshold was used for a sensitive first-pass screen, intended to avoid discarding divergent sequences given the small reference set. From the 1897 initial hits, sequences were manually curated based on the presence of the catalytic triad (*Is*PETase residues S160, D206 and H237) and/or the oxyanion hole motif (GXSXGG[GA][GA]). Sequences lacking these features were discarded, and the 683 curated sequences obtained were clustered to 80% sequence identity and coverage using CD-HIT [59] (-c 0.8 -s 0.8 -l 150). Proteins, whether complete or partial, were retained if they were at least 150 amino acids long and contained the conserved residues required for motif classification. For the Antarctic metagenomes assembled in this study, ORF completeness was assessed from the Prodigal predictions; approximately 70% of the retained PETase-like candidates were predicted to contain both start and stop codons. ORF-completeness information was not available for sequences obtained from the OM-RCG.v1 and PM-RGC catalogs. The resulting 442 dereplicated sequences were therefore classified, with modifications as described in Supplementary Table S1, according to their conserved residues into previously described motif groups M0, M3, M3 strict, M4, M4 strict, and M5 (Supplementary Table S2). This classification was adapted from the criteria previously established for PETases identification in marine and terrestrial metagenomes [25,26]. Our adaptations included an M0 category defined by the catalytic triad, relaxed and strict variants of M3 and M4 with different residue and inter-motif spacing requirements, a strictly conserved catalytic histidine, and expanded residue tolerance at selected positions in the C-terminal motif. The motif criteria were applied additively, with each group requiring the criteria of the preceding group. Lastly, BLASTp [60] was used to calculate the sequence identity and coverage of the retrieved sequences relative to the reference known PETases in Table 1.

### Taxonomic assignment, subcellular localization prediction, and phylogenetic analysis of PETase-like enzymes

Taxonomic assignments for PETase-like enzymes were inferred by aligning their sequences against proteins predicted from GTDB R220 database [61] genomes using DIAMOND v2.0.13 [62]. Each PETase-like enzyme was assigned the taxonomy of the genome encoding its closest matching protein (Supplementary Table S2 and Supplementary Fig. S1). Additionally, the subcellular location of the enzymes was predicted using ayu v0.2 [63] (Supplementary Table S2 and Supplementary Fig. S1). To reconstruct PETases phylogeny, an MSA was then performed with MUSCLE v5.3 using the PPP algorithm and stratified alignment option (-align -stratified) [64], including the 10 known PETase sequences as references (Table 1) and 11 dienolactone hydrolase domain sequences (DHL; Pfam PF01738) as outgroup. The resulting MSA was resampled with a minimal letter confidence of 0.1 and a maximum gap fraction 0.5, and the maximum confidence MSA was subsequently extracted with MUSCLE v5.3 (-maxcc). A maximum likelihood phylogenetic reconstruction was then performed with IQ-TREE v.2.0.7 [65], first finding the best evolutionary model including FreeRate heterogeneity models for constructing the phylogenetic tree with ModelFinder [66] and then running 10,000 ultrabootstrap and ultrafast jackknife replicates (-m TESTNEW -B 10000 -J 10000) to assess branch support in the tree [65,67,68]. The resulting phylogenetic tree was visualized with TreeViewer [69].

### Protein structure prediction of metagenomic PETase-like enzymes

A total of 9 potential metagenomic PETase-like enzymes (MetaG1 to MetaG9) were selected from the sequences that clustered together with the known reference PETases (hereafter, high-confidence PETase-like clade). The sequences were first subjected to analysis of signal peptides for extracellular secretion using SignalP v5.0 [70] (present in MetaG1-G5), and then to prediction of disordered regions after the signal peptide using PrDOS [71]. Based on these predictions, the MetaG1-MetaG9 enzymes were truncated in their N-terminus, with the final sequences shown in Supplementary Table S3. The structures of the selected, truncated PETase-like enzymes were predicted on ColabFold v1.5.5 [72], an AlphaFold2 [73] implementation on Google Colaboratory, using default parameters (1 seed, 5 models, 3 recycles, no templates, alphafold2_ptm model for monomers, MSAs generated with MMseqs2 [74] search against UniRef100 [75] and the ColabFoldDB environmental database [72]). The best model out of the 5 predictions was further selected for structural analysis of sequence motifs indicative of potential polyesterase activity (Supplementary Fig. S2). The resulting structures were analyzed and visualized with PyMol v3.1.0 [76].

### Evolutionary trace analysis of PETase-like enzymes

A real value evolutionary trace (rvET) analysis of amino acid sequence conservation normalized by the length of the sequences in the MSA [77] was performed on 33 sequences from the high-confidence PETase-like clade derived from the phylogenetic reconstruction (sequences *gene_956802*, *gene_9327*, *gene_870527*, and *NODE_94302* were removed due to their shorter length). This analysis was performed using the Universal Evolutionary Trace server [78] based on an MSA generated using Clustal Omega [79], which enables the use of structural alignment-based MSAs as profiles to improve the quality of the alignment. The structure-based MSA profile was generated using the chain A of all structures of the known PETases listed in Table 1 as input for the STAMP structural alignment algorithm [80] available in the MultiSeq extension [81] of VMD v1.9.4 [82]. The structure-based MSA was used as a profile (--p1) to align the sequences of the high-confidence PETase-like clade using Clustal Omega [79] with 3 refinement iterations over the full alignment and using the full distance matrix (--iter=3 --full --full-iter). The resulting MSA was analyzed using the Universal Evolutionary Trace server [78] in full or split into Type I and Type II enzymes [83].

### Identification of potential enzymes involved in the PET degradation pathway

MHETase-like enzymes were annotated using a custom HMM profile constructed from 3 known amino acid sequences with confirmed MHETase activity (Supplementary Table S4) using the *hmmbuild* module of HMMER3 [42]. Using *hmmsearch* (e-value <1e-3) [42], we screened the OM-RGC.v1 (Ocean Gene Atlas website) [55], the PM-RGC [56], and the predicted proteins from metagenomes obtained from Chile Bay, Western Antarctic Peninsula [57,58] for potential MHETase-like enzymes. Resulting sequences were manually curated by identifying the catalytic triad (*I. sakaiensis* MHETase: S225, D492, H528), a conserved oxyanion hole residue (G132) and the disulfide bond near the catalytic residues (C224-C529 in *I. sakaiensis* MHETase) [84].

Additionally, potential TPA-degrading enzymes (TphA1, TphA2, TphA3, TphB) were identified using 4 custom HMM profiles based on the protein sequences from Supplementary Table S5, generated using the *hmmbuild* module of HMMER3 [42]. These enzymes were searched in the same databases as MHETase-like enzymes using *hmmsearch* (e-value <1e-3) [42]. Resulting sequences were curated by length, retaining only those within ±10% of the average reference sequence length. Sequences were further filtered by BLASTp [85], keeping only those with >50% identity and >60% coverage relative to reference sequences.

### Abundance of PET degradation pathway enzymes in global ocean metagenomes

The abundance of potential PETase-like, MHETase-like, and TPA-degrading enzymes was determined in marine samples from both polar (38 Arctic and Antarctic metagenomes from PM-RGC; hereby “Polar-MetaG” [56,57]) and temperate (59 metagenomes from the TARA Oceans project OM-RGC.v1; hereby “Temperate-MetaG” [54]) regions. Reads from the Polar-MetaG were processed with bbduk from BBTools [86] to remove adapters and low quality bases. Quality filtered reads from Polar-MetaG and Temperate-MetaG were aligned to the non-redundant set of potential PETases, MHETases and TPA-degrading enzymes using DIAMOND [87] in BLASTx sensitive mode, using a e-value <1e-5 and a sequence identity and coverage cutoff of 80% and 70%, respectively (-e 1e-5 --query-cover 70 --id 80 --sensitive -l 10). The number of aligned reads per sample was normalized by sequencing depth and gene length to calculate the reads per kilobase per million mapped reads (RPKM) using R [88].

### Identification and abundance of marine metagenome-assembled genomes (MAGs) containing enzymes involved in PET degradation

Contigs from individual Chile Bay Antarctic metagenome assemblies (>1,000bp) [57] were grouped into bins using the metaWRAP binning module, which incorporates: metaBAT2 v2.12.1 [89], MaxBin2 [90] and CONCOCT v1.1.0 [91], with default parameters. The outputs from these three individual binning tools were combined using the metaWRAP bin_refinement module, and bins with >50% completeness and <10% contamination according to CheckM v1.0.18 [92] were selected for further analysis. This set of MAGs, along with those previously generated from the other Polar-MetaG [56] and Temperate-MetaG [93], were dereplicated using dRep (v3) [94] with default parameters. Proteins were predicted in these sets of MAGs with Prodigal (-p meta -n) [95], and enzymes involved in the PET degradation pathway were identified using BLASTp against a non-redundant version of the previously annotated enzymes (>80% identity and >80% coverage). MAG taxonomic annotation was performed using the GTDB-TK classify_wf module [96] (GTDB Release 95, [61]). A phylogenetic reconstruction of the dereplicated MAGs containing at least one PET degradation pathway enzyme was performed using 120 concatenated single-copy bacterial genes from the GTDB-TK intermediate files, and ten Aquificota MAGs were included as an outgroup. The MSA for the phylogenetic reconstruction was performed with MAFFT [97] using default parameters (--auto) and the maximum likelihood tree was constructed with IQ-TREE v.1.5.5 [98], again finding the best evolutionary model for constructing the phylogenetic tree with ModelFinder [66] and then running 10,000 ultrabootstrap and 10,000 Shimodaira-Hasegawa approximate likelihood ratio test to assess branch support in the tree (-m TESTNEW -bb 10000 -alrt 10000) [67,68,99]. The resulting tree was visualized with iTOL [100].

The abundance of MAGs containing at least one PET degradation pathway enzyme in the 112 metagenomes from polar and temperate environments was determined by read mapping with coverM (https://github.com/wwood/CoverM) [101]. Briefly, quality-filtered paired-end reads were mapped using minimap2 [102] for short genomic reads (-x sr) with a minimum alignment of 90% and minimum identity of 95% [103]. Reads were required to map as proper pairs and supplementary alignments were excluded. MAGs with a coverage >1x across at least 75% of their genome were considered present; otherwise, the abundance was considered zero.

### Gene synthesis and cloning of candidate PETase-like enzymes

Codon-optimized synthetic genes encoding 9 potential PETase-like enzymes (MetaG1 to MetaG9) for expression in *E. coli*, selected from the high-confidence PETase-like clade and truncated on their N-terminus based on the predictions from SignalP v5.0 [70] and PrDOS [71], were codon-optimized and synthesized by Genscript (Piscataway, NJ, USA) and cloned into the pET-28a(+)-TEV plasmid between the NdeI and BamHI restriction sites. This vector allows for heterologous expression in *E. coli* BL21(DE3) cells under the control of the inducible *lac* promoter by addition of isopropyl β-D-1-thiogalactopyranoside (IPTG), and incorporates a His-tag for purification by immobilized metal affinity chromatography (IMAC) and a TEV protease cleavage site for optional removal of the His-tag in the N-termini of the enzyme of interest.

For controls of enzymatic activity, we used as positive control the gene encoding for PHL7 cloned into the pET-26b(+) plasmid from our previous work [104] and ordered the readily codon-optimized synthetic gene encoding LCC^ICCG^ from previous works [105] cloned into the pET-24b(+) plasmid from Genscript. In both cases, the genes were cloned between the NdeI and XhoI restriction sites, which incorporates a C-terminal His-tag for IMAC. For the negative control, we used the TEM 1 beta-lactamase cloned into the pET-28a(+) plasmid.

Based on the predictions from ColabFold [72] of the three-dimensional structure of MetaG4 and on the analysis of the sequence of MetaG4 using CD-Search [106], which identified a ricin B-type lectin carbohydrate binding module (CBM) in its C-terminal region (residues 290-411), two variants of this enzyme were ordered: MetaG4, in which residues 261-415 were truncated from the final sequence, and MetaG4-CBM, which corresponds to the full-length sequence of MetaG4 that includes the CBM. The nucleotide and amino acid sequences of all enzymes used in this work for experimental characterization are presented in Supplementary Table S3.

### Preparation of PCL nanoparticle suspension

To prepare a polycaprolactone (PCL) nanoparticle suspension, 250 mg of amorphous PCL (Sigma-Aldrich, MI, USA) were dissolved into 10 ml acetone at 50°C. The solution was then poured into 100 ml of milliQ water under vigorous stirring and filtered through a paper filter. Residual acetone was evaporated by overnight incubation at 50°C. To determine the concentration of PCL nanoparticles in the final suspension, three 1 ml aliquots were centrifuged, the supernatants were discarded, and the pellets were dried overnight and weighted. The concentration (in mg/ml) was calculated as the mean mass of the three pellets obtained from each 1 ml sample of PCL nanoparticle suspension.

### Plate-clearing polyesterase activity assays

To assess the polyesterase activity of the 9 selected PETase-like enzymes, PCL agar plates were prepared as previously described [26,107]. Briefly, a suspension of PCL nanoparticles was added to autoclaved agar (1% w/v PCL) at 60°C under continuous stirring. The medium was then supplemented with kanamycin (50 μg/ml) for selection based on the resistance encoded by the plasmids. The agar was poured into Petri dishes that were left open for 30 minutes to allow residual acetone to evaporate.

*E. coli* BL21(DE3) cells were transformed with plasmids encoding each metagenomic PETase-like enzyme (MetaG1-MetaG9), LCC^ICCG^ and PHL7, and grown on LB agar plates overnight at 37°C. Then, colonies were selected and inoculated in 20 ml of LB media supplemented with 50 µg/ml kanamycin until reaching an optical density (OD_600_) of 0.6-0.8, upon which expression was induced overnight with 0.3 mM IPTG at 37°C. The following day, cells were harvested by centrifugation at 5,000 rpm for 30 minutes, and the resulting pellet was weighed. Each cell pellet was resuspended with 1 ml of lysis buffer (20 mM sodium phosphate pH 8.0, 200 mM NaCl) per 100 mg of pellet to obtain samples with a similar amount of resuspended cells, and then sonicated on ice (Qsonica Q125, USA) to obtain the crude extracts. Lastly, a total of 5 μl of crude extract were inoculated onto the PCL-containing agar plates and incubated at 14°C, 25°C, 37°C and 50°C for up to two days. Photographs were taken daily to record the PCL nanoparticle degradation progress.

### Protein expression tests

Plasmids encoding for the MetaG1-MetaG9 proteins transformed in *E. coli* BL21(DE3) were subjected to soluble protein expression tests. For each variant, 5 ml of *E. coli* culture were grown in LB media supplemented with 50 µg/ml of kanamycin overnight at 37 °C and 200 rpm of agitation. These cultures were used for starting 25 ml inoculums with OD_600_ lower than 0.1 in LB media supplemented with 50 µg/ml of kanamycin, which were incubated at 37 °C and 200 rpm of agitation. When the cultures reached OD_600_ between 0.6 - 0.8, they were separated into 5 ml aliquots and 0.3 mM of IPTG was added for induction of protein expression. Each aliquot was incubated for 16 h at 37 °C or 18 °C at 200 rpm of agitation. The next day, the tubes were centrifugated at 10,000 rpm and 4 °C for 10 minutes and pellets were stored at -80 °C until use.

Cell lysis was carried out by resuspending the pellets in 1 ml of lysis buffer B (20 mM Tris-HCl pH 8.0, 300 mM NaCl, 5 mM 2-mercaptoethanol and 10% glycerol) and sonicated on ice using a sonicator (Qsonica Q125, USA) for 90 s (1 s ON, 2 s OFF, 40% amplitude). The lysates were centrifugated at 13,000 rpm for 10 min at 4 °C, the supernatant was collected and the resulting pellets were resuspended in 8 M Urea. Then, each lysate was purified with a simplified Ni-NTA resin (Cytiva, USA) protocol. Briefly, 250 µl of lysate were incubated in 50 µl of resin previously equilibrated with lysis buffer B in an Eppendorf tube for 1 h at 4 °C, then the resin was centrifuged for 3 min in a benchtop centrifuge, the supernatant was discarded and the resin was washed with lysis buffer B supplemented with 25 mM imidazole (Sigma-Aldrich, USA), the supernatant was discarded again and the proteins were eluted with lysis buffer B supplemented with 250 mM imidazole. Finally, 15 µl of both the eluted supernatant and the urea-resuspended pellets were loaded into a 12% SDS-PAGE gel. For all proteins, a negative control of a cell lysate without IPTG was used.

For the proteins encoding active proteins according to the PCL plate clearing assays, the plasmids encoding these proteins were also transformed in *E. coli* Rosetta-Gami 2 pLysS (DE3) [RG2] and Origami 2 pLysS (DE3) [OR2], and the same protein expression tests were performed.

### Enzyme purification of solubly expressed proteins

Solubly expressed variants based on our protein expression tests were induced in 1 L of LB media cultures using the same conditions defined by the expression tests. Briefly, a pre-inoculum of MetaG1 transformed in *E. coli* OR2, MetaG3 in *E. coli* OR2, MetaG4 and MetaG4-CBM in *E. coli* RG2, and MetaG6 in *E. coli* RG2 were grown in 15 ml of LB media supplemented with 50 µg/ml of kanamycin and 25 µg/ml chloramphenicol at 200 rpm and 37 °C for 16 h. Then, the overnight grown cells were inoculated in 1 L of LB media supplemented with 50 µg/ml of kanamycin and 25 µg/ml chloramphenicol at 200 rpm until reaching an OD_600_ between 0.6 - 0.8, after which protein overexpression was induced by the addition of 0.3 mM IPTG and the cells were grown at 200 rpm and 18 °C for 16 h.

The next day, the cell cultures were centrifugated at 5,000 rpm for 30 min at 4°C and resuspended in 50 ml of lysis buffer B supplemented with 0.25 mg/ml of lysozyme (US Biological, USA) and 5 mM Phenylmethylsulfonyl fluoride (PMSF) (Thermo Scientific, USA) and incubated at room temperature for 30 min. Then, cells were sonicated on a QSonica Q125 sonicator for 45 min (4 s ON, 8 s OFF, 40% amplitude), and the cell lysates were centrifugated for 30 minutes at 11,000 rpm and 4 °C. The resulting supernatant was loaded into 1 ml HisTrap Ni-NTA columns (Cytiva, USA) previously equilibrated with 10 column volumes of lysis buffer B, washed twice with 10 column volumes of lysis buffer B supplemented with 25 mM imidazole and the enzymes were finally eluted using 5 column volumes of lysis buffer B supplemented with 250 mM imidazole, collecting the eluted fractions every 500 µl. The eluted fractions were pooled and concentrated to 0.5-1 ml with 30 kDa 15 ml Amicon conical concentrations (Sigma Aldrich, USA) by centrifugation at 7,500 g and 4°C. The concentrated enzymes were filtered with 0.22 µM PES filters and subjected to a second round of purification by size exclusion chromatography at a flow rate of 0.5 mL/min using a Jasco LC-4000 HPLC (Jasco Inc., Japan) using a Superdex 75 Increase column (Cytiva, USA) and lysis buffer B as the mobile phase. Final protein concentration after size exclusion chromatography was determined by Bradford analysis (Thermo Fisher Scientific, USA).

### Enzymatic assays against PCL nanoparticles

The activity of MetaG3 and MetaG6 against PCL nanoparticles was determined by the decrease in turbidity of the nanoparticles due to enzymatic hydrolysis as described in previous works [31]. Briefly, enzymatic hydrolysis reactions in 125 mM sodium phosphate buffer pH 8.0 and 200 mM NaCl supplemented with a fixed concentration of MetaG3 (8 µg/ml) or MetaG6 (4 µg/ml) in 96-well microplates were started with the addition of 0.79 mg/mL PCL nanoparticles up to a volume of 200 µl, and the decrease in OD_600_ was monitored on a EPOCH2 spectrophotometer (BioTek Instruments, USA) at 2 s intervals for 10 minutes. Initial hydrolysis rates were calculated from the linear region of the OD_600_ versus time curves from three technical replicates.

To determine the optimal reaction temperature for each enzyme, initial hydrolysis rates were measured at room temperature (20 °C), 30, 40, 50 and 60 °C. The temperature yielding the highest activity from three technical replicates was selected as the optimal temperature for subsequent kinetic characterization.

Enzyme kinetic assays for MetaG3 and MetaG6 against PCL nanoparticles at their optimal PCL degradation temperature were performed in the same reaction buffer conditions as the previous experiments by measuring the initial enzymatic hydrolysis rates of PCL at a fixed substrate concentration of 0.79 mg/mL, while varying the enzyme concentration from 1 to 30 µg/mL, performing the experiments in three technical triplicates. Assays for MetaG3 and MetaG6 were performed at 40 °C 50 °C, respectively. Reactions were initiated by the addition of PCL nanoparticles and the decrease in OD_600_ was monitored at 4 s intervals for 10 minutes. Initial hydrolysis rates were calculated from the linear region of the OD_600_ vs. time curves. The kinetic parameters were determined by fitting the initial hydrolysis rate data to a pseudo-first-order heterogeneous catalysis equation [51,108].

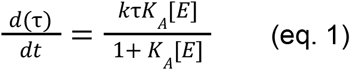

With *τ* as the turbidity of a nanoparticle suspension, *t* as the reaction time, *k_τ_* as the hydrolysis rate constant based on the turbidity change, K_A_ as the adsorption equilibrium constant, and [E] as the enzyme concentration.

### Statistical tests

Statistical differences among PETase motifs, sample regions and depths, and all the degradation pathway enzymes, were assessed by Multifactorial ANOVA and Tukey’s post hoc tests using the Minitab software [109].

## RESULTS

### Identification of new putative PETase-like enzymes

By leveraging a curated HMM built from 9 experimentally validated PETases [26] (Table 1), we systematically screened publicly available marine metagenomic datasets including high-latitude regions (Fig. 1) to uncover previously undetected enzymes with potential polyesterase activity in these regions.

**Fig. 1.**
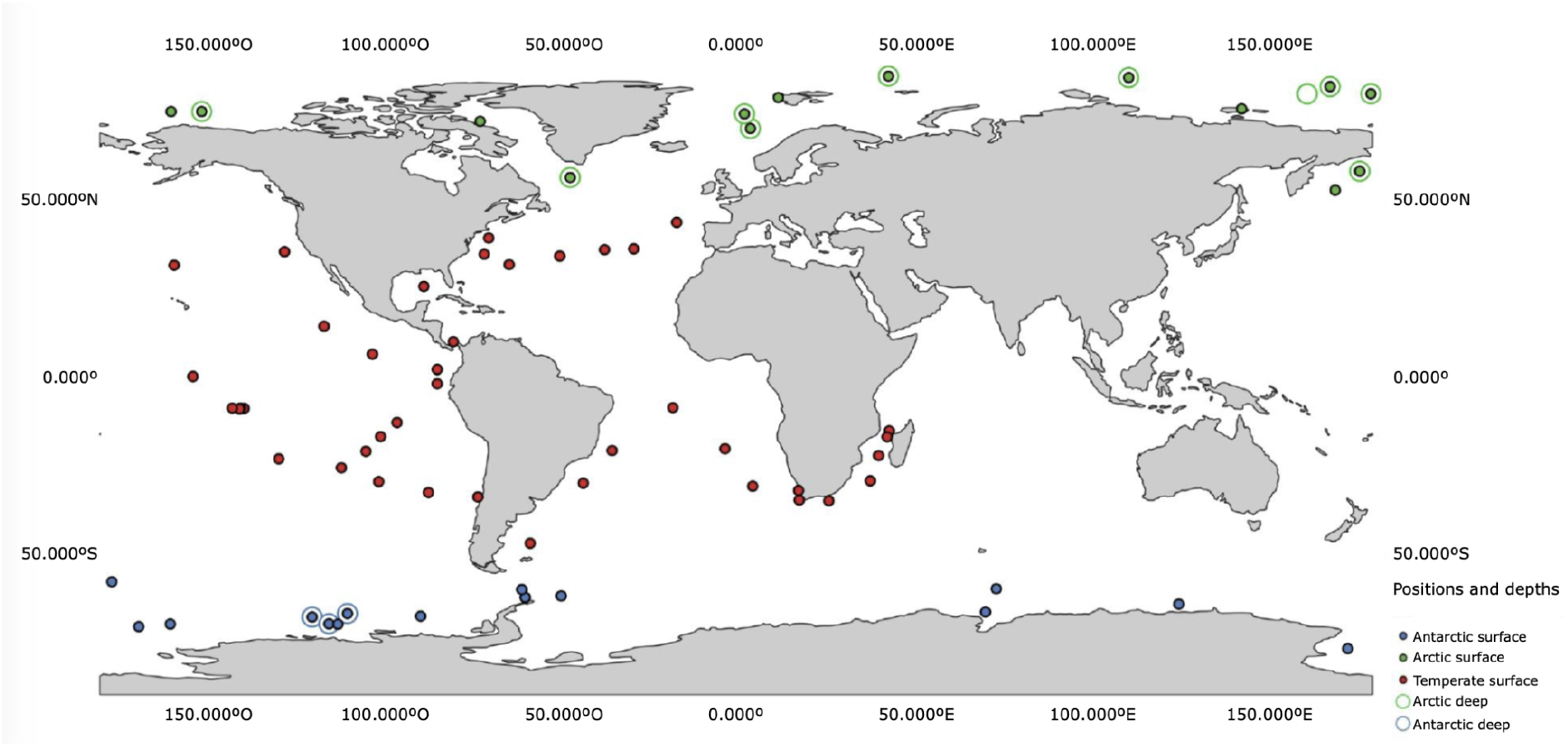
Distribution of metagenomic samples used to search for potential marine PETase-like enzymes. World map showing the geographical distribution of samples from the three metagenomic sources analyzed in this work: OM-RGC.v1, PM-RGC, and Chile Bay (Western Antarctic Peninsula). The color code describes the temperate (red), Arctic (green) or Antarctic (blue) location of the samples, while the symbol code indicates whether the sample is from the water surface (0-100 m, filled circles) and/or deep ocean (empty symbols, >100 m).

We identified 683 putative PETase-like sequences, which, together with previously characterized PETases (Table 1), were classified into six conserved motif groups (M0, M3, M3 strict, M4, M4 strict, and M5) [25,26]. Specifically, M0 requires only the catalytic triad (Ser-Asp-His), and motif complexity increases from M0 to M5 by sequentially incorporating additional PETase functional signatures, including the oxyanion hole and aromatic clamp, together with progressively more constraints on sequence context and motif architecture (Supplementary Table S1).

A maximum likelihood phylogenetic reconstruction of 442 sequences longer than 150 amino acids was performed, revealing a well-supported monophyletic clade, referred to as the high-confidence PETase-like clade (Fig. 2). This clade contained all 10 known PETases used as reference sequences from Table 1 and 27 novel M5 motif-containing amino acid sequences, 21 from polar datasets and 6 from temperate datasets.

**Fig. 2.**
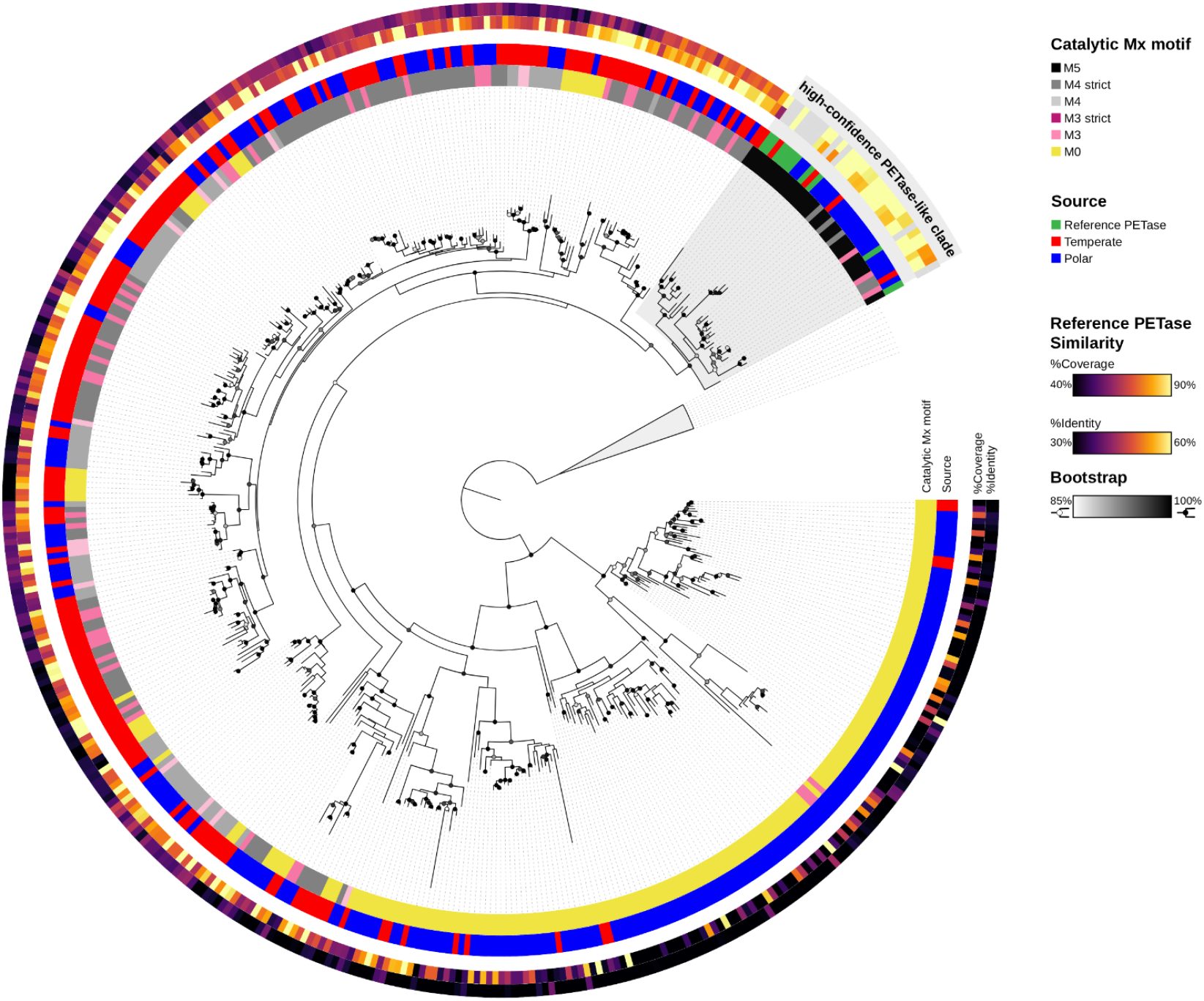
Diversity of PETase-like sequences from marine metagenomes. Phylogenetic reconstruction of potential PETase-like sequences identified in marine metagenomes, generated using maximum likelihood. From the center outwards, the first ring represents their classification according to the different PETase annotation motifs (Supplementary Tables S1 and S2), and the second ring indicates the environmental source of the sequences. The following two rings indicate the similarity to the reference by coverage and identity percentage, respectively. Bootstrap values ≥ 85% are denoted in grey scale.

The presence of polar sequences in this clade supports the hypothesis that cold-adapted PETase-like enzymes may share conserved sequence and structure features with their experimentally validated mesophilic and thermophilic counterparts, potentially maintaining enzymatic activity under psychrophilic conditions. In contrast, sequences outside this clade, particularly those from M0 and M3 groups, may represent more general esterases with limited PET specificity [34,110].

To further characterize these 442 putative PETase-like sequences, we determined their taxonomic affiliations and predicted their subcellular localization. Most sequences were affiliated with Bacteroidota (41.4%) or Pseudomonadota (32.8%), and 73.7% were predicted to localize outside the cytoplasm, either in the periplasm (49.3%) or extracellularly (24.4%), while 26.2% were predicted to be cytoplasmic (Supplementary Fig. S1 and Supplementary Table S2). This is consistent with the extracellular localization required to access an insoluble substrate such as PET, or with the periplasmic destination that is known to provide the oxidative environment that facilitates the critical disulfide bond formation of PETases [111].

### Polar regions as reservoirs of Type I and Type II PETase-like enzymes

We determined the distribution and abundance of the 442 newly identified putative PETase-like enzymes across surface and deep marine waters (Fig. 1), classifying them according to the 6 previously described catalytic motifs [25,26] (Supplementary Table S1).

Our results revealed a statistically higher abundance of sequences bearing the higher complexity motifs M4, M4 strict, and particularly M5 in polar marine regions, compared to temperate waters, a pattern that was especially pronounced for M5 motif sequences in deep Antarctic metagenomes (Fig. 3 and Supplementary Table S6).

**Fig. 3.**
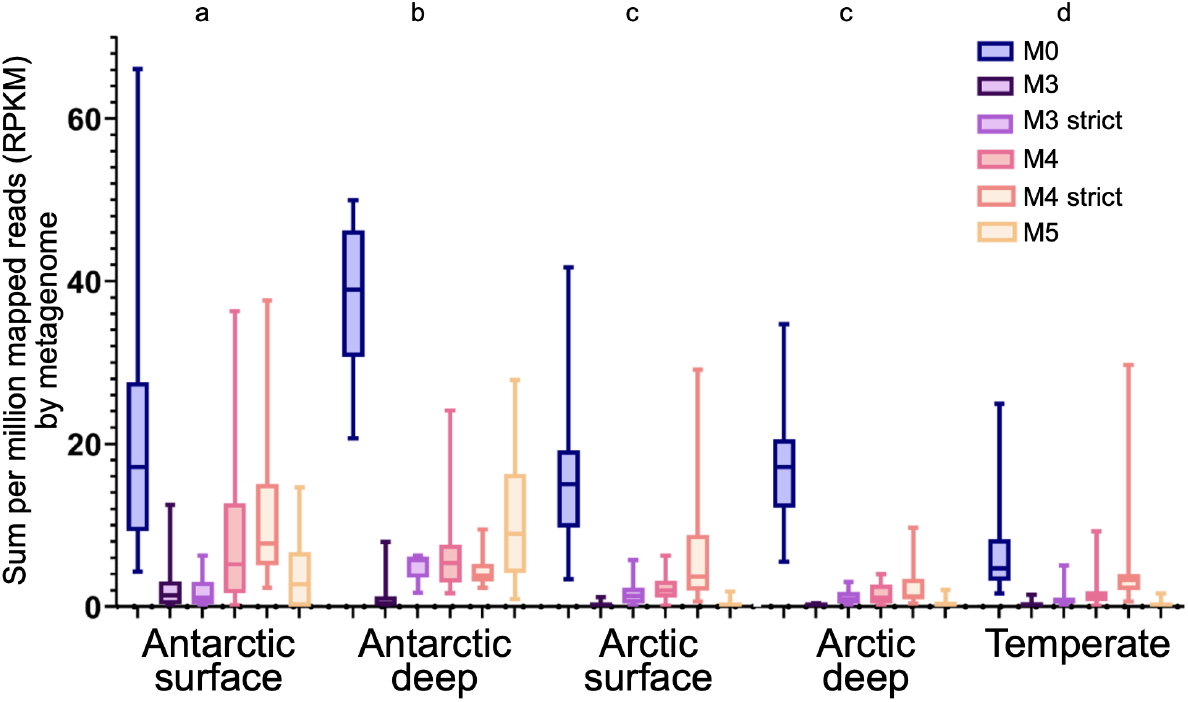
Abundance of putative PETase sequences in marine metagenomes. Bar graph representing the abundance in reads per kilobase per million mapped reads (RPKM) of potential PETase-like sequences identified in marine metagenomes, classified according to PETase annotation motifs (Supplementary Table S1). Letters a, b, c and d represent significant differences between motif groups in each area and depth, based on multifactorial ANOVA with Tukey’s a posteriori test (Supplementary Table S6).

To further investigate evolutionary relationships among these enzymes, we conducted a phylogenetic reconstruction of the high-confidence PETase-like clade, which includes both reference PETases and the 27 novel metagenome-derived sequences from polar and temperate marine environments containing mainly the M5 motif. This clade was clearly resolved into two phylogenetic groups (Fig. 4), which can be classified into Type I and Type II PETases, as previously described [83]. These types are distinguishable by the absence or presence of an additional active site disulfide bond (formed by C203 and C239 in *Is*PETase). Type I enzymes, typically thermophilic, lack this covalent interaction. In contrast, Type II enzymes, prevalent in mesophilic and psychrophilic organisms, include this disulfide bond and also an extended active site loop, which enables higher flexibility of the active site loops at room temperature compared to thermophilic counterparts, thus allowing the degradation of PET at moderate temperatures [31,34,83]. Based on the topology of the tree and the presence or absence of the active site disulfide bond [31,83], we identified 23 novel Type II and 4 novel Type I putative PETases. Within this clade, 21 of the 27 sequences originated from polar datasets, and 19 of these corresponded to Type II enzymes. Polar marine environments thus emerge as a substantial source of Type II PETase-like candidates.

**Fig. 4.**
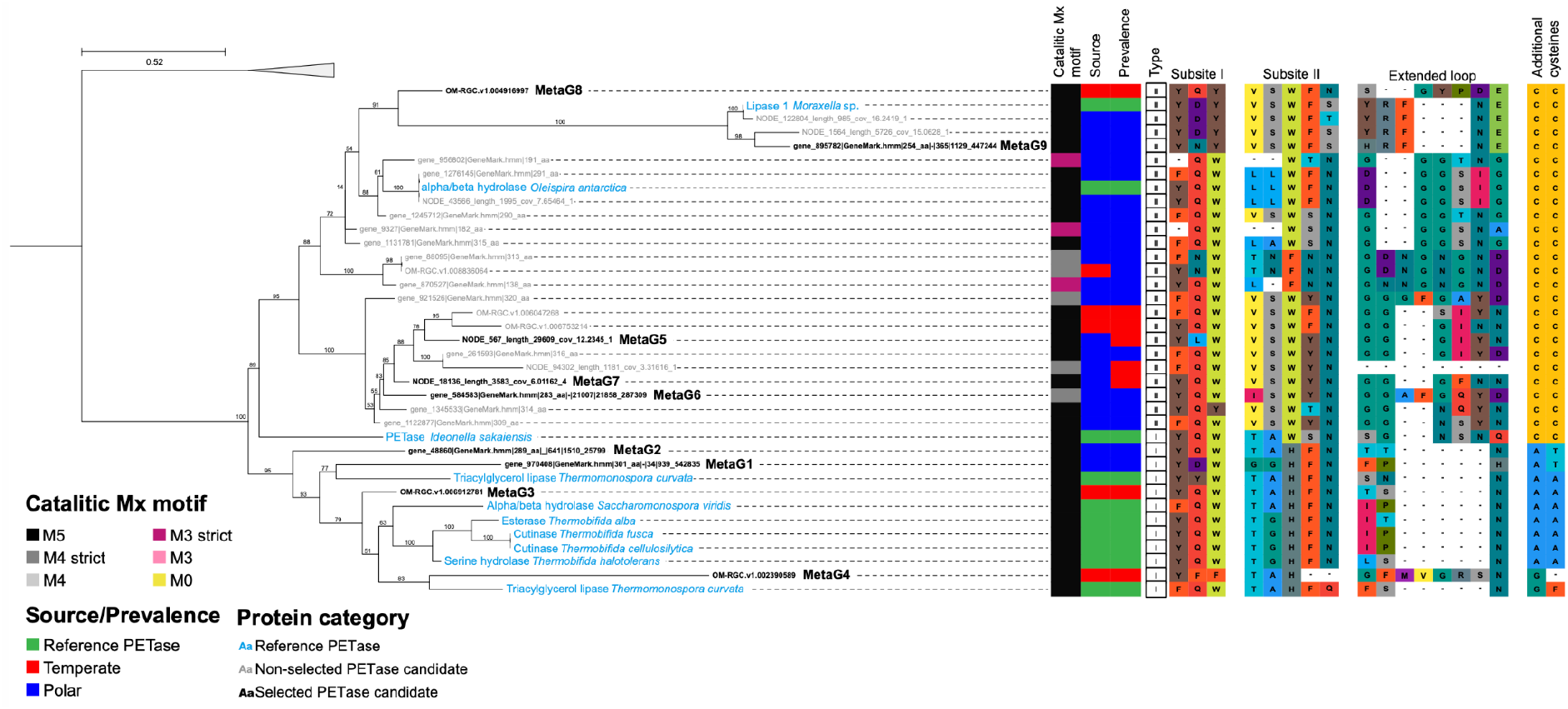
Conservation and diversity of sequences from the high-confidence PETase-like clade. Truncated phylogenetic reconstruction of the high-confidence PETase-like clade, including both PETase-like sequences identified in the metagenomes (black) and reference enzymes (light blue). The closest clade from the phylogenetic reconstruction in Fig. 2 was used as an outgroup. Sequences selected for experimental validation are highlighted in bold. For all sequences, the Mx motif, environmental source, prevalence (polar or temperate enrichment), and the amino acid residues at conserved positions of the catalytic triad, subsite I and II, the extended loop and additional cysteines involved in the formation of an active site disulfide bond are shown.

We then quantified the sequence conservation of 33 of the enzymes (see *Methods*) in the high-confidence PETase-like clade via rvET analysis (Fig. 5). This analysis was performed either separately for Type I and Type II enzymes (Fig. 5A), or for the whole clade (Fig. 5B and 5C), to enable potential identification of type-determining residues in these PETase-like enzymes.

**Fig. 5.**
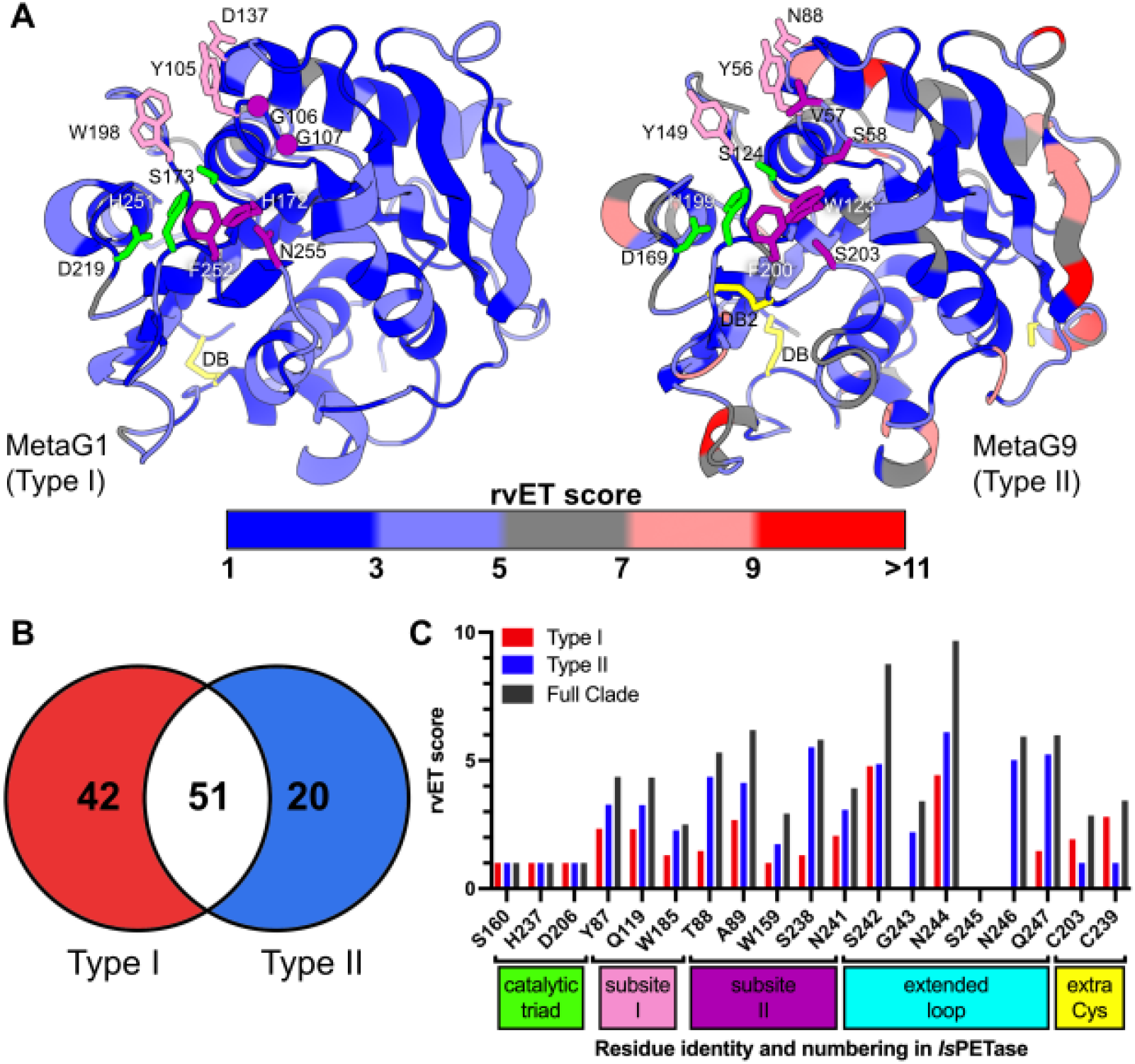
rvET analysis of enzymes in the high-confidence PETase-like clade. A. Conserved residues in the enzymes from the high-confidence PETase-like clade, visualized on cartoon representations of the best ColabFold-predicted structures of MetaG1 (Type I) and MetaG9 (Type II) using a gradient from blue (high conservation) to red (low conservation). B. Venn diagram representing the number of positions of the MSAs with rvET scores < 1.5 uniquely found in Type I and Type II enzymes, as well as common conserved positions between both groups. C. rvET scores for regions that have been deemed critical for the activity of PETases: catalytic triad, subsite I, subsite II, extended loop and additional cysteine residue making active site disulfide bonds exclusively in Type II enzymes. The colors of the boxes indicating which residues correspond to each classification match the colors of the residues in stick representation in A.

First, we found that Type I enzymes are overall more conserved – i.e., have more residues with rvET scores close to 1 – than Type II enzymes (Fig. 5A). While this could be due to a higher residue conservation, the imbalance in the number of Type I (11 sequences) and Type II (22 sequences) enzymes analyzed from the high-confidence PETase-like clade could also partly explain these results. When analyzing the residues for the whole clade with rvET scores < 1.5, which empirically correspond to strict conservation or substitution by amino acids with similar physicochemical properties, 51 residues exhibit high sequence conservation across all enzymes (Fig. 5B). When analyzing each type separately, we found greater sequence conservation within the Type I group, with more residues (42 unique to this group) than in Type II enzymes (20 residues unique to this group).

Regions relevant for catalytic activity (residues S160, D206 and H237 in *Is*PETase) exhibit strict sequence conservation (rvET score of 1) in both Type I and Type II enzymes (Fig. 5C). Additionally, Type I enzymes showed increased sequence conservation in subsites I and II, which allow for the accommodation of the aromatic rings of the PET molecule [83] (Fig. 5A), in comparison to Type II enzymes. In contrast, the conservation of the extra cysteine residue pair that forms a disulfide bond in the active site of Type II enzymes [31,83] (DB2 in Fig. 5A) exhibits the same strict conservation (rvET score of 1.0) as the catalytic triad in these enzymes (Fig. 5C). This indicates that all Type II enzymes harbor the cysteines required to form the active site disulfide bond and represents a unique type-determinant sequence signal between these PETase-like enzymes (Fig. 4 and Fig. 5C). These findings highlight the importance of these sequence determinants for the stability and optimal temperature for activity of these enzymes.

### Experimental validation of the polyesterase activity of PETase-like enzymes

To functionally validate our *in silico* predictions, we selected 9 representative metagenomic enzymes from the high-confidence PETase-like clade, encompassing both Type I (MetaG1–MetaG4) and Type II (MetaG5–MetaG9) enzymes (see Fig. 4) based on their M5 motif conservation and phylogenetic positions with the purpose of screening for polyesterase activity. These enzymes were chosen to reflect biogeographic diversity, including both polar-derived (MetaG1, MetaG2, MetaG5, MetaG6, MetaG7, MetaG9) and temperate-derived (MetaG3, MetaG4, MetaG8) sequences.

Out of the 10 enzymes tested –including MetaG4 and MetaG4-CBM –, six (MetaG1, MetaG3, MetaG4, MetaG4-CBM, MetaG6, and MetaG9) exhibited clear PCL-degrading activity (Fig. 6 and Supplementary Fig. S3), confirming their classification as polyesterases. Notably, enzymes from both Type I (typically thermophile-associated) and Type II (typically mesophile/psychrophile-associated) groups demonstrated activity at low (14°C) and moderate (25–37°C) temperatures. These findings are particularly significant in the case of MetaG1, a Type I enzyme from a polar source and more prevalent in polar than temperate environments (Fig. 4), which showed robust and broad activity across a wide thermal range (14–50°C), with optimal degradation at 37°C (Fig. 6 and Supplementary Fig. S3). This suggests that thermophilic-like enzymes can retain catalytic activity in cold-adapted microbial communities, supporting the hypothesis that cold marine environments harbor enzymes with various models of thermal adaptation.

**Fig. 6.**
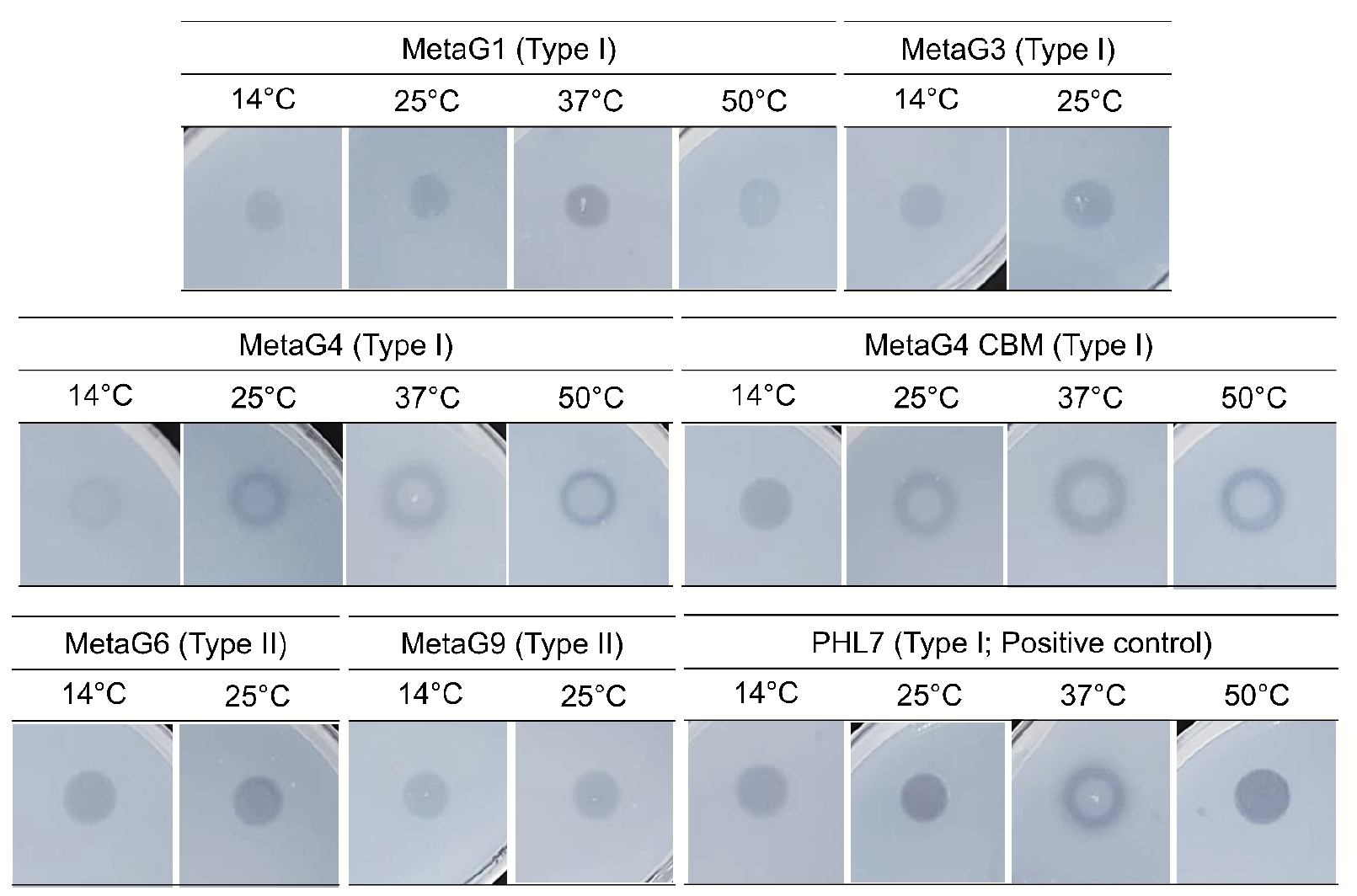
Experimental validation of the polyesterase activity for selected enzymes predicted from metagenomic sequences belonging to the high-confidence PETase-like clade. Degradation screening assays using PCL agar plates with crude extracts of enzymes heterologously expressed in *E. coli* BL21(DE3), which show degradation after 2 days of incubation at different temperatures. Polar-derived enzymes correspond to MetaG1, MetaG2, MetaG5, MetaG6, MetaG7 and MetaG9, and temperate-derived enzymes correspond to MetaG3, MetaG4, MetaG4-CBM, and MetaG8. Some of the plate clearing shows signs of potential protein aggregation after addition of the crude extracts, observed as a white center in the degradation halos, a conclusion drawn from problems with protein aggregation during the protein purification process of enzymes such as MetaG4 (data not shown).

We also tested if the enzymes that showed no activity in the PCL plate degradation assays were due to low protein yields in the host strain for recombinant expression, in this case *E. coli* BL21 (DE3). Testing recombinant protein expression at 18 and 37 °C showed that, with the exception of MetaG2 and MetaG8, most proteins are present in the cell lysate and the cell pellet after induction of protein overexpression with IPTG (Supplementary Fig. S6), thus leaving MetaG5 and MetaG7 as the only enzymes for which PCL plate clearing activity was not observed despite presence in the lysate.

Unfortunately, our results also demonstrate that *E. coli* BL21 (DE3) is not an appropriate host for protein expression and purification of these enzymes for further characterization, which is why we decided to test the expression of the active enzymes from our PCL clearing assay in other *E. coli* strains to pursue the soluble expression of these proteins and their successful purification using His-tagged based immobilized metal affinity chromatography (IMAC) using HisTrap columns followed by size exclusion chromatography (Supplementary Fig. S7).

Of all active enzymes, MetaG9 did not solubly expressed in either *E. coli* Origami 2 DE3 pLysS (OR2) and *E. coli* Rosetta-Gami 2 DE3 pLysS (RG2) (data not shown), MetaG1 quickly aggregated in our buffer conditions after purification using HisTrap columns, and MetaG4 and MetaG4-CBM quickly aggregated after purification using size exclusion chromatography. Thus, only MetaG3 and MetaG6 were successfully purified and amenable for *in vitro* polyester activity assays.

We first determined the optimal temperature for the enzymatic degradation of PCL nanoparticles by MetaG3 and MetaG6 based on the linear initial rates of degradation when measuring the decrease in turbidity of the solution at OD_600_ for 10 min. The reaction was triggered by adding 0.79 mg/mL PCL nanoparticles into a reaction buffer containing either 8 µg/ml of MetaG3 or 4 µg/ml of MetaG6 preincubated at a given temperature between 20-60 °C. Interestingly, the results showed that MetaG3, initially described as temperate-derived, has an optimal temperature of 40 °C, while MetaG6, initially described as polar-derived, has an optimal temperature of 50 °C (Supplementary Fig. S8).

Using these experimentally determined optimal temperatures, we performed enzyme kinetic assays by fixing the concentration of PCL nanoparticles while increasing the enzyme concentration in the enzymatic reactions, and fitting the initial hydrolysis rate data to a pseudo-first-order heterogeneous catalysis as described in equation 1 (see *Methods*). As seen in Fig.7, the results show that MetaG6 is slightly more active at 50 °C than MetaG3 at 40 °C. From the fit of equation 1, we determined the hydrolysis rate constant (*k_τ_*) and the adsorption equilibrium constant (K_A_) of both enzymes (Table 2), showing that MetaG6 has 1.7-fold higher *k_τ_* and half the K_A_ than MetaG3.

**Fig. 7.**
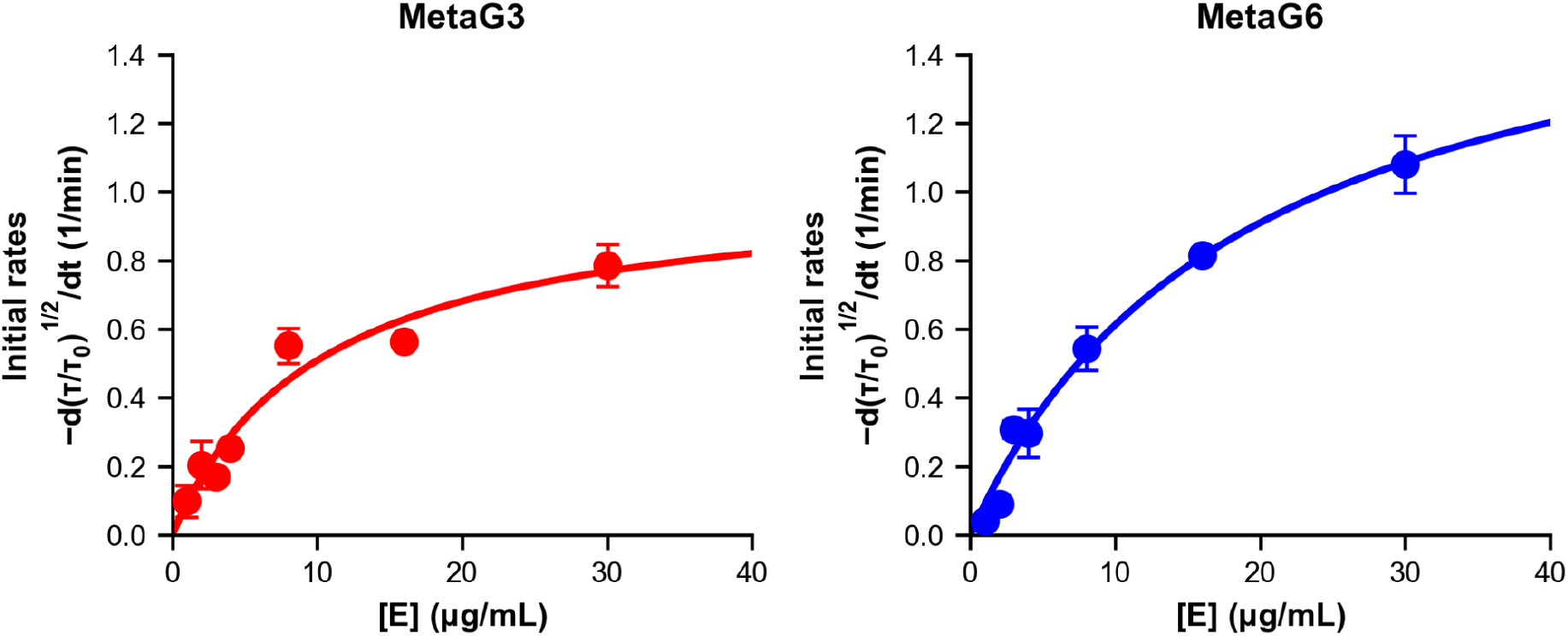
Enzyme kinetics of PCL nanoparticle degradation by MetaG3 and MetaG6. PCL nanoparticle degradation assays were performed at a fixed concentration of 0.79 mg/mL PCL nanoparticles and increasing the enzyme concentration between 1 and 30 µg/mL. Experiments were performed in three technical replicates.

**Table 2.** Kinetic parameters of MetaG3 and MetaG6 for hydrolysis of PCL nanoparticles. Enzymatic activities determined by varying enzyme concentration from 1 to 30 µg/mL. MetaG3 was assayed at 40 °C, whereas MetaG6 was assayed at 50 °C. Results are the mean ± standard deviation derived from the fit performed in technical triplicate.

| Enzyme | *k<sub>τ</sub>* (10<sup>-3</sup>/min<sup>-1</sup>) | K<sub>A</sub> (mg/ml) |
|---|---|---|
| MetaG3 | 1031 ± 146 | 97 ± 31 |
| MetaG6 | 1769 ± 211 | 53 ± 12 |

### Evidence for complete PET biodegradation potential in polar microbial communities

We evaluated the potential for complete PET degradation by extending our analysis to enzymes acting downstream of PET hydrolysis intermediates, namely the extracellular degradation of MHET [17] and the intracellular conversion of TPA into protocatechuate (PCA) for its use as carbon source via the PCA 4,5-cleavage pathway [112]. In *I. sakaiensis* 201-F6, these steps are carried out by an extracellular MHETase [18,84], and by an intracellular TPA-degrading complex, specifically homologs of TphA1, TphA2, TphA3 and TphB [17,113], respectively. Using HMM-based screening on polar and temperate metagenomic sequences, we identified 209 putative MHETase-like sequences and 442 sequences associated with the TPA-degrading complex (Supplementary Fig. S4 and Supplementary Fig. S5).

In general, we found that key catalytic residues of MHETase, including the canonical catalytic triad (S225, D492, H528) [84], the oxyanion hole (G132) [18], and a critical disulfide bond (C224-C529) [83,84], were largely conserved among the putative MHETases (Supplementary Fig. S4). Additional residues known to enhance enzymatic activity in *Is*MHETase and experimentally described homologs from *Comamonas thiooxydans* and *Hydrogenophaga* sp. PML113, such as S131, D226, R411, F415, F424 and F495 [18,114], were less conserved. This variability suggests that while a subset of the newly identified sequences may retain functional MHETase activity, others could exhibit reduced efficiency or broadened substrate specificity, potentially acting on structurally related compounds. This hypothesis requires further experimental validation through biochemical and structural characterization.

We also quantified the abundance and distribution of MHETase-like enzymes and TPA degradation complex enzymes across polar and temperate oceanic regions. Our results revealed that these enzymes are widespread throughout the global ocean but show a notable enrichment in polar environments, mirroring the pattern observed for PETase-like enzymes (Supplementary Fig. S5, Supplementary Table S7). In particular, the higher abundance of putative MHETases and TphA1 proteins in Arctic waters suggests possible regional specialization in the degradation process.

To further contextualize these functional genes within microbial community structure, we used publicly available, and also reconstructed metagenome-assembled genomes (MAGs) from polar and temperate metagenomes, and detected 112 MAGs containing at least one enzyme from the PET degradation pathway (PETase, MHETase, TphA1, TphA2, TphA3 or ThpB) (59 from temperate and 53 from polar metagenomes).

These MAGs spanned 10 distinct bacterial classes (Fig. 8), with Gammaproteobacteria (n=51), followed by Bacteroidia (n=27), Alphaproteobacteria (n=10), and Verrucomicrobiae (n=8) being the most represented. This taxonomic distribution is consistent with previous studies reporting a prevalence of plastic-degrading enzymes in these marine bacteria, particularly within Gammaproteobacteria and Bacteroidetes [25,26,115]. However, our findings expand this repertoire to Verrucomicrobiae and Alphaproteobacteria, taxa that may be underexplored contributors to PET degradation in marine systems, although their PET-degrading activity remains to be experimentally validated.

**Fig. 8.**
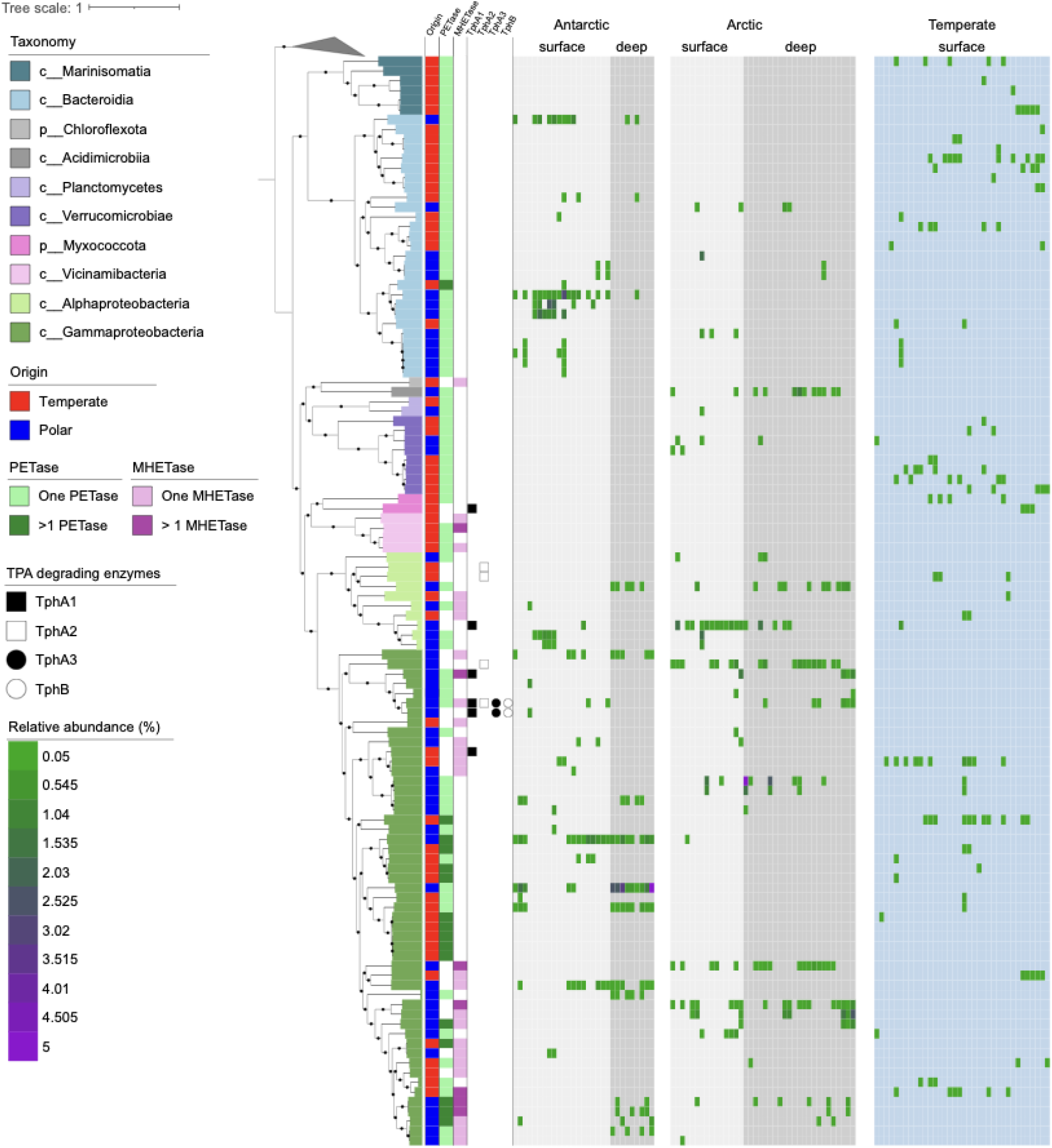
Taxonomic affiliations and abundance of marine metagenome-assembled genomes (MAGs) containing at least one PET degradation pathway enzyme. The phylogenetic reconstruction was performed using 120 concatenated single-copy bacterial genes, and 10 Aquificota MAGs were included as an outgroup.

Among the 112 MAGs, 89 encoded putative PETase genes, 31 carried putative MHETase genes, and 9 contained at least one gene from the TPA degradation complex. Sixteen MAGs encoded two or more PETase-like genes, with the highest number (seven) found in a polar MAG (Genome_085). Moreover, 15 MAGs encoded multiple putative PET-degrading enzymes, including 13 with both PETase-like and MHETase-like homologs, suggesting a metabolic potential for more complete or efficient PET biodegradation pathways (Supplementary Table S8). Comparable multi-enzyme genomic contexts have been described in temperate marine bacteria [25,26], but their recovery from polar datasets underscores once again the ecological significance of cold-adapted microbial communities. Notably, nine of the 15 multi-enzyme MAGs recovered in the present study, originated from polar datasets. Among them, Genome_042 (98.6% completeness; 0.4% contamination) was the only MAG containing candidates for PET hydrolysis, MHET hydrolysis, and TPA conversion. In this MAG, the putative PET- and MHET-hydrolase genes were recovered on separate contigs, while the putative TphA2, TphA3, and TphA1 genes are consecutive on one contig and putative TphB farther downstream. An IclR-family regulator and an IS5-family transposase occur upstream of this Tph region (Supplementary Table S9). Given the fragmented nature of the assembly, whether the PETase and MHETase genes are physically close to one another and to Tph genes remains to be determined.

## DISCUSSION

Plastic pollution is becoming increasingly pervasive in marine environments, yet the microbial potential for plastic biodegradation, particularly in cold ecosystems, remains poorly understood [116,117]. Recent efforts, including the global survey by Alam et al. [25], have expanded our knowledge of PET-degrading enzymes in temperate and tropical marine systems, but overlooked polar environments. To address this gap, we systematically identified putative PETase-like enzymes from a comprehensive set of metagenomic samples spanning Arctic and Antarctic marine environments.

By extending the biogeographical scope to polar marine ecosystems, our study uncovered PETase-like candidates from cold-adapted microbial communities in both surface (0-100 m) and deep (>100 m) Arctic and Antarctic waters. This contribution is ecologically significant, as polar regions, despite being increasingly exposed to plastic pollution [30], have been subject to limited sampling and remain understudied with regards to their plastic biodegradation potential in comparison to temperate and tropical regions. This is especially relevant in the context of accelerated polar warming and changing ocean circulation, which are expected to influence both plastic distribution and the ecological role of microbial communities in mitigating its impact.

Although by the time of review of this work, an article identifying putative PETases from Antarctic environments was published [118], it only examined potential PETases across Antarctic terrestrial ecosystems and based its conclusions on molecular docking and molecular dynamics, with no experimental validation of the polyesterase activity of these enzymes nor expanding towards finding putative MHETases and TPA-metabolizing proteins.

Because our HMM approach was built from nine experimentally validated PETases, divergent functional lineages may have been missed in this study. Thus, despite additional motif-based and phylogenetic analyses, our findings should be viewed as a set of PETase-like candidates rather than an exhaustive inventory of PET-degrading enzymes in polar waters. The same caveat applies to the PET-degradation pathway enzymes discussed below, which were identified by homology to a small set of characterized and experimentally validated enzymes.

Beyond expanding the known geographic distribution of these PETase-like candidates, our findings raise questions about how different motif groups are distributed across marine environments. The apparent greater representation of M4-, M4 strict-, and M5-containing sequences in polar waters, may point to ecological differences worth investigating. This pattern was especially pronounced for M5 sequences in deep Antarctic metagenomes. Although cold temperatures and high hydrostatic pressures may select for these potentially efficient polyesterases, the divergence in M5 motif abundance between deep Arctic and Antarctic metagenomes indicates that other factors must also drive this distribution including their fundamentally different oceanographic histories and community assembly processes [119,120]. However, these patterns should be interpreted with caution, because polar and temperate metagenomes were generated by independent studies and differences in sampling and sequencing protocols may also contribute to the RPKM-based patterns despite normalization. Cross-regional sampling and experimental characterization will be needed to evaluate their ecological and functional significance.

The presence of advanced motifs (i.e., M5) in polar samples (Fig. 3), and further validation of the polyesterase activity of selected candidates with this motifs from polar marine environments (Fig. 6), raises the question of whether PETase-like enzymes may have evolved as a selective advantage in response to increasing plastic pollution [30], enabling microorganisms to access an additional carbon and energy source. However, the evidence for direct coupling between plastic pollution and the abundance and activity of plastic-degrading enzymes remains mixed. Studies employing global metagenomic analyses have suggested that there is a correlation between the abundance of potential plastic-degrading enzymes and the degree of plastic pollution in sampled natural environments, implying that the nearly 70 years of plastic contamination has given ample time and selective pressure to evolutionarily repurpose enzymes for plastic degradation [115]. Conversely, other studies have found no significant correlation between levels of plastic and the expression of these enzymes in marine environments [121].

A simple pollution-driven explanation for this polar enrichment seems unlikely, as the Southern Ocean has been shown to yield the third-highest number of potential plastic-degrading enzyme hits despite being the least polluted ocean analyzed in previous studies [121]. An alternative proposition is that the PET molecule shares structural similarities with naturally occurring polymers that are abundant in polar environments. This possibility is drawn from the fact that the most efficient PET-degrading natural enzymes to date, LCC [52] and PHL7 [104], were both found in plant compost metagenomes and are classified as cutinases, i.e., they degrade the plant polymer cutin. Similarly, PETase-like enzymes in these polar regions may show polyesterase activity as well as hydrolytic activity against their native substrates due to enzyme promiscuity [122]. In contrast, the high abundance and widespread presence of M0 motif-containing sequences (Fig. 3) across all biogeographic regions is consistent with the ubiquity of the catalytic triad within the α/β hydrolase fold superfamily [123], and likely represents canonical esterases with limited or no PET-degrading capacity.

Our functional validation (Fig. 6 and Fig. 7) also showed that the Type I/Type II distinction, although phylogenetically well-resolved and supported by the strict conservation of the active site disulfide bond in Type II enzymes, does not translate directly into optimal temperature for activity. The two polar-derived enzymes purified and characterized against PCL nanoparticles, MetaG3 and MetaG6, spanned a broader thermal range than initially expected based on the PCL plate clearing assays (Fig. 6). MetaG3, classified as a Type I enzyme, showed activity up to 25 °C in the PCL plate clearing assays (Supplementary Fig. S3) while exhibiting an optimal temperature for activity in *in vitro* PCL nanoparticle degrading assays of 40 °C (Fig. 7). Conversely, the Type II polyesterase MetaG6 showed activity up to 37 °C in PCL plate clearing assays (Supplementary Fig. S3), yet an optimum of 50 °C against PCL nanoparticles *in vitro* (Fig. 7). This is not necessarily contradictory, as the extended loop and additional disulfide bond of Type II enzymes confer active site flexibility and stability tradeoffs, a property compatible with activity over a broad thermal range rather than with a strictly cold-shifted optimum. Rather than being restricted to low temperatures, polar polyesterases appear to combine cold activity with thermal breadth, a combination of particular interest for bioremediation under variable conditions.

Beyond PET-degrading enzymes, our work further extends on previous studies that have reported MHETases and TPA degradation genes in temperate marine systems [25,26,115], often suggesting that efficient plastic degradation may be more characteristic of warmer waters. By analyzing metagenomes from polar marine environments, our findings demonstrate that the downstream components of the PET degradation pathway are not only present (Fig. 8) but relatively enriched in cold polar microbiomes (Supplementary Fig. S5).

Notably, although PETase- and MHETase-like genes were broadly distributed among the analyzed MAGs, TPA-degradation genes were detected in relatively few genomes. This pattern may reflect cross-feeding, in which some taxa perform extracellular depolymerization while others import and assimilate aromatic intermediates such as TPA [124]. Thus, organisms encoding PETases or MHETases do not need to carry the downstream pathway for microbial community degradation of plastic pollutants. Alternatively, the pattern observed in our results may partly reflect methodological limitations. Aromatic-compound degradation genes may occur in mobile elements, whose tetranucleotide frequencies and coverage can differ from those of the host chromosome, making them more likely to be excluded from MAGs during binning. Additionally, the high overall completeness of the analyzed MAGs (median 89.2%) does not preclude that Tph genes were not detected because they were not assembled and/or binned. Therefore, the low recovery of TPA-degradation genes at the MAG level should not be taken as evidence that this step is absent from these communities.

Nevertheless, a polar MAG was found to harbor the complete set of putative PET, MHET, and TPA-degrading enzymes required for using such polyester as a carbon source, highlighting the potential for autonomous plastic degradation in polar taxa. In this MAG, the local clustering of the TPA-conversion candidates and nearby transcriptional regulators is consistent with a potentially coordinated downstream module, while the adjacent IS5-family transposase raises the possibility that genomic rearrangement or horizontal acquisition has contributed to its organization. The genomic relationship of the PET- and MHET-hydrolase candidates between them and to the Tph module in this MAG remains unresolved as they occur on separate assembled contigs; future long-read sequencing could help link these contigs and clarify the genomic organization of the candidate pathway. To our knowledge, such a complete genetic repertoire for PET biodegradation has not been previously reported from polar environments, as earlier global surveys [25,115] lacked evidence of fully reconstructed downstream pathways in cold ecosystems.

Altogether, these findings position polar environments as critical yet overlooked reservoirs of plastic biodegradation potential in high-latitude oceans. It further suggests that cold-adapted enzymes and microbial consortia could play an underestimated role in plastic degradation in high-latitude oceans, particularly under scenarios of increasing anthropogenic plastic inputs and rapid climate-driven environmental change.

## CONCLUSIONS

This study demonstrates that polar marine ecosystems represent previously overlooked reservoirs of enzymes with the potential to degrade PET and other polyesters under significantly lower temperatures than most plastic-degrading enzymes described so far [15], expanding from the previously described PET-degrading enzymes from *Moraxella sp.* TA144 and *Oleispira antarctica* RB-8 with optimum temperatures for enzymatic activity near 25°C [31].

By combining genomic, phylogenetic, bioinformatic and functional analyses, we reveal novel aspects of the taxonomic and genetic diversity underpinning plastic biodegradation in cold oceanic environments. On the one hand, our study is – to the best of our knowledge – the first to experimentally validate the polyesterase activity of PETase-like enzymes directly recovered from polar marine metagenomes, demonstrating their functional capacity at low temperatures and expanding the known thermal and ecological breadth of marine polyesterases. On the other hand, the marked enrichment of putative PET-degrading enzymatic machinery (particularly PETases, MHETases, and TPA-degrading enzymes) in polar marine metagenomes highlights both the metabolic versatility and ecological resilience of microbial communities exposed to increasing anthropogenic plastic pollution.

In contrast to earlier surveys that focused on temperate ecosystems, our findings extend the global biogeography of plastic-degrading enzymes to high-latitude oceans, where cold-adapted microbial consortia may play a more important role in plastic turnover than previously assumed, underscoring the evolutionary plasticity of these enzymes to operate across diverse temperature regimes.

Beyond advancing ecological understanding, this work also identifies polar microbiomes as a promising source of cold-active biocatalysts, with direct implications for the design of energy-efficient biotechnological applications and the development of sustainable, low-temperature plastic bioremediation strategies in both natural and engineered environments.

## ETHICS APPROVAL AND CONSENT TO PARTICIPATE

Not applicable

## CONSENT FOR PUBLICATION

Not applicable

## AVAILABILITY OF DATA AND MATERIAL

All metagenomic data used in this study is publicly available under BioProjects PRJEB7988 (OM-RGC.v1), PRJNA588686 (PM-RGC), PRJNA805531 and PRJNA1280935 (Chile Bay, Western Antarctic Peninsula). The OM-RGC.v1 catalog is also available at https://tara-oceans.mio.osupytheas.fr/ocean-gene-atlas/). The amino acid sequences of the putative PETases, MHETases and TPA-degrading enzymes, the sequences of the full-length and truncated enzymes MetaG1-MetaG9 after bioinformatic analysis of the presence of signal peptides and disordered regions, the ColabFold-predicted structures of MetaG1-MetaG9, the rvET analysis of the high-confidence PETase-like clade, and the 112 MAGs with identified putative PETases, MHETases and/or TPA-degrading enzymes are available on Zenodo (https://dx.doi.org/10.5281/zenodo.18306301).

## COMPETING INTERESTS

The authors declare that they have no competing interests.

## FUNDING

This research was funded by the National Agency for Research and Development (ANID) from Chile and the Sao Paulo State Foundation for Research (FAPESP) from Brazil through their International Cooperation Program (ANID-FAPESP PCI 2019/13259-9), the International Centre for Genetic Engineering and Biotechnology (ICGEB) through its Collaborative Research Program (CRP/CHL23-02), the Chilean Antarctic Institute (INACH RG_47-16 and RT_04-19), the ANID Millennium Science Initiative Program (ICN17_022 and ICN2021_044), the ANID Fund for Research Centers in Prioritary Areas (FONDAP 1523A0002) and the Homeworld Collective Garden Grants. Aransa Griñen was supported by an ANID doctoral scholarship (PFCHA 21220450). Powered@NLHPC: This research was partially supported by the supercomputing infrastructure of the NLHPC (CCSS210001).

## AUTHORS’ CONTRIBUTIONS

AG, JC-A, MB, PV-B, ANV and FE analyzed and interpreted the metagenomics data to identify putative PETases, MHETases and TPA-degrading enzymes. AG and CAR-S performed the protein structure predictions of the selected MetaG enzymes. AG, ANV, RP-L, DW-A and FGV performed the recombinant expression and experimental validation of the enzymes studied in this work. JCA, MB and PV-B performed the reconstruction of the MAGs and the analysis of the enzymes found in these genomes. CAR-S and BD co-supervised this work. All authors participated in writing the manuscript. All authors read and approved the final manuscript.

## ACKNOWLEDGEMENTS

We are grateful to the personnel of Armada de Chile and Instituto Antártico Chileno (INACH) at Arturo Prat station for making our work in Antarctica possible, and also to María Estrella Alcamán and Javier Tamayo-Leiva for their technical assistance during sample collection.

## Notes

https://doi.org/10.5281/zenodo.18306300

### Competing Interest Statement

The authors have declared no competing interest.

### Summary of Updates

Three authors added; additional methods and results of experiments related to protein purification and characterization for enzyme kinetics; improved discussion and conclusions for clarity; supplemental files updated

## REFERENCES

1. OECD. Policy scenarios for eliminating plastic pollution by 2040. OECD; 2024.

2. PlasticsEurope, 2024. Plastics – the fast Facts 2024 [Internet]. {Plastics Europe, 2024}. https://plasticseurope.org/es/knowledge-hub/plastics-the-fast-facts-2024/

3. Cottom JW, Cook E, Velis CA. A local-to-global emissions inventory of macroplastic pollution. Nature. Springer Science and Business Media LLC; 2024;633:101–8.

4. Geyer R, Jambeck JR, Law KL. Production, use, and fate of all plastics ever made. Sci Adv. 2017;3:e1700782.

5. Jambeck JR, Geyer R, Wilcox C, Siegler TR, Perryman M, Andrady A, et al. Marine pollution. Plastic waste inputs from land into the ocean. Science. American Association for the Advancement of Science (AAAS); 2015;347:768–71.

6. OECD. Global plastics outlook: economic drivers, environmental impacts and policy options. Paris Cedex, France: Organization for Economic Co-operation and Development (OECD); 2022.

7. PlasticsEurope, 2022. Plastics—The Facts 2022 [Internet]. {PlasticsEurope, 2022}. https://plasticseurope.org/knowledge-hub/plastics-the-facts-2022/

8. Oliveira J, Belchior A, da Silva VD, Rotter A, Petrovski Ž, Almeida PL, et al. Marine environmental plastic pollution: Mitigation by microorganism degradation and recycling valorization. Front Mar Sci [Internet]. Frontiers Media SA; 2020;7. 10.3389/fmars.2020.567126

9. Taniguchi I, Yoshida S, Hiraga K, Miyamoto K, Kimura Y, Oda K. Biodegradation of PET: Current status and application aspects. ACS Catal. American Chemical Society (ACS); 2019;9:4089–105.

10. Garcia JM, Robertson ML. The future of plastics recycling. Science. American Association for the Advancement of Science (AAAS); 2017;358:870–2.

11. Rahimi A, García JM. Chemical recycling of waste plastics for new materials production. Nat Rev Chem. Springer Science and Business Media LLC; 2017;1:0046.

12. Schyns ZOG, Shaver MP. Mechanical recycling of packaging plastics: A review. Macromol Rapid Commun. Wiley; 2021;42:e2000415.

13. García JM. Catalyst: Design challenges for the future of plastics recycling. Chem. Elsevier BV; 2016;1:813–5.

14. Kawai F, Kawabata T, Oda M. Current knowledge on enzymatic PET degradation and its possible application to waste stream management and other fields. Appl Microbiol Biotechnol. Springer Science and Business Media LLC; 2019;103:4253–68.

15. Wei R, Zimmermann W. Biocatalysis as a green route for recycling the recalcitrant plastic polyethylene terephthalate. Microb Biotechnol. John Wiley & Sons, Ltd; 2017;10:1302–7.

16. Wei R, Zimmermann W. Microbial enzymes for the recycling of recalcitrant petroleum-based plastics: how far are we? Microb Biotechnol. 2017;10:1308–22.

17. Yoshida S, Hiraga K, Takehana T, Taniguchi I, Yamaji H, Maeda Y, et al. A bacterium that degrades and assimilates poly(ethylene terephthalate). Science. American Association for the Advancement of Science (AAAS); 2016;351:1196–9.

18. Knott BC, Erickson E, Allen MD, Gado JE, Graham R, Kearns FL, et al. Characterization and engineering of a two-enzyme system for plastics depolymerization. Proc Natl Acad Sci U S A. Proceedings of the National Academy of Sciences; 2020;117:25476–85.

19. Sadler JC, Wallace S. Microbial synthesis of vanillin from waste poly(ethylene terephthalate). Green Chem. 2021;23:4665–72.

20. Schneier A, Melaugh G, Sadler JC. Engineered plastic-associated bacteria for biodegradation and bioremediation. Biotechnol Environ. 2024;1:7.

21. Diao J, Hu Y, Tian Y, Carr R, Moon TS. Upcycling of poly(ethylene terephthalate) to produce high-value bio-products. Cell Rep. 2023;42:111908.

22. Bollinger A, Thies S, Knieps-Grünhagen E, Gertzen C, Kobus S, Höppner A, et al. A novel polyester hydrolase from the marine bacterium Pseudomonas aestusnigri - structural and functional insights. Front Microbiol. Frontiers Media SA; 2020;11:114.

23. Li Z, Zhao Y, Wu P, Wang H, Li Q, Gao J, et al. Structural insight and engineering of a plastic degrading hydrolase Ple629. Biochem Biophys Res Commun. 2022;626:100–6.

24. Meyer Cifuentes IE, Wu P, Zhao Y, Liu W, Neumann-Schaal M, Pfaff L, et al. Molecular and Biochemical Differences of the Tandem and Cold-Adapted PET Hydrolases Ple628 and Ple629, Isolated From a Marine Microbial Consortium. Front Bioeng Biotechnol. 2022;10:930140.

25. Alam I, Marasco R, Momin AA, Aalismail N, Laiolo E, Martin C, et al. Widespread distribution of bacteria containing PETases with a functional motif across global oceans. ISME J [Internet]. 2025;19. 10.1093/ismejo/wraf121

26. Danso D, Schmeisser C, Chow J, Zimmermann W, Wei R, Leggewie C, et al. New insights into the function and global distribution of Polyethylene terephthalate (PET)-degrading bacteria and enzymes in marine and terrestrial metagenomes. Appl Environ Microbiol [Internet]. 2018;84. 10.1128/AEM.02773-17

27. Chen J, Jia Y, Sun Y, Liu K, Zhou C, Liu C, et al. Global marine microbial diversity and its potential in bioprospecting. Nature. Springer Science and Business Media LLC; 2024;633:371–9.

28. Leistenschneider C, Burkhardt-Holm P, Mani T, Primpke S, Taubner H, Gerdts G. Microplastics in the Weddell Sea (Antarctica): A forensic approach for discrimination between environmental and vessel-induced microplastics. Environ Sci Technol. American Chemical Society (ACS); 2021;55:15900–11.

29. Lacerda ALDF, Rodrigues LDS, van Sebille E, Rodrigues FL, Ribeiro L, Secchi ER, et al. Plastics in sea surface waters around the Antarctic Peninsula. Sci Rep. Springer Science and Business Media LLC; 2019;9:3977.

30. Waller CL, Griffiths HJ, Waluda CM, Thorpe SE, Loaiza I, Moreno B, et al. Microplastics in the Antarctic marine system: An emerging area of research. Sci Total Environ. Elsevier BV; 2017;598:220–7.

31. Blázquez-Sánchez P, Engelberger F, Cifuentes-Anticevic J, Sonnendecker C, Griñén A, Reyes J, et al. Antarctic Polyester Hydrolases Degrade Aliphatic and Aromatic Polyesters at Moderate Temperatures. Appl Environ Microbiol. 2022;88:e0184221.

32. Feller G, Gerday C. Psychrophilic enzymes: hot topics in cold adaptation. Nat Rev Microbiol. Springer Science and Business Media LLC; 2003;1:200–8.

33. D’Amico S, Collins T, Marx J-C, Feller G, Gerday C. Psychrophilic microorganisms: challenges for life. EMBO Rep. EMBO; 2006;7:385–9.

34. Fecker T, Galaz-Davison P, Engelberger F, Narui Y, Sotomayor M, Parra LP, et al. Active site flexibility as a hallmark for efficient PET degradation by I. sakaiensis PETase. Biophys J. Elsevier BV; 2018;114:1302–12.

35. Cavicchioli R, Charlton T, Ertan H, Mohd Omar S, Siddiqui KS, Williams TJ. Biotechnological uses of enzymes from psychrophiles. Microb Biotechnol. Wiley; 2011;4:449–60.

36. Santiago M, Ramírez-Sarmiento CA, Zamora RA, Parra LP. Discovery, molecular mechanisms, and industrial applications of cold-active enzymes. Front Microbiol. Frontiers Media SA; 2016;7:1408.

37. Obbard RW, Sadri S, Wong YQ, Khitun AA, Baker I, Thompson RC. Global warming releases microplastic legacy frozen in Arctic Sea ice. Earths Future. American Geophysical Union (AGU); 2014;2:315–20.

38. Bergmann M, Wirzberger V, Krumpen T, Lorenz C, Primpke S, Tekman MB, et al. High quantities of microplastic in arctic deep-sea sediments from the HAUSGARTEN observatory. Environ Sci Technol. American Chemical Society (ACS); 2017;51:11000–10.

39. González-Pleiter M, Edo C, Velázquez D, Casero-Chamorro MC, Leganés F, Quesada A, et al. First detection of microplastics in the freshwater of an Antarctic Specially Protected Area. Mar Pollut Bull. Elsevier BV; 2020;161:111811.

40. Marx J-C, Collins T, D’Amico S, Feller G, Gerday C. Cold-adapted enzymes from marine Antarctic microorganisms. Mar Biotechnol (NY). Springer Science and Business Media LLC; 2007;9:293–304.

41. Collins T, Feller G. Psychrophilic enzymes: strategies for cold-adaptation. Essays Biochem. Portland Press Ltd.; 2023;67:701–13.

42. Eddy SR. Accelerated Profile HMM Searches. PLoS Comput Biol. 2011;7:e1002195.

43. Austin HP, Allen MD, Donohoe BS, Rorrer NA, Kearns FL, Silveira RL, et al. Characterization and engineering of a plastic-degrading aromatic polyesterase. Proc Natl Acad Sci U S A. 2018;115:E4350–7.

44. Ribitsch D, Acero EH, Greimel K, Eiteljoerg I, Trotscha E, Freddi G, et al. Characterization of a new cutinase from*Thermobifida alba*for PET-surface hydrolysis [Internet]. Biocatalysis and Biotransformation. 2012. p. 2–9. 10.3109/10242422.2012.644435

45. Kitadokoro K, Kakara M, Matsui S, Osokoshi R, Thumarat U, Kawai F, et al. Structural insights into the unique polylactate-degrading mechanism of Thermobifida alba cutinase. FEBS J. Wiley; 2019;286:2087–98.

46. Acero EH, Ribitsch D, Steinkellner G, Gruber K, Greimel K, Eiteljoerg I, et al. Enzymatic Surface Hydrolysis of PET: Effect of Structural Diversity on Kinetic Properties of Cutinases from Thermobifida [Internet]. Macromolecules. 2011. p. 4632–40. 10.1021/ma200949p

47. Roth C, Wei R, Oeser T, Then J, Föllner C, Zimmermann W, et al. Structural and functional studies on a thermostable polyethylene terephthalate degrading hydrolase from Thermobifida fusca. Appl Microbiol Biotechnol. Springer Science and Business Media LLC; 2014;98:7815–23.

48. Ribitsch D, Hromic A, Zitzenbacher S, Zartl B, Gamerith C, Pellis A, et al. Small cause, large effect: Structural characterization of cutinases from Thermobifida cellulosilytica. Biotechnol Bioeng. 2017;114:2481–8.

49. Ribitsch D, Acero EH, Greimel K, Dellacher A, Zitzenbacher S, Marold A, et al. A New Esterase from Thermobifida halotolerans Hydrolyses Polyethylene Terephthalate (PET) and Polylactic Acid (PLA) [Internet]. Polymers. 2012. p. 617–29. 10.3390/polym4010617

50. Kawai F, Oda M, Tamashiro T, Waku T, Tanaka N, Yamamoto M, et al. A novel Ca2 -activated, thermostabilized polyesterase capable of hydrolyzing polyethylene terephthalate from Saccharomonospora viridis AHK190 [Internet]. Applied Microbiology and Biotechnology. 2014. p. 10053–64. 10.1007/s00253-014-5860-y

51. Wei R, Oeser T, Then J, Kühn N, Barth M, Schmidt J, et al. Functional characterization and structural modeling of synthetic polyester-degrading hydrolases from Thermomonospora curvata. AMB Express. Springer Science and Business Media LLC; 2014;4:44.

52. Sulaiman S, Yamato S, Kanaya E, Kim J-J, Koga Y, Takano K, et al. Isolation of a novel cutinase homolog with polyethylene terephthalate-degrading activity from leaf-branch compost by using a metagenomic approach. Appl Environ Microbiol. 2012;78:1556–62.

53. Sulaiman S, You D-J, Kanaya E, Koga Y, Kanaya S. Crystal structure and thermodynamic and kinetic stability of metagenome-derived LC-cutinase. Biochemistry. American Chemical Society (ACS); 2014;53:1858–69.

54. Sunagawa S, Coelho LP, Chaffron S, Kultima JR, Labadie K, Salazar G, et al. Ocean plankton. Structure and function of the global ocean microbiome. Science. 2015;348:1261359.

55. Villar E, Vannier T, Vernette C, Lescot M, Cuenca M, Alexandre A, et al. The Ocean Gene Atlas: exploring the biogeography of plankton genes online. Nucleic Acids Res. 2018;46:W289–95.

56. Cao S, Zhang W, Ding W, Wang M, Fan S, Yang B, et al. Structure and function of the Arctic and Antarctic marine microbiota as revealed by metagenomics. Microbiome. 2020;8:47.

57. Alcamán-Arias ME, Cifuentes-Anticevic J, Díez B, Testa G, Troncoso M, Bello E, et al. Surface ammonia-oxidizer abundance during the late summer in the west Antarctic coastal system. Front Microbiol. Frontiers Media SA; 2022;13:821902.

58. Buscaglia M, Iriarte JL, Schulz F, Díez B. Adaptation strategies of giant viruses to low-temperature marine ecosystems. ISME J [Internet]. Oxford University Press (OUP); 2024;18. 10.1093/ismejo/wrae162

59. Fu L, Niu B, Zhu Z, Wu S, Li W. CD-HIT: accelerated for clustering the next-generation sequencing data. Bioinformatics. 2012;28:3150–2.

60. Mahram A, Herbordt MC. NCBI BLASTP on High-Performance Reconfigurable Computing Systems [Internet]. ACM Transactions on Reconfigurable Technology and Systems. 2015. p. 1–20. 10.1145/2629691

61. Parks DH, Chuvochina M, Rinke C, Mussig AJ, Chaumeil P-A, Hugenholtz P. GTDB: an ongoing census of bacterial and archaeal diversity through a phylogenetically consistent, rank normalized and complete genome-based taxonomy. Nucleic Acids Res. Oxford University Press (OUP); 2022;50:D785–94.

62. Buchfink B, Reuter K, Drost H-G. Sensitive protein alignments at tree-of-life scale using DIAMOND. Nat Methods. Springer Science and Business Media LLC; 2021;18:366–8.

63. Zaragoza-Solas A, Baltar F. Ayu: a machine intelligence tool for identification of extracellular proteins in the marine secretome. Nat Commun. Springer Science and Business Media LLC; 2025;16:2793.

64. Edgar RC. Muscle5: High-accuracy alignment ensembles enable unbiased assessments of sequence homology and phylogeny. Nat Commun. Springer Science and Business Media LLC; 2022;13:6968.

65. Minh BQ, Schmidt HA, Chernomor O, Schrempf D, Woodhams MD, von Haeseler A, et al. IQ-TREE 2: New models and efficient methods for phylogenetic inference in the genomic era. Mol Biol Evol. Oxford University Press (OUP); 2020;37:1530–4.

66. Kalyaanamoorthy S, Minh BQ, Wong TKF, von Haeseler A, Jermiin LS. ModelFinder: fast model selection for accurate phylogenetic estimates. Nat Methods. Springer Science and Business Media LLC; 2017;14:587–9.

67. Minh BQ, Nguyen MAT, von Haeseler A. Ultrafast Approximation for Phylogenetic Bootstrap [Internet]. Molecular Biology and Evolution. 2013. p. 1188–95. 10.1093/molbev/mst024

68. Hoang DT, Chernomor O, von Haeseler A, Minh BQ, Vinh LS. UFBoot2: Improving the Ultrafast Bootstrap Approximation. Mol Biol Evol. 2018;35:518–22.

69. Bianchini G, Sánchez-Baracaldo P. TreeViewer: Flexible, modular software to visualise and manipulate phylogenetic trees. Ecol Evol. Wiley; 2024;14:e10873.

70. Almagro Armenteros JJ, Tsirigos KD, Sønderby CK, Petersen TN, Winther O, Brunak S, et al. SignalP 5.0 improves signal peptide predictions using deep neural networks. Nat Biotechnol. Springer Science and Business Media LLC; 2019;37:420–3.

71. Ishida T, Kinoshita K. PrDOS: prediction of disordered protein regions from amino acid sequence. Nucleic Acids Res. Oxford University Press (OUP); 2007;35:W460–4.

72. Mirdita M, Schütze K, Moriwaki Y, Heo L, Ovchinnikov S, Steinegger M. ColabFold: making protein folding accessible to all. Nat Methods. 2022;19:679–82.

73. Jumper J, Evans R, Pritzel A, Green T, Figurnov M, Ronneberger O, et al. Highly accurate protein structure prediction with AlphaFold. Nature. 2021;596:583–9.

74. Steinegger M, Söding J. MMseqs2 enables sensitive protein sequence searching for the analysis of massive data sets. Nat Biotechnol. 2017;35:1026–8.

75. Suzek BE, Wang Y, Huang H, McGarvey PB, Wu CH, UniProt Consortium. UniRef clusters: a comprehensive and scalable alternative for improving sequence similarity searches. Bioinformatics. Oxford University Press (OUP); 2015;31:926–32.

76. Schrödinger, LLC. The PyMOL Molecular Graphics System, Version 3.1.0. 2025.

77. Mihalek I, Res I, Lichtarge O. A family of evolution-entropy hybrid methods for ranking protein residues by importance. J Mol Biol. 2004;336:1265–82.

78. Mihalek I, Res I, Lichtarge O. Evolutionary trace report_maker: a new type of service for comparative analysis of proteins. Bioinformatics. Oxford University Press (OUP); 2006;22:1656–7.

79. Sievers F, Wilm A, Dineen D, Gibson TJ, Karplus K, Li W, et al. Fast, scalable generation of high-quality protein multiple sequence alignments using Clustal Omega. Mol Syst Biol. Springer Science and Business Media LLC; 2011;7:539.

80. Russell RB, Barton GJ. Multiple protein sequence alignment from tertiary structure comparison: assignment of global and residue confidence levels. Proteins. Wiley; 1992;14:309–23.

81. Roberts E, Eargle J, Wright D, Luthey-Schulten Z. MultiSeq: unifying sequence and structure data for evolutionary analysis. BMC Bioinformatics [Internet]. Springer Science and Business Media LLC; 2006;7. 10.1186/1471-2105-7-382

82. Humphrey W, Dalke A, Schulten K. VMD: visual molecular dynamics. J Mol Graph. Elsevier BV; 1996;14:33–8, 27–8.

83. Joo S, Cho IJ, Seo H, Son HF, Sagong H-Y, Shin TJ, et al. Structural insight into molecular mechanism of poly(ethylene terephthalate) degradation. Nat Commun [Internet]. Springer Science and Business Media LLC; 2018;9. 10.1038/s41467-018-02881-1

84. Palm GJ, Reisky L, Böttcher D, Müller H, Michels EAP, Walczak MC, et al. Structure of the plastic-degrading Ideonella sakaiensis MHETase bound to a substrate. Nat Commun. 2019;10:1717.

85. Gautom T, Dheeman D, Levy C, Butterfield T, Alvarez Gonzalez G, Le Roy P, et al. Structural basis of terephthalate recognition by solute binding protein TphC. Nat Commun. 2021;12:6244.

86. Bushnell B. BBMap. 2015.

87. Buchfink B, Xie C, Huson DH. Fast and sensitive protein alignment using DIAMOND. Nat Methods. 2015;12:59–60.

88. R Core Team. R: A Language and Environment for Statistical Computing [Internet]. Vienna, Austria: R Foundation for Statistical Computing; 2021. https://www.R-project.org/

89. Kang DD, Li F, Kirton E, Thomas A, Egan R, An H, et al. MetaBAT 2: an adaptive binning algorithm for robust and efficient genome reconstruction from metagenome assemblies. PeerJ. 2019;7:e7359.

90. Wu Y-W, Simmons BA, Singer SW. MaxBin 2.0: an automated binning algorithm to recover genomes from multiple metagenomic datasets [Internet]. Bioinformatics. 2016. p. 605–7. 10.1093/bioinformatics/btv638

91. Alneberg J, Bjarnason BS, de Bruijn I, Schirmer M, Quick J, Ijaz UZ, et al. Binning metagenomic contigs by coverage and composition [Internet]. Nature Methods. 2014. p. 1144–6. 10.1038/nmeth.3103

92. Parks DH, Imelfort M, Skennerton CT, Hugenholtz P, Tyson GW. CheckM: assessing the quality of microbial genomes recovered from isolates, single cells, and metagenomes. Genome Res. 2015;25:1043–55.

93. Delmont TO, Quince C, Shaiber A, Esen ÖC, Lee STM, Rappé MS, et al. Nitrogen-fixing populations of Planctomycetes and Proteobacteria are abundant in surface ocean metagenomes [Internet]. Nature Microbiology. 2018. p. 804–13. 10.1038/s41564-018-0176-9

94. Olm MR, Brown CT, Brooks B, Banfield JF. dRep: a tool for fast and accurate genomic comparisons that enables improved genome recovery from metagenomes through de-replication [Internet]. The ISME Journal. 2017. p. 2864–8. 10.1038/ismej.2017.126

95. Hyatt D, Chen G-L, Locascio PF, Land ML, Larimer FW, Hauser LJ. Prodigal: prokaryotic gene recognition and translation initiation site identification. BMC Bioinformatics. 2010;11:119.

96. Chaumeil P-A, Mussig AJ, Hugenholtz P, Parks DH. GTDB-Tk: a toolkit to classify genomes with the Genome Taxonomy Database. Bioinformatics. Oxford University Press (OUP); 2019;36:1925–7.

97. Katoh K, Rozewicki J, Yamada KD. MAFFT online service: multiple sequence alignment, interactive sequence choice and visualization. Brief Bioinform. Oxford University Press (OUP); 2019;20:1160–6.

98. Nguyen L-T, Schmidt HA, von Haeseler A, Minh BQ. IQ-TREE: a fast and effective stochastic algorithm for estimating maximum-likelihood phylogenies. Mol Biol Evol. Oxford University Press (OUP); 2015;32:268–74.

99. Trifinopoulos J, Nguyen L-T, von Haeseler A, Minh BQ. W-IQ-TREE: a fast online phylogenetic tool for maximum likelihood analysis. Nucleic Acids Res. 2016;44:W232–5.

100. Letunic I, Bork P. Interactive Tree Of Life (iTOL) v4: recent updates and new developments. Nucleic Acids Res. 2019;47:W256–9.

101. Aroney STN, Newell RJP, Nissen JN, Camargo AP, Tyson GW, Woodcroft BJ. CoverM: read alignment statistics for metagenomics. Bioinformatics [Internet]. 2025;41. 10.1093/bioinformatics/btaf147

102. Li H. Minimap2: pairwise alignment for nucleotide sequences. Bioinformatics. 2018;34:3094–100.

103. Alcorta J, Alarcón-Schumacher T, Salgado O, Díez B. Taxonomic Novelty and Distinctive Genomic Features of Hot Spring Cyanobacteria [Internet]. Frontiers in Genetics. 2020. 10.3389/fgene.2020.568223

104. Sonnendecker C, Oeser J, Konstantin Richter P, Hille P, Zhao Z, Fischer C, et al. Low Carbon Footprint Recycling of Post-Consumer PET Plastic with a Metagenomic Polyester Hydrolase [Internet]. ChemSusChem. 2022. 10.1002/cssc.202101062

105. Bell EL, Smithson R, Kilbride S, Foster J, Hardy FJ, Ramachandran S, et al. Directed evolution of an efficient and thermostable PET depolymerase. Nature Catalysis. Nature Publishing Group; 2022;5:673–81.

106. Marchler-Bauer A, Bryant SH. CD-Search: protein domain annotations on the fly. Nucleic Acids Res. Oxford University Press (OUP); 2004;32:W327–31.

107. Blázquez-Sánchez P, Vargas JA, Furtado AA, Griñen A, Leonardo DA, Sculaccio SA, et al. Engineering the catalytic activity of an Antarctic PET-degrading enzyme by loop exchange. Protein Sci. 2023;32:e4757.

108. Griñen A, Eltit-Villarroel V, Durán-Osorio F, Avilés J, Zacconi FC, Cárcamo Noriega EN, et al. MPNN-guided redesign of PET hydrolases with enhanced catalytic activity below the PET glass transition temperature. ACS Sustain Chem Eng. American Chemical Society (ACS); 2026;14:17074–86.

109. Alin A. Minitab. Wiley Interdiscip Rev Comput Stat. Wiley; 2010;2:723–7.

110. Magalhães RP, Cunha JM, Sousa SF. Perspectives on the role of enzymatic biocatalysis for the degradation of plastic PET. Int J Mol Sci. MDPI AG; 2021;22:11257.

111. Kwon J, Koh S, Jang S, Cho H, Shin M, Jeon H, et al. Improving yield and thermostability of PETase as a maltose binding protein fusion in the periplasm of Escherichia coli. Int J Mol Sci. MDPI AG; 2026;27:2962.

112. Sasoh M, Masai E, Ishibashi S, Hara H, Kamimura N, Miyauchi K, et al. Characterization of the terephthalate degradation genes of Comamonas sp. strain E6. Appl Environ Microbiol. 2006;72:1825–32.

113. Hosaka M, Kamimura N, Toribami S, Mori K, Kasai D, Fukuda M, et al. Novel tripartite aromatic acid transporter essential for terephthalate uptake in Comamonas sp. strain E6. Appl Environ Microbiol. American Society for Microbiology; 2013;79:6148–55.

114. Sagong H-Y, Seo H, Kim T, Son HF, Joo S, Lee SH, et al. Decomposition of the PET film by MHETase using exo-PETase function. ACS Catal. American Chemical Society (ACS); 2020;10:4805–12.

115. Zrimec J, Kokina M, Jonasson S, Zorrilla F, Zelezniak A. Plastic-degrading potential across the global microbiome correlates with recent pollution trends. MBio. American Society for Microbiology; 2021;12:e0215521.

116. Urbanek AK, Rymowicz W, Mirończuk AM. Degradation of plastics and plastic-degrading bacteria in cold marine habitats. Appl Microbiol Biotechnol. Springer Science and Business Media LLC; 2018;102:7669–78.

117. Jacquin J, Cheng J, Odobel C, Pandin C, Conan P, Pujo-Pay M, et al. Microbial ecotoxicology of marine plastic debris: A review on colonization and biodegradation by the “plastisphere.” Front Microbiol. Frontiers Media SA; 2019;10:865.

118. Berríos-Farías V, Guajardo-Leiva S, Gallardo-Cerda J, Galbán-Malagón C, Egas C, Molina-Montenegro MA, et al. Metagenomic insights into potential PET hydrolases from Antarctic soils and rhizospheres. Front Microbiol. Frontiers Media SA; 2026;17:1749101.

119. Ghiglione J-F, Galand PE, Pommier T, Pedrós-Alió C, Maas EW, Bakker K, et al. Pole-to-pole biogeography of surface and deep marine bacterial communities. Proc Natl Acad Sci U S A. Proceedings of the National Academy of Sciences; 2012;109:17633–8.

120. Vass M, Langenheder S. The legacy of the past: effects of historical processes on microbial metacommunities. Aquat Microb Ecol. Inter-Research Science Center; 2017;79:13–9.

121. Gambarini V, Drost CJ, Kingsbury JM, Weaver L, Pantos O, Handley KM, et al. Uncoupled: investigating the lack of correlation between the transcription of putative plastic-degrading genes in the global ocean microbiome and marine plastic pollution. Environ Microbiome. Springer Science and Business Media LLC; 2024;19:34.

122. Khersonsky O, Roodveldt C, Tawfik DS. Enzyme promiscuity: evolutionary and mechanistic aspects. Curr Opin Chem Biol. Elsevier BV; 2006;10:498–508.

123. Rauwerdink A, Kazlauskas RJ. How the Same Core Catalytic Machinery Catalyzes 17 Different Reactions: the Serine-Histidine-Aspartate Catalytic Triad of α/β-Hydrolase Fold Enzymes. ACS Catal. 2015;5:6153–76.

124. Edwards S, Rice D, Palomino P, Newton I, Mellies JL. Ancestral hydrocarbon metabolism enables PET degradation by a natural bacterial consortium. mSystems. American Society for Microbiology; 2026;11:e0082926.

