## Supplementary File 1 for "Polar Marine Microbial Communities as Reservoirs of Polyester Degrading Enzymes"

### SUPPLEMENTARY FIGURES

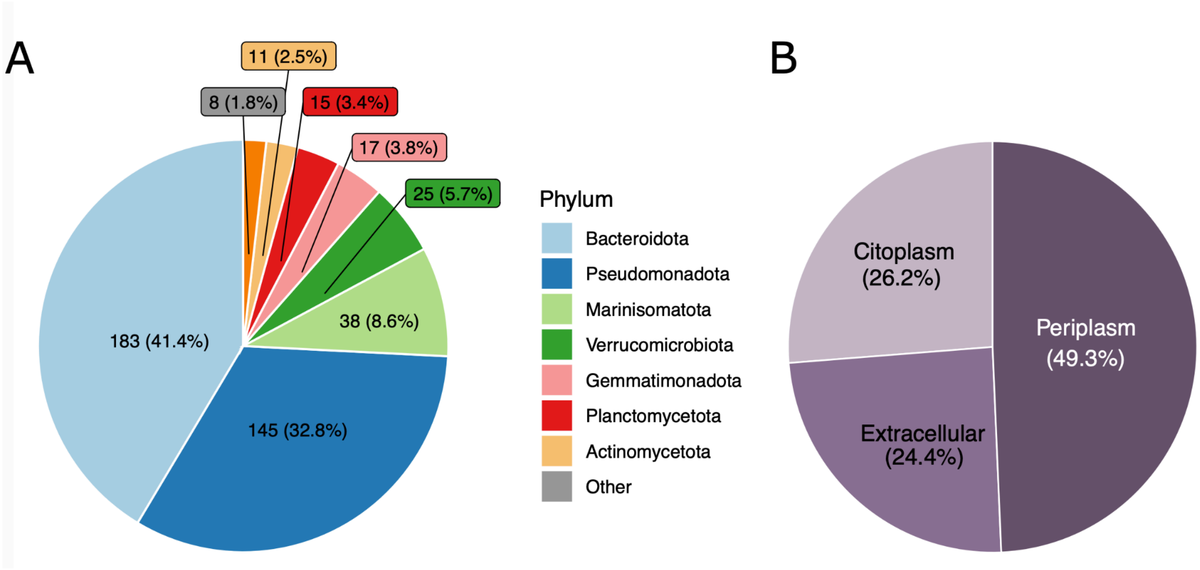

**Supplementary Fig. S1. Taxonomic assignments and predicted subcellular localization of PETase-like sequences from marine metagenomes.** (A) Phylum-level taxonomic affiliation of the 443 non-redundant putative PETase-like sequences (B) Predicted subcellular localization of the same set of putative PETase-like sequences.

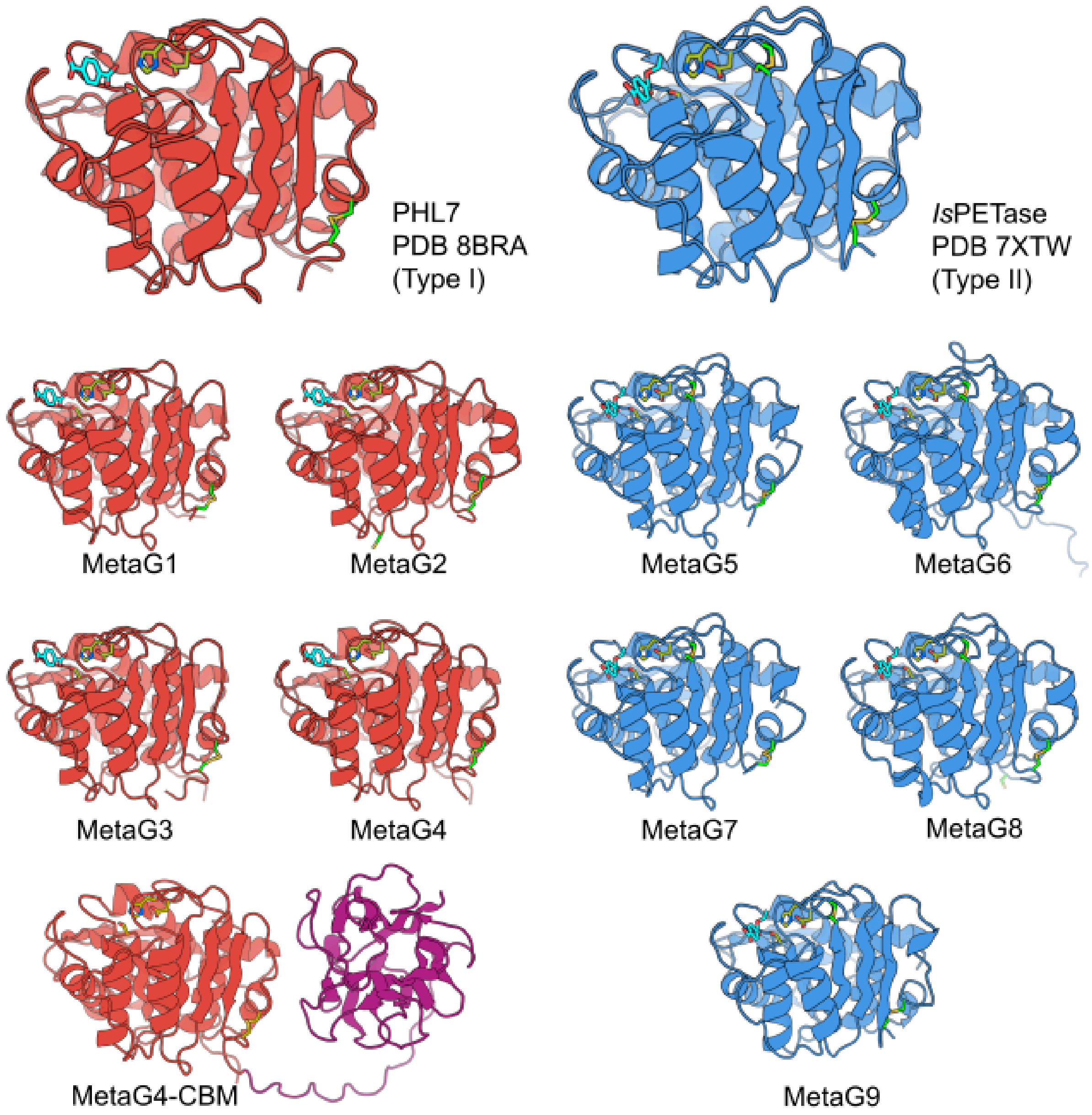

**Supplementary Fig. S2. ColabFold-predicted three-dimensional structures of MetaG1-MetaG9.** Cartoon representation of all Type I (red) and Type II (blue) enzymes selected from the high-confidence PETase-like clade, as well as the most efficient natural representatives found for each class so far, corresponding to PHL7 (experimental structure bound to TPA, PDB 8BRA) and *Is*PETase (experimental structure bound to MHET, PDB 7XTW). The ligands bound to the experimental structures of PHL7 and *Is*PETase are shown in cyan sticks, and maintained in the ColabFold-predicted structures of MetaG1-MetaG9 for reference. Catalytic residues are shown in green sticks and disulfide bonds in yellow sticks. Type II enzymes have an additional disulfide bond in the active site. For MetaG4, the full-length sequence contains a CBM domain, shown in purple.

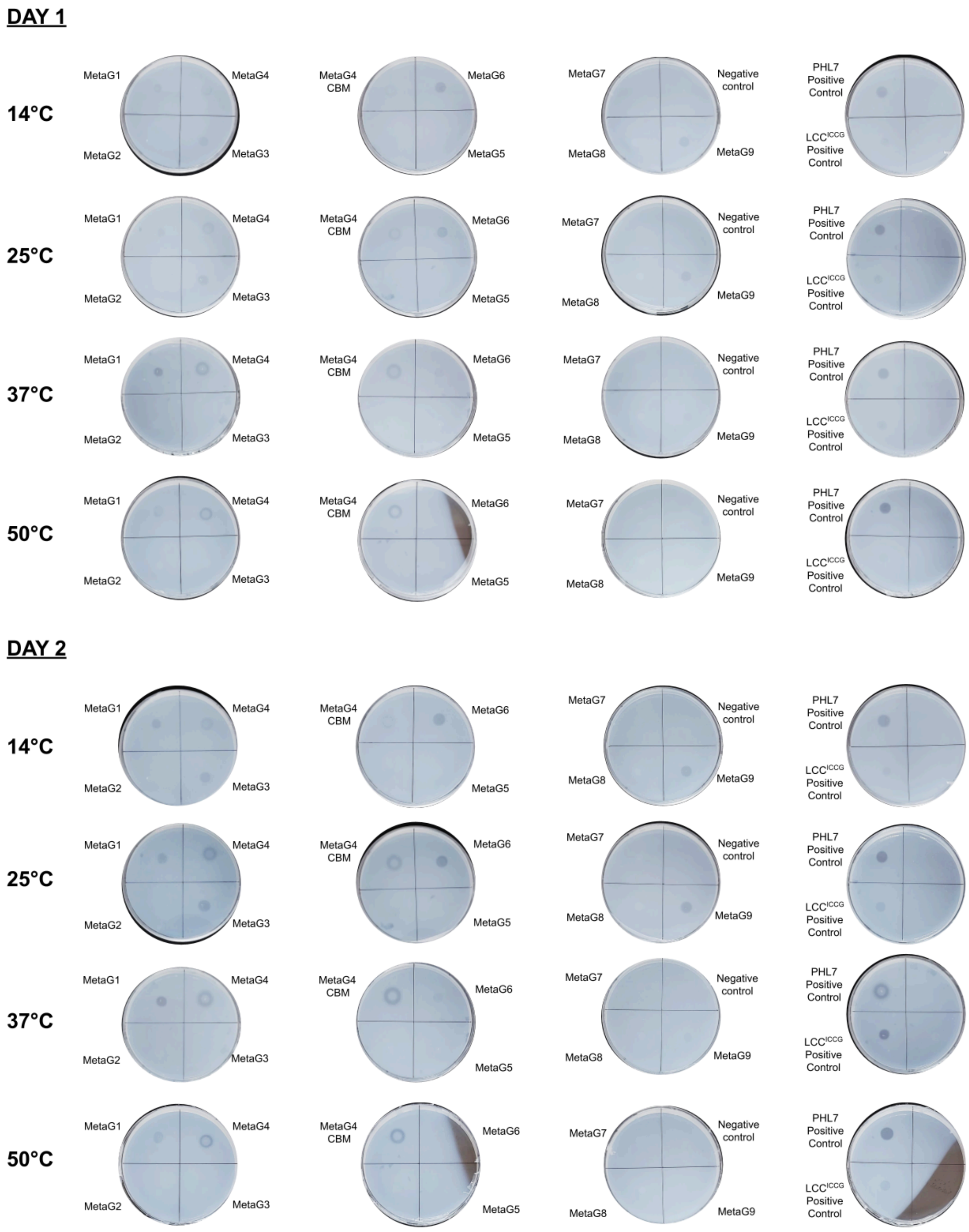

**Supplementary Fig. S3. PCL plate degradation assays for MetaG1-MetaG9 at different temperatures and days of incubation.** LB agar plates with PCL nanoparticles, inoculated with crude extracts of *E. coli* BL21(DE3) expressing the different enzymes, were incubated for 1 (top) and 2 days (bottom) at different temperatures ranging from 14ºC to 50ºC. PHL7 and $LCC_{ICCG}$ were used as positive controls and TEM 1 beta-lactamase as negative control for PCL degradation activity.

| Names | Active site | | | Catalytic site cysteines | | Oxyanion hole | Catalytic activity-relevant residues | | | | | |
|---|---|---|---|---|---|---|---|---|---|---|---|---|
| WP 080747404.1 | S | D | H | C | C | G | G | E | R | F | F | F |
| MHETase | S | D | H | C | C | G | S | E | R | F | F | F |
| gene 811634 | S | D | H | C | C | G | G | E | R | S | F | F |
| WP 083293388.1 | S | D | H | C | C | G | G | E | R | S | F | F |
| gene 362593 | S | D | H | C | C | G | G | G | I | L | F | F |
| gene 907838 | S | D | H | C | C | G | G | T | - | - | - | F |
| gene 631658 | S | D | H | C | C | G | G | N | - | L | F | F |
| gene 489853 | S | D | H | C | C | G | G | N | - | L | F | F |
| gene 1473514 | S | D | H | C | C | G | G | N | - | L | F | F |
| gene 700226 | S | D | H | C | C | G | G | N | Q | R | F | F |
| gene 463040 | S | D | H | C | C | G | G | N | - | R | F | F |
| gene 1102685 | S | D | H | C | C | G | G | N | - | R | F | F |
| gene 867315 | S | D | H | C | C | G | G | N | - | R | F | F |
| NODE 151 length 60169 cov 17.1422 2 | S | D | H | C | C | G | G | N | - | R | F | F |
| gene 1033182 | S | D | H | C | C | G | G | N | - | L | I | L |
| gene 941996 | S | D | H | C | C | G | G | N | - | L | V | L |
| gene 1614930 | S | D | H | C | C | G | G | T | - | - | A | L |
| gene 1336282 | S | D | H | C | C | G | G | T | - | - | A | L |
| NODE 63191 length 1194 cov 2.189640 1 | S | D | H | C | C | G | G | D | F | I | F | I |
| gene 548027 | S | D | H | C | C | G | G | G | G | I | F | I |
| gene 275925 | S | D | H | C | C | G | G | G | G | I | F | I |
| gene 598154 | S | D | H | C | C | G | G | G | G | I | F | I |
| gene 612768 | S | D | H | C | C | G | G | G | G | I | M | C |
| gene 612765 | S | D | H | C | C | G | G | G | A | I | F | C |
| gene 1188405 | S | D | H | C | C | G | G | T | - | L | F | V |
| gene 546387 | S | D | H | C | C | G | G | T | - | L | F | V |
| gene 803527 | S | D | H | C | C | G | G | T | - | L | F | V |
| gene 380150 | S | D | H | C | C | G | G | T | - | L | F | V |
| gene 685800 | S | D | H | C | C | G | G | T | - | I | F | T |
| gene 704409 | S | D | H | C | C | G | G | T | - | P | F | S |
| gene 1007646 | S | D | H | C | C | G | G | T | - | P | F | S |
| gene 1769998 | S | D | H | C | C | G | G | T | - | P | F | S |
| gene 800034 | S | D | H | C | C | G | G | T | - | V | F | I |
| gene 1317527 | S | D | H | C | C | G | G | T | - | V | F | I |
| gene 694486 | S | D | H | C | C | G | G | T | - | V | F | I |
| gene 714036 | S | D | H | C | C | G | G | T | - | V | F | I |
| gene 427082 | S | D | H | C | C | G | G | T | - | V | F | I |
| NODE 274 length 38879 cov 5.258448 12 | S | D | H | C | C | G | G | T | - | V | F | I |
| gene 214653 | S | D | H | C | C | G | G | N | W | A | R | I |
| gene 1241012 | S | D | H | C | C | G | G | N | P | L | G | V |
| gene 287293 | S | D | H | C | C | G | G | N | P | L | G | V |
| gene 325030 | S | D | H | C | C | G | G | N | P | L | G | V |
| gene 934192 | S | D | H | C | C | G | G | N | P | L | G | V |
| gene 82884 | S | D | H | C | C | G | G | N | P | L | G | V |
| gene 1168518 | S | D | H | C | C | G | G | N | P | L | G | V |
| gene 273559 | S | D | H | C | C | G | G | N | P | L | G | V |
| gene 284216 | S | D | H | C | C | G | G | N | L | F | L | L |
| NODE 54155 length 1326 cov 2.498820 1 | S | D | H | C | C | G | G | N | Y | F | L | L |
| gene 824229 | S | D | H | C | C | G | G | R | - | F | I | L |
| gene 79171 | S | D | H | C | C | G | G | R | - | F | I | L |
| gene 1330454 | S | D | H | C | C | G | G | R | - | F | V | L |
| gene 853210 | S | D | H | C | C | G | G | R | - | F | V | L |
| gene 825251 | S | D | H | C | C | G | G | R | - | F | V | L |
| gene 280146 | S | D | H | C | C | G | G | R | - | F | V | V |
| gene 1119716 | S | D | H | C | C | G | G | R | - | F | I | V |
| gene 940652 | S | D | H | C | C | G | G | R | - | F | V | L |
| gene 730180 | S | D | H | C | C | G | G | R | - | F | V | L |
| gene 631927 | S | D | H | C | C | G | G | R | - | F | V | L |
| gene 416980 | S | D | H | C | C | G | G | T | - | - | - | I |
| gene 338437 | S | D | H | C | C | G | G | T | - | - | - | V |
| gene 90570 | S | D | H | C | C | G | G | T | - | - | - | V |
| gene 960259 | S | D | H | C | C | G | G | T | - | Y | V | I |
| gene 1206697 | S | D | H | C | C | G | G | T | - | Y | V | I |
| gene 112801 | S | D | H | C | C | G | H | L | - | - | V | M |
| gene 279159 | S | D | H | C | C | G | G | S | - | L | M | I |
| gene 128585 | S | D | H | C | C | G | G | T | - | - | V | I |
| gene 1032598 | S | D | H | C | C | G | G | G | - | L | V | L |
| gene 499058 | S | D | H | C | C | G | G | N | - | L | V | L |
| gene 1109367 | S | D | H | C | C | G | G | K | - | A | V | I |
| gene 691950 | S | D | H | C | C | G | G | K | - | A | V | I |
| gene 1410989 | S | D | H | C | C | G | G | K | - | A | V | I |
| gene 1930325 | S | D | H | C | C | G | G | K | - | A | V | I |
| gene 454141 | S | D | H | C | C | G | G | K | - | G | V | I |
| gene 443322 | S | D | H | C | C | G | G | K | - | G | V | I |
| gene 297145 | S | D | H | C | C | G | G | K | - | G | V | I |
| gene 1231951 | S | D | H | C | C | G | G | K | - | G | V | I |
| gene 327842 | S | D | H | C | C | G | G | K | - | G | V | I |
| gene 901675 | S | D | H | C | C | G | G | K | - | G | V | I |
| NODE 25871 length 1378 cov 2.007559 1 | S | D | H | C | C | G | G | K | - | G | V | I |
| NODE 20111 length 3344 cov 6.96899 1 | S | D | H | C | C | G | G | K | - | G | V | I |

**Supplementary Fig. S4. Residue conservation of identified MHETase-like enzymes in this study**. Residue conservation of the active site catalytic residues, the catalytic site cysteines, the oxyanion hole, and the catalytic activity-relevant residues S131, E226, R411, F415, F424 and F495, derived from a MSA of all 209 putative MHETases found in the metagenomes analyzed in this work.

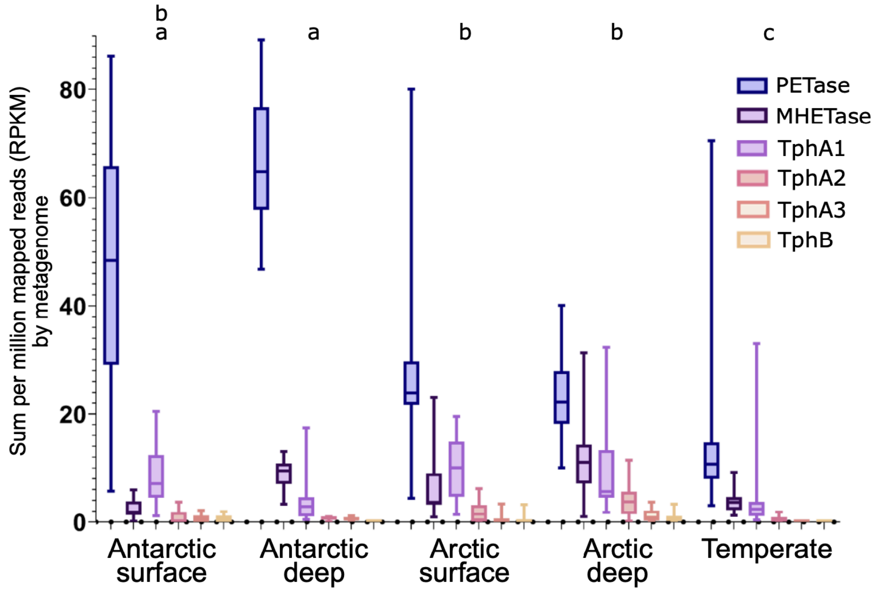

**Supplementary Fig. S5. Abundance of enzymes related to the PET degradation pathway in marine metagenomes.** Bar graph representing the sum of the abundance in reads per kilobase per million mapped reads (RPKM) of the enzyme sequences involved in the PET degradation pathway that were identified in marine metagenomes, classified according to their PET (PETase), MHET (MHETase) and TPA degradation activity (TphA1, TphA2, TphA3, TphB) and the geographical location of the metagenomic sample. Letters a, b, and c represent significant differences between motif groups in each area, based on multifactorial ANOVA with Tukey's a posteriori test (Supplementary Table S6).

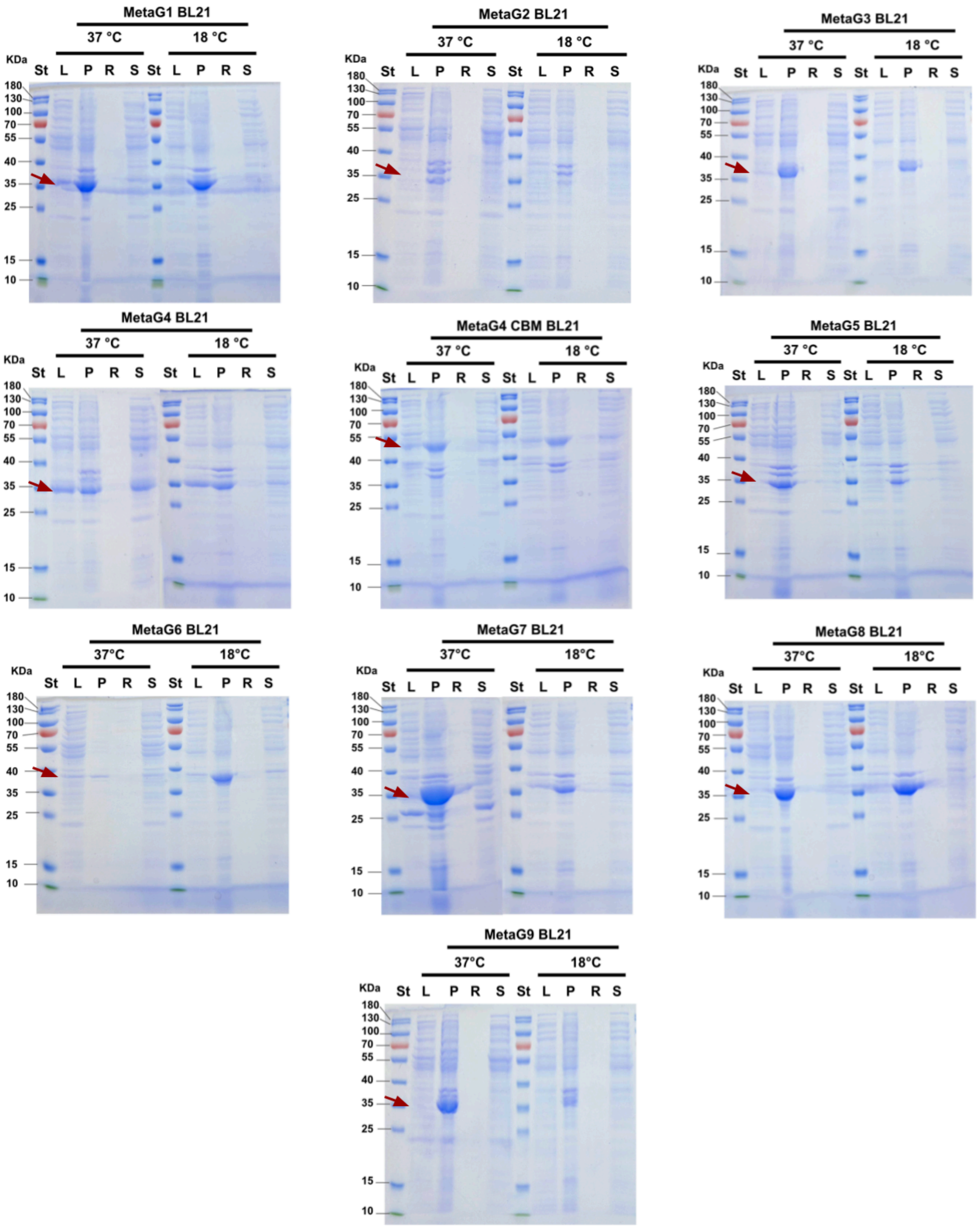

**Supplementary Fig. S6. Expression test of MetaG1-MetaG9 enzymes in *E. coli* BL21 (DE3).** SDS-PAGE of the enzyme expression test of MetaG1-MetaG9 enzymes in the recombinant expression host *E. coli* BL21 (DE3) under two different IPTG induction temperature conditions, 37 and 18 ºC. In each gel, St corresponds to the molecular weight standards, L to the cell lysate, P to the cell pellet, R to the soluble fraction after purification with Ni-NTA resin, and S to the lysate of non-induced cells as negative controls. The arrows indicate the expected molecular weight for each enzyme.

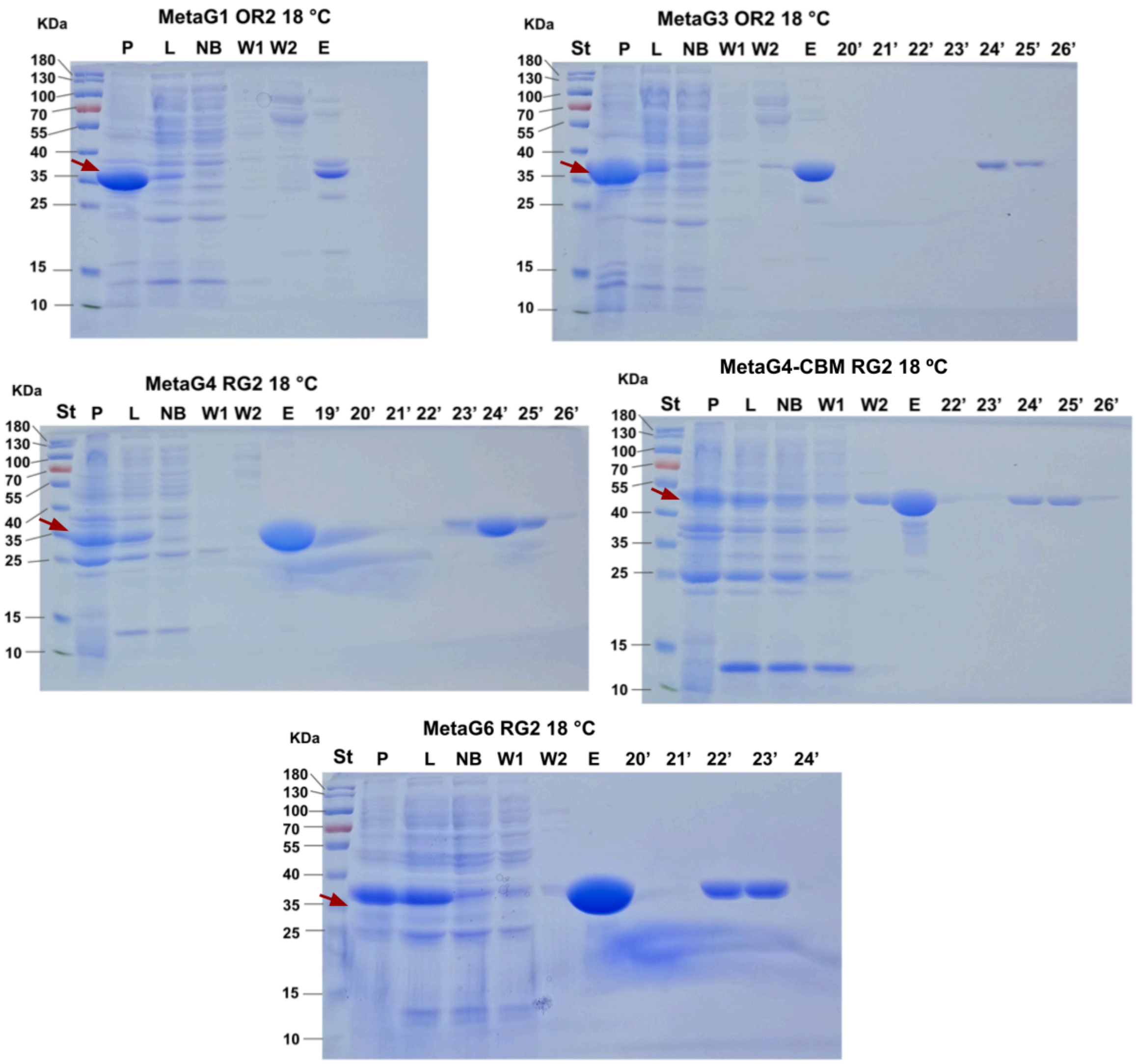

**Supplementary Fig. S7. SDS-PAGE of the successful purification of active MetaG enzymes in different *E. coli* strains.** Enzymes MetaG1, MetaG3, MetaG4, MetaG4-CBM, MetaG6 and MetaG9 were transformed into *E. coli* Origami 2 DE3 pLysS (OR2) and *E. coli* Rosetta-Gami 2 DE3 pLysS (RG2) and purified as described in *Methods*. The strain and temperature conditions for induction of protein expression are indicated above each gel. St corresponds to the molecular weight standards, P to the cell pellet, L to the cell lysate, NB to the front load of the lysate onto the HisTrap columns, W1 and W2 to the wash fraction of the columns, E to the elution fraction after purification with buffer supplemented with 250 mM imidazol and the numbers to the elution time after size exclusion chromatography on an HPLC connected to a Superdex S75 column. MetaG9 is not shown as it did not solubly express in either *E. coli* strain tested. The arrows indicate the expected molecular weight for each enzyme.

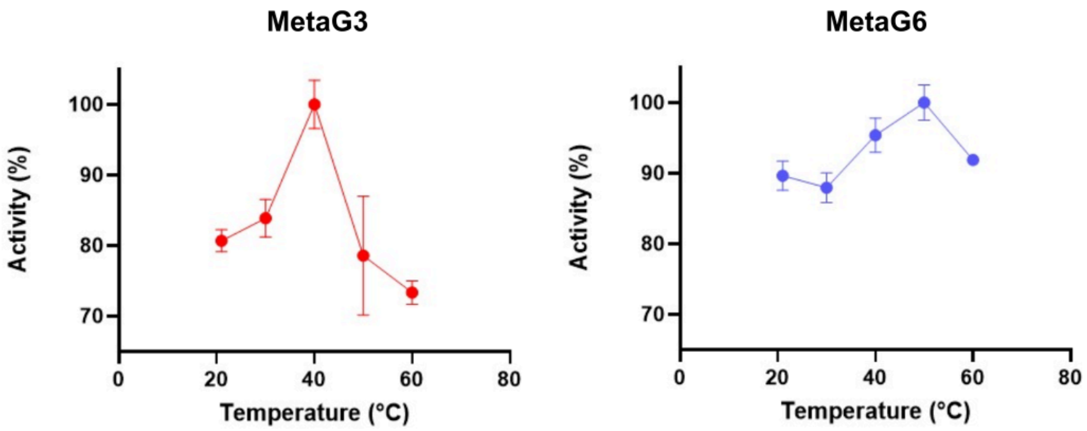

**Supplementary Fig. S8. Determination of the optimal temperature for the enzymatic PCL nanoparticle degradation reactions using MetaG3 and MetaG6.** Relative initial hydrolysis rates of PCL nanoparticles by MetaG3 and MetaG6 at different reaction temperatures, ranging from 20 to 60 ºC, using a concentration of 8 µg/ml of MetaG3 and 4 µg/ml of MetaG6 against 0.79 mg/mL PCL nanoparticles in 200 µl of reaction buffer. The whiskers indicate the standard deviation from three technical replicates.

### SUPPLEMENTARY TABLES

**Supplementary Table S1. Classification of amino acid motifs according to conserved amino acid pattern.** The colors represent the conserved amino acid residues for the catalytic triad (red), oxyanion hole (blue) and aromatic clamp (green). The brackets indicate a less conserved amino acid pattern position. The letter x shows a non-conserved position in the amino acid pattern. Numbers indicate the number of residues between conserved patterns. Patterns are additive, meaning that in order to classify into the next motif the sequence must classify into the previous motif.

| Motif | Amino acid pattern |
|---|---|
| M0 | Catalytic triad (S, D, H) |
| M3 | GxS[MKL]GG[GA][GA]x(~20)[WYF]x(19-27)Dx[IVLTFP]X(~30)H |
| M3 strict | GxS[MK]GGGGx(17-20)[WYF]x(19-23)Dx[IVLTFP]x(27-31)H |
| M4 | [PG]G[YF]x(~70)-M3 |
| M4 strict | [PG]G[YF]x(65-75)-M3 |
| M5 | M4-Hx(20-25)[ND]xDxR[YF]xxFx[CY] |

**Supplementary Table S2. PETase-like motif classification, taxonomy assignment, and subcellular localization prediction.** All enzymes belong to the Bacteria domain.

| PETase-like Enzyme | Closest Genome | Identity | Phylum | Class | Order | Family | Genus | Species | Cytoplasm | Periplasm | Extracellular | Potential Location | Mx |
|---|---|---|---|---|---|---|---|---|---|---|---|---|---|
| gene_1012555\|GeneMark.hmm\|216_aa\|-\|2\|649_512307 | GB_GCA_002721895.1 | 90.8 | Verrucomicrobiota | Verrucomicrobiae | Limisphaerales | UBA1100 | UBA6614 | UBA6614 sp002721895 | 0.486 | 0.000 | 0.008 | Cytoplasm | M0 |
| gene_1085474\|GeneMark.hmm\|282_aa\|+\|20618\|21466_535898 | RS_GCF_002890895.1 | 99.6 | Pseudomonadota | Gammaproteobacteria | Pseudomonadales | Pseudomonadaceae | Stutzerimonas | Stutzerimonas stutzeri_AB | 0.005 | 0.988 | 0.012 | Periplasm | M0 |
| gene_1087197\|GeneMark.hmm\|229_aa\|-\|2\|688_560872 | GB_GCA_002708345.1 | 53.8 | Planctomycetota | Planctomycetia | Pirellulales | Pirellulaceae | SAT2750 | SAT2750 sp002708345 | 0.549 | 0.003 | 0.070 | Cytoplasm | M0 |
| gene_109985\|GeneMark.hmm\|293_aa\|-\|1098\|1979_55471 | RS_GCF_000355675.1 | 98.0 | Pseudomonadota | Gammaproteobacteria | Pseudomonadales | DSM-6294 | Thalassolituus | Thalassolituus oleivorans | 0.004 | 0.125 | 0.875 | Extracellular | M0 |
| gene_1121959\|GeneMark.hmm\|133_aa\|-\|3\|401_591996 | GB_GCA_016873095.1 | 71.8 | Planctomycetota | Planctomycetia | Gemmatales | Gemmataceae | VGYC01 | VGYC01 sp016873095 | 0.304 | 0.023 | 0.050 | Cytoplasm | M0 |
| gene_112573\|GeneMark.hmm\|407_aa\|+\|51153\|52376_66789 | GB_GCA_024644285.1 | 84.9 | Pseudomonadota | Alphaproteobacteria | Rhodobacterales | Rhodobacteraceae | JAHELH01 | JAHELH01 sp024644285 | 0.978 | 0.000 | 0.000 | Cytoplasm | M0 |
| gene_1143709\|GeneMark.hmm\|386_aa\|+\|1209\|2369_554038 | GB_GCA_913020075.1 | 99.5 | Bacteroidota | Bacteroidia | Flavobacteriales | Flavobacteriaceae | MS024-2A | MS024-2A sp913020075 | 0.004 | 1.000 | 0.000 | Periplasm | M0 |
| gene_1160958\|GeneMark.hmm\|244_aa\|-\|33\|767_598736 | GB_GCA_002721705.1 | 87.2 | Verrucomicrobiota | Verrucomicrobiae | Opitutales | MB11C04 | UBA2993 | UBA2993 sp002721705 | 0.839 | 0.001 | 0.045 | Cytoplasm | M0 |
| gene_1171882\|GeneMark.hmm\|131_aa\|-\|2\|394_617080 | GB_GCA_022560615.1 | 84.7 | Gemmatimonadota | Gemmatimonadetes | Longimicrobiales | UBA6960 | UBA1138 | UBA1138 sp022560615 | 0.653 | 0.000 | 0.009 | Cytoplasm | M0 |
| gene_1191696\|GeneMark.hmm\|269_aa\|-\|59\|868_594534 | GB_GCA_937775205.1 | 79.8 | Verrucomicrobiota | Verrucomicrobiae | Limisphaerales | UBA1100 | JACNJL01 | JACNJL01 sp937775205 | 0.015 | 0.012 | 0.932 | Extracellular | M0 |
| gene_1209087\|GeneMark.hmm\|401_aa\|+\|1108\|2313_583256 | GB_GCA_000981645.1 | 78.4 | Bacteroidota | Bacteroidia | Flavobacteriales | UA16 | ASP10-05a | ASP10-05a sp000981645 | 0.004 | 1.000 | 0.000 | Periplasm | M0 |
| gene_120993\|GeneMark.hmm\|196_aa\|+\|1\|591_62053 | GB_GCA_937775205.1 | 88.3 | Verrucomicrobiota | Verrucomicrobiae | Limisphaerales | UBA1100 | JACNJL01 | JACNJL01 sp937775205 | 0.676 | 0.000 | 0.024 | Cytoplasm | M0 |
| gene_1252150\|GeneMark.hmm\|167_aa\|+\|2\|505_624110 | GB_GCA_012960305.1 | 94.0 | Pseudomonadota | Gammaproteobacteria | Pseudomonadales | Pseudohongiellaceae | UBA9145 | UBA9145 sp012960305 | 0.066 | 0.005 | 0.344 | Extracellular | M0 |
| gene_1261948\|GeneMark.hmm\|325_aa\|-\|4054\|5031_682411 | GB_GCA_028821775.1 | 74.6 | Verrucomicrobiota | Verrucomicrobiae | Verrucomicrobiales | Akkermansiaceae | Roseibacillus_B | Roseibacillus_B sp028821775 | 0.005 | 0.045 | 0.954 | Extracellular | M0 |
| gene_1275123\|GeneMark.hmm\|134_aa\|-\|3\|404_619116 | GB_GCA_949372635.1 | 97.0 | Pseudomonadota | Gammaproteobacteria | Pseudomonadales | Pseudohongiellaceae | UBA9145 | UBA9145 sp949372635 | 0.075 | 0.002 | 0.152 | Extracellular | M0 |
| gene_1292303\|GeneMark.hmm\|223_aa\|-\|2\|670_664316 | GB_GCA_002708465.1 | 92.4 | Planctomycetota | UBA8108 | UBA8108 | UBA8108 | UBA8108 | UBA8108 sp002708465 | 0.005 | 0.000 | 0.999 | Extracellular | M0 |
| gene_130721\|GeneMark.hmm\|256_aa\|+\|540\|1310_69104 | GB_GCA_013215655.1 | 83.1 | Pseudomonadota | Gammaproteobacteria | Enterobacterales | Alteromonadaceae | JABSOX01 | JABSOX01 sp013215655 | 0.976 | 0.000 | 0.000 | Cytoplasm | M0 |
| gene_1317505\|GeneMark.hmm\|270_aa\|-\|4\|816_634786 | GB_GCA_030652635.1 | 75.6 | Pseudomonadota | Gammaproteobacteria | Pseudomonadales | Pseudohongiellaceae | Pseudohongiella | Pseudohongiella sp030652635 | 0.004 | 0.011 | 0.987 | Extracellular | M0 |

| | | | | | | | | | | | | | |
|---|---|---|---|---|---|---|---|---|---|---|---|---|---|
| gene_1338719\|GeneMark.hmm\|288_aa\|-\|27\|893_717276 | GB_GCA_905182215.1 | 91.3 | Bacteroidota | Bacteroidia | Flavobacteriales | UBA7430 | UBA6772 | UBA6772 sp905182215 | 0.035 | 0.983 | 0.000 | Periplasm | M0 |
| gene_1356216\|GeneMark.hmm\|149_aa\|+\|3\|452_693333 | GB_GCA_030747505.1 | 93.8 | Verrucomicrobiota | Verrucomicrobiae | Limisphaerales | UBA1100 | UBA1100 | UBA1100 sp030747505 | 0.573 | 0.000 | 0.016 | Cytoplasm | M0 |
| gene_1362649\|GeneMark.hmm\|279_aa\|+\|3\|842_696456 | GB_GCA_913041895.1 | 87.5 | Verrucomicrobiota | Verrucomicrobiae | Limisphaerales | UBA1100 | UBA1100 | UBA1100 sp913041895 | 0.696 | 0.000 | 0.015 | Cytoplasm | M0 |
| gene_136465\|GeneMark.hmm\|207_aa\|+\|3\|626_72385 | GB_GCA_913048385.1 | 86.4 | Pseudomonadota | Gammaproteobacteria | Pseudomonadales | Pseudohongiellaceae | UBA9145 | UBA9145 sp913048385 | 0.311 | 0.001 | 0.011 | Cytoplasm | M0 |
| gene_1369197\|GeneMark.hmm\|346_aa\|+\|42\|1082_763490 | GB_GCA_028821775.1 | 94.2 | Verrucomicrobiota | Verrucomicrobiae | Verrucomicrobiales | Akkermansiaceae | Roseibacillus_B | Roseibacillus_B sp028821775 | 0.009 | 0.018 | 0.966 | Extracellular | M0 |
| gene_1371316\|GeneMark.hmm\|431_aa\|+\|724\|2019_706731 | GB_GCA_000496475.1 | 91.3 | Pseudomonadota | Gammaproteobacteria | Burkholderiales | Burkholderiaceae | RS62 | RS62 sp000496475 | 0.917 | 0.000 | 0.000 | Cytoplasm | M0 |
| gene_1400107\|GeneMark.hmm\|365_aa\|-\|2\|1096_717672 | GB_GCA_937775205.1 | 88.1 | Verrucomicrobiota | Verrucomicrobiae | Limisphaerales | UBA1100 | JACNJL01 | JACNJL01 sp937775205 | 0.005 | 0.001 | 0.999 | Extracellular | M0 |
| gene_1402240\|GeneMark.hmm\|268_aa\|-\|360\|1166_696575 | GB_GCA_913058135.1 | 90.3 | Pseudomonadota | Gammaproteobacteria | Pseudomonadales | Pseudohongiellaceae | Pseudohongiella | Pseudohongiella sp913058135 | 0.008 | 0.013 | 0.986 | Extracellular | M0 |
| gene_1417276\|GeneMark.hmm\|441_aa\|+\|228\|1553_727820 | GB_GCA_021741155.1 | 70.1 | Bacteroidota | Bacteroidia | UBA10329 | UBA10329 | UBA10329 | UBA10329 sp021741155 | 0.005 | 0.999 | 0.001 | Periplasm | M0 |
| gene_144832\|GeneMark.hmm\|409_aa\|-\|68444\|69673_74673 | GB_GCA_943788915.1 | 85.6 | Actinomycetota | Acidimicrobiia | Acidimicrobiales | UBA10347 | UBA10347 | UBA10347 sp003485885 | 0.984 | 0.000 | 0.000 | Cytoplasm | M0 |
| gene_1455025\|GeneMark.hmm\|402_aa\|-\|42\|1250_749043 | RS_GCF_000155735.2 | 98.8 | Pseudomonadota | Alphaproteobacteria | Rhodobacterales | Rhodobacteraceae | Octadecabacter | Octadecabacter arcticus | 0.979 | 0.000 | 0.000 | Cytoplasm | M0 |
| gene_1491923\|GeneMark.hmm\|348_aa\|+\|708\|1754_763287 | GB_GCA_937896535.1 | 86.2 | Myxococcota | Polyangia | Nannocystales | Nannocystaceae | CALBMM01 | CALBMM01 sp937896535 | 0.004 | 0.993 | 0.006 | Periplasm | M0 |
| gene_1553921\|GeneMark.hmm\|265_aa\|+\|18879\|19676_771020 | GB_GCA_012960305.1 | 98.1 | Pseudomonadota | Gammaproteobacteria | Pseudomonadales | Pseudohongiellaceae | UBA9145 | UBA9145 sp012960305 | 0.006 | 0.037 | 0.962 | Extracellular | M0 |
| gene_1571789\|GeneMark.hmm\|348_aa\|+\|291\|1337_803051 | GB_GCA_902589825.1 | 71.4 | Verrucomicrobiota | Verrucomicrobiae | Verrucomicrobiales | DEV007 | CACIUL01 | CACIUL01 sp902589825 | 0.004 | 0.023 | 0.976 | Extracellular | M0 |
| gene_165765\|GeneMark.hmm\|407_aa\|+\|19485\|20708_87318 | RS_GCF_947492965.1 | 62.6 | Bacteroidota | Bacteroidia | Flavobacteriales | Flavobacteriaceae | Aquimarina | Aquimarina sp947492965 | 0.982 | 0.000 | 0.000 | Cytoplasm | M0 |
| gene_1678976\|GeneMark.hmm\|292_aa\|+\|2816\|3694_844153 | GB_GCA_015664885.1 | 99.3 | Gemmatimonadota | Gemmatimonadetes | Longimicrobiales | UBA6960 | UBA1138 | UBA1138 sp003447875 | 0.006 | 0.011 | 0.988 | Extracellular | M0 |
| gene_173684\|GeneMark.hmm\|454_aa\|-\|7325\|8689_89375 | GB_GCA_913009205.1 | 99.1 | Bacteroidota | Bacteroidia | Flavobacteriales | Crocinitomicaceae | UBA4466 | UBA4466 sp913009205 | 0.005 | 1.000 | 0.000 | Periplasm | M0 |
| gene_197109\|GeneMark.hmm\|311_aa\|-\|2113\|3048_101426 | GB_GCA_011523945.1 | 74.3 | Planctomycetota | Planctomycetia | Pirellulales | Pirellulaceae | Mariniblastus | Mariniblastus sp011523945 | 0.005 | 0.020 | 0.980 | Extracellular | M0 |
| gene_198505\|GeneMark.hmm\|462_aa\|-\|101\|1489_97983 | RS_GCF_007993035.1 | 68.0 | Bacteroidota | Bacteroidia | Flavobacteriales | Vicingaceae | Vicingus | Vicingus serpentipes | 0.004 | 1.000 | 0.000 | Periplasm | M0 |
| gene_209106\|GeneMark.hmm\|131_aa\|+\|2\|397_106870 | GB_GCA_029246605.1 | 91.6 | Pseudomonadota | Gammaproteobacteria | Pseudomonadales | Azotimanducaceae_A | UBA7455 | UBA7455 sp029246605 | 0.055 | 0.012 | 0.238 | Extracellular | M0 |

| | | | | | | | | | | | | | |
|---|---|---|---|---|---|---|---|---|---|---|---|---|---|
| gene_217628\|GeneMark.hmm\|405_aa\|+\|73640\|74857_124341 | RS_GCF_900115185.1 | 99.8 | Bacteroidota | Bacteroidia | Flavobacteriales | Flavobacteriaceae | Algorimicrobium | Algorimicrobium echini | 0.980 | 0.000 | 0.000 | Cytoplasm | M0 |
| gene_217716\|GeneMark.hmm\|290_aa\|+\|24473\|25345_109552 | GB_GCA_002733645.1 | 99.0 | Pseudomonadota | Gammaproteobacteria | Pseudomonadales | DSM-6294 | Oleibacter | Oleibacter sp002733645 | 0.007 | 0.049 | 0.950 | Extracellular | M0 |
| gene_224666\|GeneMark.hmm\|222_aa\|-\|2\|667_119118 | GB_GCA_913041895.1 | 86.4 | Verrucomicrobiota | Verrucomicrobiae | Limisphaerales | UBA1100 | UBA1100 | UBA1100 sp913041895 | 0.642 | 0.000 | 0.015 | Cytoplasm | M0 |
| gene_270641\|GeneMark.hmm\|390_aa\|+\|81\|1253_152373 | GB_GCA_016780365.1 | 78.2 | Bacteroidota | Bacteroidia | Flavobacteriales | Parvicellaceae | MED-G20 | MED-G20 sp016780365 | 0.004 | 1.000 | 0.000 | Periplasm | M0 |
| gene_277699\|GeneMark.hmm\|149_aa\|-\|63\|512_141287 | GB_GCA_017302595.1 | 55.0 | Pseudomonadota | Alphaproteobacteria | Rickettsiales | Rickettsiaceae | GCA-2402195 | GCA-2402195 sp017302595 | 0.977 | 0.000 | 0.000 | Cytoplasm | M0 |
| gene_290504\|GeneMark.hmm\|403_aa\|+\|54664\|55875_148855 | GB_GCA_028280565.1 | 60.7 | Bacteroidota | Bacteroidia | Cytophagales | Cyclobacteriaceae | Fulvivirga | Fulvivirga sp028280565 | 0.971 | 0.000 | 0.000 | Cytoplasm | M0 |
| gene_302440\|GeneMark.hmm\|350_aa\|-\|1343\|2395_155963 | GB_GCA_943788405.1 | 93.1 | Bacteroidota | Bacteroidia | Flavobacteriales | BACL11 | UWMA-0277 | UWMA-0277 sp943788405 | 0.004 | 0.999 | 0.001 | Periplasm | M0 |
| gene_302828\|GeneMark.hmm\|291_aa\|+\|408\|1283_152183 | GB_GCA_016763285.1 | 97.9 | Pseudomonadota | Gammaproteobacteria | Pseudomonadales | Pseudomonadaceae | Halopseudomonas | Halopseudomonas sp016763285 | 0.005 | 0.999 | 0.001 | Periplasm | M0 |
| gene_325311\|GeneMark.hmm\|226_aa\|+\|714\|1394_167384 | GB_GCA_022563405.1 | 91.2 | Gemmatimonadota | Gemmatimonadetes | Longimicrobiales | UBA6960 | UBA822 | UBA822 sp022563405 | 0.611 | 0.007 | 0.349 | Cytoplasm | M0 |
| gene_327288\|GeneMark.hmm\|290_aa\|+\|2509\|3381_191893 | RS_GCF_002197985.1 | 99.0 | Pseudomonadota | Gammaproteobacteria | Pseudomonadales | Pseudomonadaceae | Halopseudomonas | Halopseudomonas aestusnigri | 0.005 | 1.000 | 0.000 | Periplasm | M0 |
| gene_330950\|GeneMark.hmm\|258_aa\|-\|2885\|3661_166315 | GB_GCA_021775495.1 | 52.4 | Pseudomonadota | Alphaproteobacteria | Rhizobiales | Rhizobiaceae | JAJFHT01 | JAJFHT01 sp021775495 | 0.983 | 0.000 | 0.000 | Cytoplasm | M0 |
| gene_331320\|GeneMark.hmm\|388_aa\|-\|2520\|3686_170558 | GB_GCA_905478325.1 | 94.8 | Bacteroidota | Bacteroidia | Flavobacteriales | UA16 | UA16 | UA16 sp002390615 | 0.005 | 0.144 | 0.855 | Extracellular | M0 |
| gene_363000\|GeneMark.hmm\|278_aa\|-\|1\|834_183040 | GB_GCA_000981645.1 | 85.3 | Bacteroidota | Bacteroidia | Flavobacteriales | UA16 | ASP10-05a | ASP10-05a sp000981645 | 0.026 | 0.994 | 0.000 | Periplasm | M0 |
| gene_392030\|GeneMark.hmm\|159_aa\|+\|1\|480_228638 | RS_GCF_017991995.1 | 100.0 | Pseudomonadota | Gammaproteobacteria | Burkholderiales | Burkholderiaceae | Xenophilus | Xenophilus sp017991995 | 0.369 | 0.000 | 0.076 | Cytoplasm | M0 |
| gene_397285\|GeneMark.hmm\|285_aa\|-\|242\|1099_225891 | GB_GCA_002712885.1 | 99.6 | Acidobacteriota | Vicinamibacteria | Vicinamibacterales | UBA8438 | UBA8438 | UBA8438 sp002712885 | 0.006 | 0.038 | 0.960 | Extracellular | M0 |
| gene_411397\|GeneMark.hmm\|272_aa\|+\|380\|1198_220478 | GB_GCA_002724415.1 | 82.4 | Planctomycetota | Planctomycetia | Planctomycetales | Planctomycetaceae | UBA8437 | UBA8437 sp002724415 | 0.005 | 0.006 | 0.990 | Extracellular | M0 |
| gene_414845\|GeneMark.hmm\|420_aa\|+\|1336\|2598_203127 | GB_GCA_913061315.1 | 84.0 | Pseudomonadota | Gammaproteobacteria | GCA-001735895 | GCA-001735895 | GCA-001735895 | GCA-001735895 sp913061315 | 0.974 | 0.000 | 0.000 | Cytoplasm | M0 |
| gene_416483\|GeneMark.hmm\|316_aa\|-\|30116\|31066_205930 | GB_GCA_009497895.1 | 80.3 | Pseudomonadota | Gammaproteobacteria | Pseudomonadales | Pseudomonadaceae | Halopseudomonas | Halopseudomonas pelagia_A | 0.024 | 0.879 | 0.114 | Periplasm | M0 |
| gene_416485\|GeneMark.hmm\|308_aa\|-\|31749\|32675_205931 | GB_GCA_009497895.1 | 66.0 | Pseudomonadota | Gammaproteobacteria | Pseudomonadales | Pseudomonadaceae | Halopseudomonas | Halopseudomonas pelagia_A | 0.011 | 0.686 | 0.311 | Periplasm | M0 |
| gene_446518\|GeneMark.hmm\|383_aa\|-\|2977\|4128_220359 | GB_GCA_905182175.1 | 88.5 | Bacteroidota | Bacteroidia | Flavobacteriales | Vicingaceae | NORP294 | NORP294 sp905182175 | 0.004 | 0.991 | 0.009 | Periplasm | M0 |

| | | | | | | | | | | | | | |
|---|---|---|---|---|---|---|---|---|---|---|---|---|---|
| gene_451376\|GeneMark.hmm\|127_aa\|-\|727\|1110_221382 | GB_GCA_913041895.1 | 65.3 | Verrucomicrobiota | Verrucomicrobiae | Limisphaerales | UBA1100 | UBA1100 | UBA1100 sp913041895 | 0.738 | 0.000 | 0.004 | Cytoplasm | M0 |
| gene_476810\|GeneMark.hmm\|113_aa\|-\|2\|340_271254 | RS_GCF_000935215.1 | 100.0 | Pseudomonadota | Gammaproteobacteria | Pseudomonadales | Pseudomonadaceae | Stutzerimonas | Stutzerimonas stutzeri_AD | 0.020 | 0.994 | 0.000 | Periplasm | M0 |
| gene_479614\|GeneMark.hmm\|321_aa\|+\|3035\|4000_250156 | GB_GCA_913045265.1 | 99.4 | Pseudomonadota | Gammaproteobacteria | Pseudomonadales | Azotimanducaceae_A | UBA11889 | UBA11889 sp002313255 | 0.005 | 0.138 | 0.860 | Extracellular | M0 |
| gene_495017\|GeneMark.hmm\|140_aa\|-\|1\|420_272339 | GB_GCA_029940535.1 | 69.3 | Acidobacteriota | Vicinamibacteria | Vicinamibacterales | UBA8438 | UBA8438 | UBA8438 sp029940535 | 0.350 | 0.003 | 0.044 | Cytoplasm | M0 |
| gene_499232\|GeneMark.hmm\|338_aa\|-\|1646\|2662_259897 | GB_GCA_021741155.1 | 63.3 | Bacteroidota | Bacteroidia | UBA10329 | UBA10329 | UBA10329 | UBA10329 sp021741155 | 0.273 | 0.870 | 0.058 | Periplasm | M0 |
| gene_502192\|GeneMark.hmm\|272_aa\|+\|359\|1177_253891 | GB_GCA_002731735.1 | 99.3 | Gemmatimonadota | Gemmatimonadetes | Longimicrobiales | UBA6960 | UBA822 | UBA822 sp002731735 | 0.056 | 0.049 | 0.739 | Extracellular | M0 |
| gene_528133\|GeneMark.hmm\|416_aa\|+\|108\|1358_300246 | GB_GCA_913057795.1 | 96.5 | Actinomycetota | Actinomycetes | Actinomycetales | Microbacteriaceae | Aquiluna | Aquiluna sp913057795 | 0.983 | 0.000 | 0.000 | Cytoplasm | M0 |
| gene_547592\|GeneMark.hmm\|143_aa\|-\|215\|646_292190 | GB_GCA_937888665.1 | 87.3 | Acidobacteriota | Vicinamibacteria | Vicinamibacterales | UBA8438 | UBA8438 | UBA8438 sp937888665 | 0.199 | 0.109 | 0.592 | Extracellular | M0 |
| gene_566171\|GeneMark.hmm\|262_aa\|+\|3251\|4039_285907 | GB_GCA_016716275.1 | 77.5 | Pseudomonadota | Gammaproteobacteria | Burkholderiales | Burkholderiaceae | JADJWR01 | JADJWR01 sp016716275 | 0.976 | 0.000 | 0.000 | Cytoplasm | M0 |
| gene_569849\|GeneMark.hmm\|414_aa\|-\|13009\|14253_318220 | GB_GCA_902727945.1 | 64.7 | Pseudomonadota | Gammaproteobacteria | Thiotrichales | Thiotrichaceae | Thiolinea | Thiolinea sp902727945 | 0.985 | 0.000 | 0.000 | Cytoplasm | M0 |
| gene_586071\|GeneMark.hmm\|448_aa\|-\|101\|1447_287259 | GB_GCA_018401455.1 | 69.1 | Bacteroidota | Bacteroidia | Flavobacteriales | Parvicellaceae | Parvicella | Parvicella sp018401455 | 0.004 | 1.000 | 0.000 | Periplasm | M0 |
| gene_589840\|GeneMark.hmm\|279_aa\|+\|1578\|2417_299208 | GB_GCA_001438145.1 | 98.2 | Pseudomonadota | Gammaproteobacteria | Pseudomonadales | Pseudohongiellaceae | UBA9145 | UBA9145 sp001438145 | 0.006 | 0.002 | 0.988 | Extracellular | M0 |
| gene_600874\|GeneMark.hmm\|174_aa\|+\|1\|525_292980 | GB_GCA_913041895.1 | 90.9 | Verrucomicrobiota | Verrucomicrobiae | Limisphaerales | UBA1100 | UBA1100 | UBA1100 sp913041895 | 0.459 | 0.000 | 0.003 | Cytoplasm | M0 |
| gene_618362\|GeneMark.hmm\|241_aa\|-\|405\|1130_318093 | GB_GCA_913041895.1 | 85.8 | Verrucomicrobiota | Verrucomicrobiae | Limisphaerales | UBA1100 | UBA1100 | UBA1100 sp913041895 | 0.009 | 0.043 | 0.956 | Extracellular | M0 |
| gene_622560\|GeneMark.hmm\|204_aa\|-\|1\|612_317731 | GB_GCA_030747505.1 | 76.4 | Verrucomicrobiota | Verrucomicrobiae | Limisphaerales | UBA1100 | UBA1100 | UBA1100 sp030747505 | 0.538 | 0.003 | 0.016 | Cytoplasm | M0 |
| gene_634633\|GeneMark.hmm\|354_aa\|-\|16\|1080_310533 | RS_GCF_009796785.1 | 89.7 | Bacteroidota | Bacteroidia | Flavobacteriales | Flavobacteriaceae | Polaribacter | Polaribacter sp009796785 | 0.967 | 0.000 | 0.000 | Cytoplasm | M0 |
| gene_652870\|GeneMark.hmm\|181_aa\|+\|3\|548_328898 | GB_GCA_030747505.1 | 63.6 | Verrucomicrobiota | Verrucomicrobiae | Limisphaerales | UBA1100 | UBA1100 | UBA1100 sp030747505 | 0.249 | 0.000 | 0.027 | Cytoplasm | M0 |
| gene_654691\|GeneMark.hmm\|265_aa\|-\|1045\|1842_362295 | GB_GCA_913061515.1 | 80.8 | Pseudomonadota | Gammaproteobacteria | Pseudomonadales | Pseudohongiellaceae | UBA5078 | UBA5078 sp913061515 | 0.007 | 0.008 | 0.989 | Extracellular | M0 |
| gene_67252\|GeneMark.hmm\|274_aa\|+\|60201\|61025_34097 | GB_GCA_012960305.1 | 99.6 | Pseudomonadota | Gammaproteobacteria | Pseudomonadales | Pseudohongiellaceae | UBA9145 | UBA9145 sp012960305 | 0.004 | 0.004 | 0.996 | Extracellular | M0 |
| gene_685487\|GeneMark.hmm\|334_aa\|+\|13504\|14508_346449 | GB_GCA_949372665.1 | 93.1 | Verrucomicrobiota | Verrucomicrobiae | Verrucomicrobiales | Akkermansiaceae | SW10 | SW10 sp905181965 | 0.004 | 0.005 | 0.995 | Extracellular | M0 |

| | | | | | | | | | | | | | |
|---|---|---|---|---|---|---|---|---|---|---|---|---|---|
| gene_696814\|GeneMark.hmm\|142_aa\|-\|35\|463_365790 | GB_GCA_949487135.1 | 66.4 | Pseudomonadota | Gammaproteobacteria | Pseudomonadales | Porticoccaceae | HTCC2207 | HTCC2207 sp949487135 | 0.296 | 0.001 | 0.499 | Extracellular | M0 |
| gene_704864\|GeneMark.hmm\|405_aa\|-\|26803\|28020_362019 | GB_GCA_003045865.1 | 85.4 | Bacteroidota | Bacteroidia | Flavobacteriales | Schleiferiaceae | Croceimicrobium | Croceimicrobium sp003045865 | 0.979 | 0.000 | 0.000 | Cytoplasm | M0 |
| gene_70635\|GeneMark.hmm\|281_aa\|+\|966\|1811_38678 | RS_GCF_021495345.1 | 91.0 | Pseudomonadota | Gammaproteobacteria | Pseudomonadales | Pseudomonadaceae | Pseudomonas_E | Pseudomonas_E hydrolytica | 0.005 | 0.909 | 0.090 | Periplasm | M0 |
| gene_706699\|GeneMark.hmm\|161_aa\|+\|1\|486_365664 | GB_GCA_002708465.1 | 91.9 | Planctomycetota | UBA8108 | UBA8108 | UBA8108 | UBA8108 | UBA8108 sp002708465 | 0.752 | 0.000 | 0.005 | Cytoplasm | M0 |
| gene_707045\|GeneMark.hmm\|519_aa\|-\|1\|1557_406346 | GB_GCA_913058215.1 | 79.2 | Verrucomicrobiota | Verrucomicrobiae | Verrucomicrobiales | DEV007 | Arctic95D-9 | Arctic95D-9 sp913058215 | 0.004 | 0.000 | 0.999 | Extracellular | M0 |
| gene_707782\|GeneMark.hmm\|270_aa\|-\|46\|858_366505 | GB_GCA_029863645.1 | 63.9 | Gemmatimonadota | Gemmatimonadetes | Longimicrobiales | UBA6960 | UBA2982 | UBA2982 sp029863645 | 0.005 | 0.364 | 0.636 | Extracellular | M0 |
| gene_714537\|GeneMark.hmm\|280_aa\|+\|586261\|587103_371923 | GB_GCA_913058135.1 | 86.4 | Pseudomonadota | Gammaproteobacteria | Pseudomonadales | Pseudohongiellaceae | Pseudohongiella | Pseudohongiella sp913058135 | 0.005 | 0.553 | 0.446 | Periplasm | M0 |
| gene_74001\|GeneMark.hmm\|471_aa\|+\|630\|2045_37295 | GB_GCA_000961845.1 | 96.2 | Bacteroidota | Bacteroidia | Flavobacteriales | Vicingaceae | BRH-c54 | BRH-c54 sp000961845 | 0.006 | 1.000 | 0.000 | Periplasm | M0 |
| gene_758694\|GeneMark.hmm\|258_aa\|-\|54\|830_372875 | GB_GCA_002721705.1 | 59.0 | Verrucomicrobiota | Verrucomicrobiae | Opitutales | MB11C04 | UBA2993 | UBA2993 sp002721705 | 0.006 | 0.006 | 0.987 | Extracellular | M0 |
| gene_764761\|GeneMark.hmm\|306_aa\|+\|1\|921_432735 | GB_GCA_030731175.1 | 81.4 | Bacteroidota | Bacteroidia | Cytophagales | Spirosomataceae | JANQUM01 | JANQUM01 sp030731175 | 0.888 | 0.000 | 0.000 | Cytoplasm | M0 |
| gene_767267\|GeneMark.hmm\|343_aa\|-\|425\|1456_426799 | RS_GCF_900115565.1 | 53.7 | Actinomycetota | Actinomycetes | Mycobacteriales | Pseudonocardiaceae | Yuhushiella | Yuhushiella arida | 0.009 | 0.919 | 0.063 | Periplasm | M0 |
| gene_771387\|GeneMark.hmm\|400_aa\|-\|2334\|3536_375433 | RS_GCF_014905615.1 | 99.8 | Pseudomonadota | Gammaproteobacteria | Enterobacterales | Alteromonadaceae | Pseudoalteromonas | Pseudoalteromonas aliena | 0.959 | 0.000 | 0.000 | Cytoplasm | M0 |
| gene_787535\|GeneMark.hmm\|145_aa\|-\|60\|497_400902 | GB_GCA_002721705.1 | 73.2 | Verrucomicrobiota | Verrucomicrobiae | Opitutales | MB11C04 | UBA2993 | UBA2993 sp002721705 | 0.446 | 0.000 | 0.006 | Cytoplasm | M0 |
| gene_79821\|GeneMark.hmm\|528_aa\|-\|31306\|32892_42157 | GB_GCA_022448945.1 | 46.7 | Bacteroidota | Bacteroidia | Chitinophagales | Saprospiraceae | Aureispira | Aureispira sp022448945 | 0.004 | 1.000 | 0.000 | Periplasm | M0 |
| gene_820300\|GeneMark.hmm\|179_aa\|-\|72\|611_414572 | GB_GCA_009692065.1 | 66.2 | Planctomycetota | Planctomycetia | Gemmatales | Gemmataceae | SIAQ01 | SIAQ01 sp009692065 | 0.814 | 0.000 | 0.020 | Cytoplasm | M0 |
| gene_826957\|GeneMark.hmm\|339_aa\|+\|3\|1022_375770 | GB_GCA_905181585.1 | 86.8 | Bacteroidota | Bacteroidia | Flavobacteriales | BACL11 | CAJIYG01 | CAJIYG01 sp905181585 | 0.004 | 0.982 | 0.017 | Periplasm | M0 |
| gene_833796\|GeneMark.hmm\|294_aa\|+\|5040\|5924_414627 | RS_GCF_950066645.1 | 77.8 | Pseudomonadota | Gammaproteobacteria | Pseudomonadales | DSM-6294 | Oleibacter | Oleibacter sp950066645 | 0.005 | 0.020 | 0.980 | Extracellular | M0 |
| gene_844270\|GeneMark.hmm\|293_aa\|-\|1100\|1981_419449 | GB_GCA_016763285.1 | 100.0 | Pseudomonadota | Gammaproteobacteria | Pseudomonadales | Pseudomonadaceae | Halopseudomonas | Halopseudomonas sp016763285 | 0.004 | 0.996 | 0.004 | Periplasm | M0 |
| gene_848863\|GeneMark.hmm\|263_aa\|-\|1034\|1825_438410 | GB_GCA_002731775.1 | 100.0 | Pseudomonadota | Gammaproteobacteria | Pseudomonadales | Pseudohongiellaceae | UBA9145 | UBA9145 sp002731775 | 0.005 | 0.007 | 0.991 | Extracellular | M0 |

| | | | | | | | | | | | | | |
|---|---|---|---|---|---|---|---|---|---|---|---|---|---|
| gene_851835\|GeneMark.hmm\|400_aa\|-\|8268\|9470_442134 | GB_GCA_913061435.1 | 86.5 | Pseudomonadota | Gammaproteobacteria | Pseudomonadales | DSM-6294 | Oleispira | Oleispira sp002632185 | 0.955 | 0.000 | 0.000 | Cytoplasm | M0 |
| gene_858637\|GeneMark.hmm\|396_aa\|+\|67945\|69135_472616 | GB_GCA_021776025.1 | 76.1 | Pseudomonadota | Alphaproteobacteria | Micavibrionales | Micavibrionaceae | TMED27 | TMED27 sp021776025 | 0.955 | 0.000 | 0.000 | Cytoplasm | M0 |
| gene_863938\|GeneMark.hmm\|398_aa\|+\|1\|1197_420713 | GB_GCA_905182175.1 | 89.4 | Bacteroidota | Bacteroidia | Flavobacteriales | Vicingaceae | NORP294 | NORP294 sp905182175 | 0.972 | 0.000 | 0.000 | Cytoplasm | M0 |
| gene_869587\|GeneMark.hmm\|258_aa\|-\|2\|775_490072 | GB_GCA_913057795.1 | 97.7 | Actinomycetota | Actinomycetes | Actinomycetales | Microbacteriaceae | Aquiluna | Aquiluna sp913057795 | 0.981 | 0.000 | 0.000 | Cytoplasm | M0 |
| gene_872455\|GeneMark.hmm\|291_aa\|-\|103\|978_442796 | GB_GCA_002699725.1 | 97.3 | Actinomycetota | Acidimicrobiia | Acidimicrobiales | MedAcidi-G1 | S20-B6 | S20-B6 sp002699725 | 0.037 | 0.010 | 0.563 | Extracellular | M0 |
| gene_881306\|GeneMark.hmm\|156_aa\|-\|3\|470_438135 | GB_GCA_002705605.1 | 68.6 | Marinisomatota | Marinisomatia | SCGC-AAA003-L08 | GCA-002707645 | GCA-002707645 | GCA-002707645 sp002705605 | 0.167 | 0.008 | 0.015 | Cytoplasm | M0 |
| gene_920204\|GeneMark.hmm\|192_aa\|-\|1\|576_447317 | RS_GCF_019222765.1 | 51.8 | Actinomycetota | Actinomycetes | Mycobacteriales | Mycobacteriaceae | Nocardia | Nocardia iowensis | 0.406 | 0.452 | 0.000 | Periplasm | M0 |
| gene_937543\|GeneMark.hmm\|210_aa\|-\|2\|631_515685 | GB_GCA_018829385.1 | 81.4 | Pseudomonadota | Alphaproteobacteria | Rhizobiales | Rhizobiaceae | Hoeflea | Hoeflea sp018829385 | 0.982 | 0.000 | 0.000 | Cytoplasm | M0 |
| gene_940510\|GeneMark.hmm\|269_aa\|+\|167\|976_488213 | GB_GCA_028825335.1 | 84.0 | Pseudomonadota | Gammaproteobacteria | Pseudomonadales | Pseudohongiellaceae | UBA9145 | UBA9145 sp028825335 | 0.007 | 0.001 | 0.994 | Extracellular | M0 |
| gene_94857\|GeneMark.hmm\|402_aa\|+\|2\|1210_50381 | GB_GCA_030747505.1 | 89.3 | Verrucomicrobiota | Verrucomicrobiae | Limisphaerales | UBA1100 | UBA1100 | UBA1100 sp030747505 | 0.750 | 0.000 | 0.012 | Cytoplasm | M0 |
| gene_951958\|GeneMark.hmm\|121_aa\|-\|1\|363_523525 | GB_GCA_020635795.1 | 100.0 | Bacteroidota | Bacteroidia | Flavobacteriales | Vicingaceae | Vicingus | Vicingus sp020635795 | 0.607 | 0.151 | 0.046 | Cytoplasm | M0 |
| gene_97796\|GeneMark.hmm\|355_aa\|-\|3256\|4323_50967 | GB_GCA_018222765.1 | 95.2 | Verrucomicrobiota | Verrucomicrobiae | Verrucomicrobiales | Akkermansiaceae | SW10 | SW10 sp018222765 | 0.012 | 0.006 | 0.952 | Extracellular | M0 |
| gene_995120\|GeneMark.hmm\|388_aa\|-\|2\|1165_509923 | GB_GCA_002471805.1 | 80.1 | Marinisomatota | Marinisomatia | Marinisomatales | UBA1611 | UBA2134 | UBA2134 sp002471805 | 0.005 | 0.795 | 0.205 | Periplasm | M0 |
| gene_995183\|GeneMark.hmm\|164_aa\|+\|60\|554_498082 | GB_GCA_029941275.1 | 74.4 | Gemmatimonadota | Gemmatimonadetes | Longimicrobiales | UBA6960 | UBA822 | UBA822 sp029941275 | 0.484 | 0.000 | 0.020 | Cytoplasm | M0 |
| gene_999805\|GeneMark.hmm\|285_aa\|-\|180\|1037_549520 | GB_GCA_015664885.1 | 98.6 | Gemmatimonadota | Gemmatimonadetes | Longimicrobiales | UBA6960 | UBA1138 | UBA1138 sp003447875 | 0.013 | 0.075 | 0.923 | Extracellular | M0 |
| NODE_111145_length_1055_cov_2.696_1 | RS_GCF_001678675.1 | 91.7 | Bacteroidota | Bacteroidia | Flavobacteriales | Flavobacteriaceae | Gelidibacter | Gelidibacter algens | 0.980 | 0.000 | 0.000 | Cytoplasm | M0 |
| NODE_130825_length_945_cov_3.49775_1 | RS_GCF_002364335.1 | 92.2 | Bacteroidota | Bacteroidia | Flavobacteriales | Flavobacteriaceae | Winogradskyella | Winogradskyella sp002364335 | 0.016 | 0.984 | 0.000 | Periplasm | M0 |
| NODE_142_length_39438_cov_17.053627_6 | GB_GCA_000981645.1 | 80.1 | Bacteroidota | Bacteroidia | Flavobacteriales | UA16 | ASP10-05a | ASP10-05a sp000981645 | 0.059 | 0.999 | 0.000 | Periplasm | M0 |
| NODE_18431_length_3542_cov_16.1566_2 | RS_GCF_029238425.1 | 69.2 | Bacteroidota | Bacteroidia | Flavobacteriales | Flavobacteriaceae | Flavicella | Flavicella sp029238425 | 0.964 | 0.000 | 0.001 | Cytoplasm | M0 |
| NODE_1850_length_14933_cov_11.3174_2 | GB_GCA_011053055.1 | 99.6 | Pseudomonadota | Gammaproteobacteria | Pseudomonadales | Pseudomonadaceae | Halopseudomonas | Halopseudomonas sabulinigri_A | 0.370 | 0.909 | 0.000 | Periplasm | M0 |

| | | | | | | | | | | | | | |
|---|---|---|---|---|---|---|---|---|---|---|---|---|---|
| NODE_19222_length_2634_cov_2.400155_2 | RS_GCF_014636175.1 | 96.8 | Pseudomonadota | Alphaproteobacteria | Sphingomonadales | Sphingomonadaceae | Sphingomonas | Sphingomonas psychrolutea | 0.983 | 0.000 | 0.000 | Cytoplasm | M0 |
| NODE_21048_length_3245_cov_14.3122_4 | GB_GCA_905480405.1 | 95.4 | Bacteroidota | Bacteroidia | Flavobacteriales | Flavobacteriaceae | Polaribacter | Polaribacter sp905480405 | 0.972 | 0.000 | 0.000 | Cytoplasm | M0 |
| NODE_2181_length_10778_cov_8.024900_5 | RS_GCF_017948265.1 | 89.2 | Pseudomonadota | Gammaproteobacteria | Burkholderiales | Burkholderiaceae | Rhodoferax | Rhodoferax sp017948265 | 0.980 | 0.000 | 0.000 | Cytoplasm | M0 |
| NODE_219_length_48746_cov_15.4281_30 | GB_GCA_905182175.1 | 93.6 | Bacteroidota | Bacteroidia | Flavobacteriales | Vicingaceae | NORP294 | NORP294 sp905182175 | 0.972 | 0.000 | 0.000 | Cytoplasm | M0 |
| NODE_276275_length_586_cov_1.80979_1 | RS_GCF_008641105.1 | 87.1 | Pseudomonadota | Gammaproteobacteria | Pseudomonadales | Pseudomonadaceae | Halopseudomonas | Halopseudomonas salina | 0.575 | 0.000 | 0.009 | Cytoplasm | M0 |
| NODE_28437_length_2040_cov_2.286650_3 | GB_GCA_019638845.1 | 84.0 | Actinomycetota | Actinomycetes | Actinomycetales | Microbacteriaceae | Rhodoglobus | Rhodoglobus sp019638845 | 0.977 | 0.000 | 0.000 | Cytoplasm | M0 |
| NODE_3365_length_4932_cov_5.332787_3 | GB_GCA_943788405.1 | 88.1 | Bacteroidota | Bacteroidia | Flavobacteriales | BACL11 | UWMA-0277 | UWMA-0277 sp943788405 | 0.015 | 0.999 | 0.000 | Periplasm | M0 |
| NODE_34054_length_1174_cov_2.913315_1 | GB_GCA_913031035.1 | 98.4 | Pseudomonadota | Gammaproteobacteria | Pseudomonadales | Pseudohongiellaceae | UBA9145 | UBA9145 sp913031035 | 0.974 | 0.000 | 0.000 | Cytoplasm | M0 |
| NODE_35840_length_2274_cov_5.21361_1 | RS_GCF_001650875.1 | 99.3 | Pseudomonadota | Alphaproteobacteria | Rhodobacterales | Rhodobacteraceae | Sulfitobacter | Sulfitobacter sp001650875 | 0.985 | 0.000 | 0.000 | Cytoplasm | M0 |
| NODE_513_length_10408_cov_18.2396_7 | RS_GCF_000425505.1 | 98.0 | Bacteroidota | Bacteroidia | Flavobacteriales | Flavobacteriaceae | Flavobacterium | Flavobacterium frigidarium | 0.959 | 0.000 | 0.000 | Cytoplasm | M0 |
| NODE_54646_length_1710_cov_4.69245_1 | RS_GCF_001418015.1 | 80.9 | Bacteroidota | Bacteroidia | Flavobacteriales | Flavobacteriaceae | Lacinutrix | Lacinutrix mariniflava | 0.955 | 0.000 | 0.000 | Cytoplasm | M0 |
| NODE_54699_length_902_cov_1.639906_1 | GB_GCA_913031035.1 | 97.0 | Pseudomonadota | Gammaproteobacteria | Pseudomonadales | Pseudohongiellaceae | UBA9145 | UBA9145 sp913031035 | 0.918 | 0.000 | 0.141 | Cytoplasm | M0 |
| NODE_5596_length_7718_cov_9.28957_6 | GB_GCA_028825475.1 | 100.0 | Pseudomonadota | Alphaproteobacteria | Rhodobacterales | Rhodobacteraceae | Ascidiaceihabitans | Ascidiaceihabitans sp028825475 | 0.982 | 0.000 | 0.000 | Cytoplasm | M0 |
| NODE_59288_length_1618_cov_4.66795_1 | RS_GCF_008641105.1 | 99.6 | Pseudomonadota | Gammaproteobacteria | Pseudomonadales | Pseudomonadaceae | Halopseudomonas | Halopseudomonas salina | 0.431 | 0.915 | 0.000 | Periplasm | M0 |
| NODE_61763_length_844_cov_2.671736_1 | GB_GCA_949372635.1 | 91.1 | Pseudomonadota | Gammaproteobacteria | Pseudomonadales | Pseudohongiellaceae | UBA9145 | UBA9145 sp949372635 | 0.942 | 0.000 | 0.011 | Cytoplasm | M0 |
| NODE_65999_length_1504_cov_4.89027_1 | RS_GCF_002934445.1 | 92.9 | Bacteroidota | Bacteroidia | Flavobacteriales | Flavobacteriaceae | Nonlabens | Nonlabens xylanidelens | 0.981 | 0.000 | 0.000 | Cytoplasm | M0 |
| NODE_6901_length_6763_cov_14.4232_3 | GB_GCA_905182175.1 | 83.8 | Bacteroidota | Bacteroidia | Flavobacteriales | Vicingaceae | NORP294 | NORP294 sp905182175 | 0.113 | 0.966 | 0.012 | Periplasm | M0 |
| NODE_703_length_26339_cov_26.6636_28 | RS_GCF_900114485.1 | 99.3 | Pseudomonadota | Alphaproteobacteria | Rhodobacterales | Rhodobacteraceae | Loktanella | Loktanella salsilacus | 0.979 | 0.000 | 0.000 | Cytoplasm | M0 |
| NODE_7399_length_6475_cov_8.66153_1 | GB_GCA_000967895.1 | 100.0 | Pseudomonadota | Gammaproteobacteria | Pseudomonadales | DSM-6294 | Oleispira | Oleispira antarctica | 0.972 | 0.000 | 0.000 | Cytoplasm | M0 |
| NODE_7455_length_6444_cov_5.71107_2 | RS_GCF_008641105.1 | 97.2 | Pseudomonadota | Gammaproteobacteria | Pseudomonadales | Pseudomonadaceae | Halopseudomonas | Halopseudomonas salina | 0.967 | 0.000 | 0.000 | Cytoplasm | M0 |

| | | | | | | | | | | | | | |
|---|---|---|---|---|---|---|---|---|---|---|---|---|---|
| NODE_78629_length_648_cov_2.217538_1 | GB_GCA_938075455.1 | 80.5 | Bacteroidota | Bacteroidia | Flavobacteriales | Parvicellaceae | MED-G20 | MED-G20 sp938075455 | 0.105 | 0.445 | 0.000 | Periplasm | M0 |
| NODE_828_length_23525_cov_9.01623_8 | RS_GCF_003075045.1 | 95.3 | Bacteroidota | Bacteroidia | Flavobacteriales | Flavobacteriaceae | Maribacter | Maribacter litoralis | 0.143 | 0.004 | 0.688 | Extracellular | M0 |
| NODE_841_length_23354_cov_16.6602_5 | RS_GCF_002221505.1 | 100.0 | Pseudomonadota | Gammaproteobacteria | Enterobacterales | Alteromonadaceae | Pseudoalteromonas | Pseudoalteromonas nigrifaciens | 0.924 | 0.000 | 0.000 | Cytoplasm | M0 |
| NODE_890_length_22621_cov_10.8946_9 | GB_GCA_913061335.1 | 94.7 | Pseudomonadota | Alphaproteobacteria | Rhodobacterales | Rhodobacteraceae | Yoonia | Yoonia sp913061335 | 0.985 | 0.000 | 0.000 | Cytoplasm | M0 |
| OM-RGC.v1.000148874 | GB_GCA_004321745.1 | 69.5 | Marinisomatota | Marinisomatia | Marinisomatales | UBA1611 | TMED80 | TMED80 sp002170405 | 0.004 | 1.000 | 0.000 | Periplasm | M0 |
| OM-RGC.v1.000590104 | GB_GCA_018651745.1 | 61.2 | Marinisomatota | Marinisomatia | Marinisomatales | TCS55 | JAAZYX01 | JAAZYX01 sp018651745 | 0.006 | 1.000 | 0.000 | Periplasm | M0 |
| OM-RGC.v1.001308623 | GB_GCA_002721705.1 | 76.7 | Verrucomicrobiota | Verrucomicrobiae | Opitutales | MB11C04 | UBA2993 | UBA2993 sp002721705 | 0.004 | 0.012 | 0.985 | Extracellular | M0 |
| OM-RGC.v1.001522890 | GB_GCA_002724245.1 | 100.0 | Planctomycetota | Planctomycetia | Pirellulales | Pirellulaceae | UBA9264 | UBA9264 sp002724245 | 0.005 | 0.027 | 0.971 | Extracellular | M0 |
| OM-RGC.v1.002264619 | GB_GCA_913041775.1 | 52.4 | Bacteroidota | Bacteroidia | Flavobacteriales | UBA7430 | UBA6772 | UBA6772 sp913041775 | 0.004 | 0.999 | 0.001 | Periplasm | M0 |
| OM-RGC.v1.003146980 | GB_GCA_002500505.1 | 100.0 | Planctomycetota | UBA1135 | UBA1135 | UBA1135 | UBA7888 | UBA7888 sp002500505 | 0.004 | 0.308 | 0.692 | Extracellular | M0 |
| OM-RGC.v1.003147666 | GB_GCA_018402085.1 | 84.4 | Bacteroidota | Bacteroidia | Flavobacteriales | UA16 | M30B41 | M30B41 sp018402085 | 0.004 | 0.427 | 0.573 | Extracellular | M0 |
| OM-RGC.v1.003733882 | GB_GCA_913047475.1 | 74.2 | Marinisomatota | Marinisomatia | SCGC-AAA003-L08 | TMED6 | TMED6 | TMED6 sp913047475 | 0.004 | 1.000 | 0.000 | Periplasm | M0 |
| OM-RGC.v1.003915512 | GB_GCA_002704045.1 | 92.3 | Marinisomatota | Marinisomatia | SCGC-AAA003-L08 | GCA-002704045 | GCA-002704045 | GCA-002704045 sp002704045 | 0.004 | 0.999 | 0.001 | Periplasm | M0 |
| OM-RGC.v1.003922963 | GB_GCA_029941685.1 | 64.5 | Planctomycetota | UBA1135 | UBA1135 | UBA1135 | JAPKEZ01 | JAPKEZ01 sp029941685 | 0.087 | 0.999 | 0.000 | Periplasm | M0 |
| OM-RGC.v1.003925493 | GB_GCA_913051865.1 | 100.0 | Bacteroidota | Bacteroidia | Flavobacteriales | UA16 | UBA974 | UBA974 sp002336655 | 0.005 | 0.583 | 0.417 | Periplasm | M0 |
| OM-RGC.v1.003930091 | GB_GCA_002689995.1 | 65.5 | Marinisomatota | Marinisomatia | SCGC-AAA003-L08 | GCA-2689995 | GCA-2689995 | GCA-2689995 sp002689995 | 0.004 | 1.000 | 0.000 | Periplasm | M0 |
| OM-RGC.v1.003986021 | GB_GCA_002731795.1 | 99.2 | Marinisomatota | Marinisomatia | SCGC-AAA003-L08 | TMED161 | TMED161 | TMED161 sp002731795 | 0.004 | 1.000 | 0.000 | Periplasm | M0 |
| OM-RGC.v1.004008314 | GB_GCA_014237975.1 | 62.0 | Planctomycetota | Planctomycetia | Pirellulales | Pirellulaceae | Mariniblastus | Mariniblastus sp014237975 | 0.822 | 0.000 | 0.012 | Cytoplasm | M0 |
| OM-RGC.v1.004441477 | GB_GCA_002453265.1 | 53.4 | Bacteroidota | Bacteroidia | Flavobacteriales | UBA7430 | UBA6772 | UBA6772 sp002453265 | 0.019 | 0.999 | 0.000 | Periplasm | M0 |

| | | | | | | | | | | | | | |
|---|---|---|---|---|---|---|---|---|---|---|---|---|---|
| OM-RGC.v1.004502153 | GB_GCA_022448765.1 | 37.6 | Pseudomonadota | Gammaproteobacteria | Pseudomonadales | Azotimanducaceae_A | UBA11889 | UBA11889 sp022448765 | 0.137 | 0.002 | 0.021 | Cytoplasm | M0 |
| OM-RGC.v1.004765404 | GB_GCA_018651745.1 | 63.6 | Marinisomatota | Marinisomatia | Marinisomatales | TCS55 | JAAZYX01 | JAAZYX01 sp018651745 | 0.023 | 0.607 | 0.387 | Periplasm | M0 |
| OM-RGC.v1.005164687 | GB_GCA_913045265.1 | 43.1 | Pseudomonadota | Gammaproteobacteria | Pseudomonadales | Azotimanducaceae_A | UBA11889 | UBA11889 sp002313255 | 0.251 | 0.001 | 0.048 | Cytoplasm | M0 |
| OM-RGC.v1.005469377 | GB_GCA_002471805.1 | 85.2 | Marinisomatota | Marinisomatia | Marinisomatales | UBA1611 | UBA2134 | UBA2134 sp002471805 | 0.004 | 0.995 | 0.005 | Periplasm | M0 |
| OM-RGC.v1.005608287 | GB_GCA_002471805.1 | 77.7 | Marinisomatota | Marinisomatia | Marinisomatales | UBA1611 | UBA2134 | UBA2134 sp002471805 | 0.005 | 0.985 | 0.015 | Periplasm | M0 |
| OM-RGC.v1.005734125 | GB_GCA_018651745.1 | 52.8 | Marinisomatota | Marinisomatia | Marinisomatales | TCS55 | JAAZYX01 | JAAZYX01 sp018651745 | 0.004 | 0.377 | 0.622 | Extracellular | M0 |
| OM-RGC.v1.005837034 | GB_GCA_022448085.1 | 96.6 | Pseudomonadota | Gammaproteobacteria | Pseudomonadales | Azotimanducaceae_A | UBA11889 | UBA11889 sp022448085 | 0.007 | 0.145 | 0.852 | Extracellular | M0 |
| OM-RGC.v1.006347060 | GB_GCA_913030615.1 | 81.6 | Pseudomonadota | Gammaproteobacteria | Pseudomonadales | Azotimanducaceae_A | UBA11889 | UBA11889 sp913030615 | 0.005 | 0.003 | 0.996 | Extracellular | M0 |
| OM-RGC.v1.006456638 | GB_GCA_002471665.1 | 100.0 | Pseudomonadota | Gammaproteobacteria | Pseudomonadales | Ketobacteraceae | Ketobacter | Ketobacter sp002471665 | 0.004 | 0.889 | 0.111 | Periplasm | M0 |
| OM-RGC.v1.006605551 | RS_GCF_003444685.1 | 100.0 | Pseudomonadota | Gammaproteobacteria | Pseudomonadales | Pseudomonadaceae | Halopseudomonas | Halopseudomonas gallaeciensis | 0.004 | 1.000 | 0.000 | Periplasm | M0 |
| OM-RGC.v1.006725628 | GB_GCA_002721705.1 | 76.1 | Verrucomicrobiota | Verrucomicrobiae | Opitutales | MB11C04 | UBA2993 | UBA2993 sp002721705 | 0.386 | 0.001 | 0.011 | Cytoplasm | M0 |
| OM-RGC.v1.007193723 | GB_GCA_002731795.1 | 87.6 | Marinisomatota | Marinisomatia | SCGC-AAA003-L08 | TMED161 | TMED161 | TMED161 sp002731795 | 0.006 | 1.000 | 0.000 | Periplasm | M0 |
| OM-RGC.v1.007235507 | GB_GCA_002298335.1 | 100.0 | Pseudomonadota | Gammaproteobacteria | Pseudomonadales | DSM-6294 | Oleibacter | Oleibacter sp002298335 | 0.006 | 0.098 | 0.901 | Extracellular | M0 |
| OM-RGC.v1.007298884 | RS_GCF_913019975.1 | 99.0 | Pseudomonadota | Gammaproteobacteria | Pseudomonadales | DSM-6294 | Oleibacter | Oleibacter sp002331105 | 0.004 | 0.416 | 0.584 | Extracellular | M0 |
| OM-RGC.v1.007606929 | GB_GCA_002724925.1 | 100.0 | Pseudomonadota | Gammaproteobacteria | Pseudomonadales | DSM-6294 | Oleibacter | Oleibacter sp002724925 | 0.006 | 0.005 | 0.993 | Extracellular | M0 |
| OM-RGC.v1.008027965 | GB_GCA_002706685.1 | 100.0 | Pseudomonadota | Gammaproteobacteria | Pseudomonadales | Pseudohongiellaceae | Pseudohongiella | Pseudohongiella sp002706685 | 0.004 | 0.020 | 0.980 | Extracellular | M0 |
| OM-RGC.v1.008076071 | GB_GCA_913030615.1 | 41.9 | Pseudomonadota | Gammaproteobacteria | Pseudomonadales | Azotimanducaceae_A | UBA11889 | UBA11889 sp913030615 | 0.232 | 0.013 | 0.348 | Extracellular | M0 |
| OM-RGC.v1.008213585 | RS_GCF_000219605.1 | 100.0 | Pseudomonadota | Gammaproteobacteria | Pseudomonadales | Pseudomonadaceae | Stutzerimonas | Stutzerimonas stutzeri | 0.004 | 0.868 | 0.132 | Periplasm | M0 |
| OM-RGC.v1.009081041 | GB_GCA_002171125.1 | 90.9 | Marinisomatota | Marinisomatia | SCGC-AAA003-L08 | TMED161 | TMED161 | TMED161 sp002171125 | 0.058 | 0.999 | 0.000 | Periplasm | M0 |
| OM-RGC.v1.009176700 | GB_GCA_002704685.1 | 59.9 | Marinisomatota | Marinisomatia | SCGC-AAA003-L08 | GCA-002707645 | GCA-002707645 | GCA-002707645 sp002704685 | 0.004 | 0.989 | 0.010 | Periplasm | M0 |

| | | | | | | | | | | | | | |
|---|---|---|---|---|---|---|---|---|---|---|---|---|---|
| OM-RGC.v1.009512838 | GB_GCA_028110365.1 | 87.1 | Bacteroidota | Bacteroidia | Flavobacteriales | UBA7430 | UBA6772 | UBA6772 sp028110365 | 0.018 | 0.870 | 0.108 | Periplasm | M0 |
| OM-RGC.v1.011041340 | GB_GCA_937995755.1 | 59.9 | Bacteroidota | Bacteroidia | Chitinophagales | Saprospiraceae | UBA1994 | UBA1994 sp937995755 | 0.005 | 0.854 | 0.145 | Periplasm | M0 |
| OM-RGC.v1.011341023 | RS_GCF_003797945.1 | 99.6 | Pseudomonadota | Gammaproteobacteria | Pseudomonadales | Pseudomonadaceae | Halopseudomonas | Halopseudomonas neustonica | 0.005 | 0.999 | 0.001 | Periplasm | M0 |
| OM-RGC.v1.011579526 | GB_GCA_002725955.1 | 100.0 | Bdellovibrionota | Oligoflexia | Oligoflexales | Oligoflexaceae | GCA-2725955 | GCA-2725955 sp002725955 | 0.136 | 0.015 | 0.882 | Extracellular | M0 |
| OM-RGC.v1.013470237 | GB_GCA_002715785.1 | 95.5 | Pseudomonadota | Gammaproteobacteria | Pseudomonadales | Azotimanducaceae_A | UBA11889 | UBA11889 sp002715785 | 0.569 | 0.004 | 0.030 | Cytoplasm | M0 |
| OM-RGC.v1.013716341 | GB_GCA_002453265.1 | 50.0 | Bacteroidota | Bacteroidia | Flavobacteriales | UBA7430 | UBA6772 | UBA6772 sp002453265 | 0.808 | 0.186 | 0.000 | Cytoplasm | M0 |
| OM-RGC.v1.014433907 | GB_GCA_913030615.1 | 99.1 | Pseudomonadota | Gammaproteobacteria | Pseudomonadales | Azotimanducaceae_A | UBA11889 | UBA11889 sp913030615 | 0.961 | 0.000 | 0.000 | Cytoplasm | M0 |
| OM-RGC.v1.015367204 | GB_GCA_002724925.1 | 96.6 | Pseudomonadota | Gammaproteobacteria | Pseudomonadales | DSM-6294 | Oleibacter | Oleibacter sp002724925 | 0.773 | 0.001 | 0.729 | Cytoplasm | M0 |
| OM-RGC.v1.019488603 | RS_GCF_950066645.1 | 100.0 | Pseudomonadota | Gammaproteobacteria | Pseudomonadales | DSM-6294 | Oleibacter | Oleibacter sp950066645 | 0.161 | 0.001 | 0.169 | Extracellular | M0 |
| gene_1236733\|GeneMark.hmm\|117_aa\|-\|2\|352_600506 | GB_GCA_913061205.1 | 87.2 | Bacteroidota | Bacteroidia | Flavobacteriales | Flavobacteriaceae | GCA-002733185 | GCA-002733185 sp913061205 | 0.018 | 0.804 | 0.000 | Periplasm | M3 |
| gene_4973\|GeneMark.hmm\|111_aa\|-\|2\|334_2427 | GB_GCA_009692305.1 | 86.5 | Pseudomonadota | Gammaproteobacteria | Pseudomonadales | Pseudohongiellaceae | UBA9145 | UBA9145 sp009692305 | 0.122 | 0.019 | 0.176 | Extracellular | M3 |
| gene_62801\|GeneMark.hmm\|220_aa\|-\|3\|662_32058 | GB_GCA_028825365.1 | 94.5 | Bacteroidota | Bacteroidia | Flavobacteriales | Flavobacteriaceae | SGZJ01 | SGZJ01 sp028825365 | 0.068 | 0.780 | 0.121 | Periplasm | M3 |
| gene_824001\|GeneMark.hmm\|314_aa\|-\|3\|944_410215 | GB_GCA_002453265.1 | 99.4 | Bacteroidota | Bacteroidia | Flavobacteriales | UBA7430 | UBA6772 | UBA6772 sp002453265 | 0.245 | 0.981 | 0.000 | Periplasm | M3 |
| gene_898832\|GeneMark.hmm\|303_aa\|+\|152\|1063_498169 | RS_GCF_014203755.1 | 38.0 | Actinomycetota | Actinomycetes | Mycobacteriales | Pseudonocardiaceae | Actinosynnema | Actinosynnema violaceirubrum | 0.004 | 0.143 | 0.855 | Extracellular | M3 |
| gene_937290\|GeneMark.hmm\|244_aa\|+\|3\|737_425371 | GB_GCA_028825365.1 | 95.5 | Bacteroidota | Bacteroidia | Flavobacteriales | Flavobacteriaceae | SGZJ01 | SGZJ01 sp028825365 | 0.052 | 0.823 | 0.158 | Periplasm | M3 |
| NODE_78639_length_839_cov_1.920918_1 | GB_GCA_028823435.1 | 79.9 | Bacteroidota | Bacteroidia | Flavobacteriales | UA16 | UBA974 | UBA974 sp028823435 | 0.546 | 0.002 | 0.065 | Cytoplasm | M3 |
| OM-RGC.v1.004122961 | GB_GCA_022425465.1 | 90.8 | Bacteroidota | Bacteroidia | NS11-12g | UBA9320 | UBA9320 | UBA9320 sp022425465 | 0.004 | 0.977 | 0.023 | Periplasm | M3 |
| OM-RGC.v1.006333173 | GB_GCA_002453265.1 | 52.6 | Bacteroidota | Bacteroidia | Flavobacteriales | UBA7430 | UBA6772 | UBA6772 sp002453265 | 0.035 | 0.998 | 0.000 | Periplasm | M3 |
| OM-RGC.v1.008984481 | GB_GCA_913020765.1 | 77.0 | Bacteroidota | Bacteroidia | Flavobacteriales | Crocinitomicaceae | UBA952 | UBA952 sp913020765 | 0.015 | 0.870 | 0.122 | Periplasm | M3 |
| OM-RGC.v1.009392135 | GB_GCA_002683585.1 | 96.2 | Bacteroidota | Bacteroidia | Flavobacteriales | Crocinitomicaceae | UBA952 | UBA952 sp002683585 | 0.067 | 0.483 | 0.501 | Extracellular | M3 |

| | | | | | | | | | | | | | |
|---|---|---|---|---|---|---|---|---|---|---|---|---|---|
| OM-RGC.v1.014810629 | GB_GCA_905612305.1 | 87.3 | Bacteroidota | Bacteroidia | Flavobacteriales | BACL11 | UWMA-0277 | UWMA-0277 sp905612305 | 0.519 | 0.016 | 0.518 | Cytoplasm | M3 |
| OM-RGC.v1.018010008 | GB_GCA_028825365.1 | 95.8 | Bacteroidota | Bacteroidia | Flavobacteriales | Flavobacteriaceae | SGZJ01 | SGZJ01 sp028825365 | 0.023 | 0.990 | 0.000 | Periplasm | M3 |
| OM-RGC.v1.018377826 | GB_GCA_002727775.1 | 96.2 | Pseudomonadota | Gammaproteobacteria | Pseudomonadales | Azotimanducaceae_A | UBA4421 | UBA4421 sp002727775 | 0.973 | 0.022 | 0.000 | Cytoplasm | M3 |
| OM-RGC.v1.018743943 | GB_GCA_030744915.1 | 100.0 | Pseudomonadota | Gammaproteobacteria | Pseudomonadales | Pseudohongiellaceae | UBA9145 | UBA9145 sp003451715 | 0.965 | 0.007 | 0.000 | Cytoplasm | M3 |
| gene_1016734\|GeneMark.hmm\|147_aa\|+\|3\|446_523805 | GB_GCA_002313925.1 | 96.6 | Gemmatimonadota | Gemmatimonadetes | Longimicrobiales | UBA6960 | UBA1142 | UBA1142 sp002313925 | 0.508 | 0.000 | 0.059 | Cytoplasm | M3_strict |
| gene_1026470\|GeneMark.hmm\|193_aa\|-\|1009\|1590_489237 | GB_GCA_949372635.1 | 95.3 | Pseudomonadota | Gammaproteobacteria | Pseudomonadales | Pseudohongiellaceae | UBA9145 | UBA9145 sp949372635 | 0.072 | 0.001 | 0.380 | Extracellular | M3_strict |
| gene_109989\|GeneMark.hmm\|245_aa\|-\|5643\|6380_55475 | RS_GCF_000355675.1 | 98.4 | Pseudomonadota | Gammaproteobacteria | Pseudomonadales | DSM-6294 | Thalassolituus | Thalassolituus oleivorans | 0.949 | 0.000 | 0.005 | Cytoplasm | M3_strict |
| gene_1423506\|GeneMark.hmm\|152_aa\|+\|1\|459_715061 | GB_GCA_002724925.1 | 73.2 | Pseudomonadota | Gammaproteobacteria | Pseudomonadales | DSM-6294 | Oleibacter | Oleibacter sp002724925 | 0.882 | 0.003 | 0.092 | Cytoplasm | M3_strict |
| gene_146818\|GeneMark.hmm\|179_aa\|-\|3\|539_75732 | GB_GCA_004213995.1 | 88.2 | Bacteroidota | Bacteroidia | Flavobacteriales | Flavobacteriaceae | SGZJ01 | SGZJ01 sp004213995 | 0.041 | 0.998 | 0.000 | Periplasm | M3_strict |
| gene_1482122\|GeneMark.hmm\|148_aa\|+\|1\|447_761018 | GB_GCA_020439995.1 | 63.3 | Verrucomicrobiota | Verrucomicrobiae | Verrucomicrobiales | DEV007 | JAGROY01 | JAGROY01 sp020439995 | 0.831 | 0.001 | 0.011 | Cytoplasm | M3_strict |
| gene_158516\|GeneMark.hmm\|188_aa\|-\|3\|566_78197 | RS_GCF_000355905.1 | 83.2 | Bacteroidota | Bacteroidia | Flavobacteriales | Flavobacteriaceae | Psychroflexus | Psychroflexus gondwanensis | 0.065 | 0.706 | 0.000 | Periplasm | M3_strict |
| gene_297484\|GeneMark.hmm\|211_aa\|-\|164\|799_149428 | RS_GCF_003052585.1 | 91.5 | Pseudomonadota | Gammaproteobacteria | Pseudomonadales | Pseudomonadaceae | Pseudomonas_E | Pseudomonas_E mangrovi | 0.482 | 0.013 | 0.493 | Extracellular | M3_strict |
| gene_365600\|GeneMark.hmm\|377_aa\|+\|3454\|4587_179795 | GB_GCA_000170815.1 | 61.6 | Bacteroidota | Bacteroidia | Flavobacteriales | Flavobacteriaceae | Patiriisocius | Patiriisocius sp000170815 | 0.004 | 0.845 | 0.155 | Periplasm | M3_strict |
| gene_616298\|GeneMark.hmm\|250_aa\|+\|2\|754_350161 | RS_GCF_003688405.1 | 78.4 | Bacteroidota | Bacteroidia | Flavobacteriales | Flavobacteriaceae | Ulvibacter | Ulvibacter antarcticus | 0.075 | 0.229 | 0.293 | Extracellular | M3_strict |
| gene_862723\|GeneMark.hmm\|209_aa\|-\|3\|629_431484 | GB_GCA_905181825.1 | 99.5 | Bacteroidota | Bacteroidia | Flavobacteriales | Flavobacteriaceae | Hel1-33-131 | Hel1-33-131 sp905181825 | 0.013 | 0.995 | 0.000 | Periplasm | M3_strict |
| gene_870527\|GeneMark.hmm\|138_aa\|+\|1\|417_432251 | RS_GCF_030070105.1 | 87.7 | Pseudomonadota | Gammaproteobacteria | Pseudomonadales | DSM-6294 | Bermanella | Bermanella sp030070105 | 0.039 | 0.973 | 0.000 | Periplasm | M3_strict |
| gene_9327\|GeneMark.hmm\|182_aa\|-\|97\|645_4668 | GB_GCA_002354735.1 | 84.1 | Pseudomonadota | Gammaproteobacteria | Pseudomonadales | Oleiphilaceae | UBA2679 | UBA2679 sp002354735 | 0.081 | 0.990 | 0.000 | Periplasm | M3_strict |
| gene_949725\|GeneMark.hmm\|102_aa\|-\|2\|307_471436 | GB_GCA_018675825.1 | 78.4 | Bacteroidota | Bacteroidia | Flavobacteriales | BACL11 | GCA-002715885 | GCA-002715885 sp018675825 | 0.010 | 1.000 | 0.000 | Periplasm | M3_strict |
| gene_956802\|GeneMark.hmm\|191_aa\|-\|847\|1422_473856 | GB_GCA_029938135.1 | 84.3 | Pseudomonadota | Gammaproteobacteria | Pseudomonadales | DSM-6294 | Bermanella | Bermanella sp029938135 | 0.104 | 0.385 | 0.168 | Periplasm | M3_strict |
| gene_98230\|GeneMark.hmm\|244_aa\|-\|36\|770_51834 | GB_GCA_002313925.1 | 94.3 | Gemmatimonadota | Gemmatimonadetes | Longimicrobiales | UBA6960 | UBA1142 | UBA1142 sp002313925 | 0.174 | 0.004 | 0.129 | Cytoplasm | M3_strict |

| | | | | | | | | | | | | | |
|---|---|---|---|---|---|---|---|---|---|---|---|---|---|
| gene_994125\|GeneMark.hmm\|188_aa\|-\|58\|624_509422 | GB_GCA_002731735.1 | 93.1 | Gemmatimonadota | Gemmatimonadetes | Longimicrobiales | UBA6960 | UBA822 | UBA822 sp002731735 | 0.928 | 0.000 | 0.001 | Cytoplasm | M3_strict |
| NODE_118344_length_541_cov_1.382716_1 | GB_GCA_002713705.1 | 92.7 | Bacteroidota | Bacteroidia | Flavobacteriales | Flavobacteriaceae | GCA-002733185 | GCA-002733185 sp002713705 | 0.473 | 0.075 | 0.080 | Cytoplasm | M3_strict |
| NODE_54999_length_820_cov_1.777778_1 | GB_GCA_938082955.1 | 88.2 | Pseudomonadota | Gammaproteobacteria | Pseudomonadales | Pseudohongiellaceae | UBA9145 | UBA9145 sp938082955 | 0.712 | 0.000 | 0.040 | Cytoplasm | M3_strict |
| OM-RGC.v1.003841053 | GB_GCA_016704485.1 | 54.2 | Bacteroidota | Bacteroidia | Flavobacteriales | PHOS-HE28 | PHOS-HE28 | PHOS-HE28 sp016704485 | 0.004 | 0.980 | 0.019 | Periplasm | M3_strict |
| OM-RGC.v1.009079167 | GB_GCA_002691285.1 | 76.8 | Marinisomatota | Marinisomatia | Marinisomatales | UBA1611 | GCA-002708715 | GCA-002708715 sp002691285 | 0.022 | 0.782 | 0.216 | Periplasm | M3_strict |
| OM-RGC.v1.010419055 | GB_GCA_902579065.1 | 77.1 | Bacteroidota | Bacteroidia | Flavobacteriales | BACL11 | GCA-002715885 | GCA-002715885 sp902579065 | 0.047 | 0.912 | 0.047 | Periplasm | M3_strict |
| OM-RGC.v1.011038036 | GB_GCA_002704685.1 | 60.8 | Marinisomatota | Marinisomatia | SCGC-AAA003-L08 | GCA-002707645 | GCA-002707645 | GCA-002707645 sp002704685 | 0.138 | 0.860 | 0.000 | Periplasm | M3_strict |
| OM-RGC.v1.011153177 | GB_GCA_002731735.1 | 94.7 | Gemmatimonadota | Gemmatimonadetes | Longimicrobiales | UBA6960 | UBA822 | UBA822 sp002731735 | 0.843 | 0.022 | 0.178 | Cytoplasm | M3_strict |
| OM-RGC.v1.011640759 | GB_GCA_002375495.1 | 98.7 | Bacteroidota | Bacteroidia | Flavobacteriales | Flavobacteriaceae | Marinirhabdus | Marinirhabdus sp002375495 | 0.870 | 0.003 | 0.010 | Cytoplasm | M3_strict |
| OM-RGC.v1.011859498 | GB_GCA_018675825.1 | 65.1 | Bacteroidota | Bacteroidia | Flavobacteriales | BACL11 | GCA-002715885 | GCA-002715885 sp018675825 | 0.958 | 0.011 | 0.000 | Cytoplasm | M3_strict |
| OM-RGC.v1.011985504 | GB_GCA_002171125.1 | 53.8 | Marinisomatota | Marinisomatia | SCGC-AAA003-L08 | TMED161 | TMED161 | TMED161 sp002171125 | 0.518 | 0.259 | 0.000 | Cytoplasm | M3_strict |
| OM-RGC.v1.012390758 | GB_GCA_018675825.1 | 65.2 | Bacteroidota | Bacteroidia | Flavobacteriales | BACL11 | GCA-002715885 | GCA-002715885 sp018675825 | 0.005 | 0.049 | 0.951 | Extracellular | M3_strict |
| OM-RGC.v1.012733741 | RS_GCF_004022485.1 | 51.6 | Bacteroidota | Bacteroidia | Flavobacteriales | Flavobacteriaceae | Aequorivita | Aequorivita sp004022485 | 0.048 | 0.946 | 0.051 | Periplasm | M3_strict |
| OM-RGC.v1.012885762 | GB_GCA_002728175.1 | 58.8 | Bacteroidota | Bacteroidia | Flavobacteriales | BACL11 | GCA-2697505 | GCA-2697505 sp002728175 | 0.946 | 0.000 | 0.002 | Cytoplasm | M3_strict |
| OM-RGC.v1.013906147 | GB_GCA_018674025.1 | 94.1 | Bacteroidota | Bacteroidia | Flavobacteriales | Flavobacteriaceae | SGZJ01 | SGZJ01 sp018674025 | 0.040 | 0.995 | 0.000 | Periplasm | M3_strict |
| OM-RGC.v1.014276511 | GB_GCA_002171125.1 | 64.1 | Marinisomatota | Marinisomatia | SCGC-AAA003-L08 | TMED161 | TMED161 | TMED161 sp002171125 | 0.236 | 0.171 | 0.342 | Extracellular | M3_strict |
| OM-RGC.v1.014915678 | GB_GCA_018675825.1 | 66.7 | Bacteroidota | Bacteroidia | Flavobacteriales | BACL11 | GCA-002715885 | GCA-002715885 sp018675825 | 0.744 | 0.003 | 0.151 | Cytoplasm | M3_strict |
| OM-RGC.v1.015400321 | GB_GCA_028122735.1 | 86.9 | Bacteroidota | Bacteroidia | Flavobacteriales | BACL11 | GCA-002715885 | GCA-002715885 sp028122735 | 0.966 | 0.000 | 0.006 | Cytoplasm | M3_strict |
| OM-RGC.v1.015644235 | GB_GCA_002691285.1 | 71.4 | Marinisomatota | Marinisomatia | Marinisomatales | UBA1611 | GCA-002708715 | GCA-002708715 sp002691285 | 0.021 | 0.579 | 0.365 | Periplasm | M3_strict |

| | | | | | | | | | | | | | |
|---|---|---|---|---|---|---|---|---|---|---|---|---|---|
| OM-RGC.v1.015683812 | GB_GCA_002718285.1 | 91.7 | Marinisomatota | Marinisomatia | Marinisomatales | UBA1611 | UBA2134 | UBA2134 sp002718285 | 0.327 | 0.210 | 0.313 | Cytoplasm | M3_strict |
| OM-RGC.v1.016600741 | GB_GCA_002724895.1 | 70.2 | Bacteroidota | Bacteroidia | Flavobacteriales | BACL11 | GCA-2693335 | GCA-2693335 sp002724895 | 0.177 | 0.093 | 0.253 | Extracellular | M3_strict |
| OM-RGC.v1.016858116 | GB_GCA_012960225.1 | 96.9 | Gemmatimonadota | Gemmatimonadetes | Longimicrobiales | UBA6960 | UBA1142 | UBA1142 sp012960225 | 0.961 | 0.000 | 0.001 | Cytoplasm | M3_strict |
| OM-RGC.v1.017132898 | GB_GCA_002691285.1 | 69.6 | Marinisomatota | Marinisomatia | Marinisomatales | UBA1611 | GCA-002708715 | GCA-002708715 sp002691285 | 0.020 | 0.576 | 0.398 | Periplasm | M3_strict |
| OM-RGC.v1.018982081 | GB_GCA_002691285.1 | 90.8 | Marinisomatota | Marinisomatia | Marinisomatales | UBA1611 | GCA-002708715 | GCA-002708715 sp002691285 | 0.013 | 0.690 | 0.257 | Periplasm | M3_strict |
| OM-RGC.v1.019511658 | GB_GCA_004214175.1 | 98.9 | Bacteroidota | Bacteroidia | Flavobacteriales | Flavobacteriaceae | GCA-002733185 | GCA-002733185 sp004214175 | 0.036 | 0.089 | 0.786 | Extracellular | M3_strict |
| OM-RGC.v1.020969699 | GB_GCA_002704685.1 | 83.2 | Marinisomatota | Marinisomatia | SCGC-AAA003-L08 | GCA-002707645 | GCA-002707645 | GCA-002707645 sp002704685 | 0.032 | 0.999 | 0.000 | Periplasm | M3_strict |
| OM-RGC.v1.021496243 | GB_GCA_002168105.2 | 91.2 | Planctomycetota | Phycisphaerae | Phycisphaerales | SM1A02 | UBA8653 | UBA8653 sp002168105 | 0.539 | 0.000 | 0.073 | Cytoplasm | M3_strict |
| OM-RGC.v1.021999261 | GB_GCA_002683195.1 | 98.8 | Planctomycetota | Phycisphaerae | Phycisphaerales | SM1A02 | GCA-002862325 | GCA-002862325 sp002683195 | 0.238 | 0.002 | 0.194 | Cytoplasm | M3_strict |
| OM-RGC.v1.024275817 | GB_GCA_030749035.1 | 98.0 | Myxococcota | XYA12-FULL-58-9 | XYA12-FULL-58-9 | JASMKV01 | JASMKV01 | JASMKV01 sp030749035 | 0.108 | 0.008 | 0.274 | Extracellular | M3_strict |
| gene_1117313\|GeneMark.hmm\|229_aa\|+\|3\|692_578386 | GB_GCA_938082955.1 | 98.7 | Pseudomonadota | Gammaproteobacteria | Pseudomonadales | Pseudohongiellaceae | UBA9145 | UBA9145 sp938082955 | 0.159 | 0.022 | 0.227 | Extracellular | M4 |
| gene_157716\|GeneMark.hmm\|297_aa\|-\|559\|1452_80508 | GB_GCA_002340765.1 | 98.6 | Bacteroidota | Bacteroidia | Flavobacteriales | Flavobacteriaceae | Polaribacter | Polaribacter sp002340765 | 0.004 | 0.384 | 0.615 | Extracellular | M4 |
| gene_19826\|GeneMark.hmm\|305_aa\|-\|12430\|13347_9719 | GB_GCA_002457215.1 | 98.4 | Pseudomonadota | Gammaproteobacteria | Pseudomonadales | Pseudohongiellaceae | UBA9145 | UBA9145 sp002457215 | 0.006 | 0.839 | 0.161 | Periplasm | M4 |
| gene_215830\|GeneMark.hmm\|311_aa\|-\|52\|987_110996 | GB_GCA_943789485.1 | 82.3 | Pseudomonadota | Gammaproteobacteria | Pseudomonadales | Pseudohongiellaceae | UBA9145 | UBA9145 sp943789485 | 0.007 | 0.087 | 0.913 | Extracellular | M4 |
| gene_244426\|GeneMark.hmm\|306_aa\|-\|3016\|3936_122766 | GB_GCA_002694855.1 | 100.0 | Pseudomonadota | Gammaproteobacteria | Pseudomonadales | Pseudohongiellaceae | UBA9145 | UBA9145 sp002694855 | 0.005 | 0.631 | 0.369 | Periplasm | M4 |
| gene_29487\|GeneMark.hmm\|307_aa\|+\|55080\|56003_14877 | GB_GCA_012960305.1 | 99.7 | Pseudomonadota | Gammaproteobacteria | Pseudomonadales | Pseudohongiellaceae | UBA9145 | UBA9145 sp012960305 | 0.007 | 0.544 | 0.456 | Periplasm | M4 |
| gene_411737\|GeneMark.hmm\|370_aa\|+\|35500\|36612_203524 | GB_GCA_003537695.1 | 99.5 | Bacteroidota | Bacteroidia | Flavobacteriales | Flavobacteriaceae | Polaribacter | Polaribacter sp003537695 | 0.004 | 0.993 | 0.007 | Periplasm | M4 |
| gene_4335\|GeneMark.hmm\|375_aa\|+\|282\|1409_2048 | GB_GCA_002384205.1 | 93.5 | Bacteroidota | Bacteroidia | NS11-12g | UBA9320 | UBA9320 | UBA9320 sp002384205 | 0.004 | 0.984 | 0.015 | Periplasm | M4 |
| gene_44648\|GeneMark.hmm\|368_aa\|-\|1852\|2958_23138 | GB_GCA_905182495.1 | 100.0 | Bacteroidota | Bacteroidia | Flavobacteriales | Crocinitomicaceae | UBA952 | UBA952 sp905182495 | 0.004 | 0.992 | 0.008 | Periplasm | M4 |
| gene_47475\|GeneMark.hmm\|370_aa\|+\|3149\|4261_24276 | GB_GCA_003537695.1 | 60.5 | Bacteroidota | Bacteroidia | Flavobacteriales | Flavobacteriaceae | Polaribacter | Polaribacter sp003537695 | 0.004 | 0.985 | 0.015 | Periplasm | M4 |

| | | | | | | | | | | | | | |
|---|---|---|---|---|---|---|---|---|---|---|---|---|---|
| gene_485677\|GeneMark.hmm\|289_aa\|+\|522\|1391_221903 | GB_GCA_913048385.1 | 88.9 | Pseudomonadota | Gammaproteobacteria | Pseudomonadales | Pseudohongiellaceae | UBA9145 | UBA9145 sp913048385 | 0.028 | 0.000 | 0.845 | Extracellular | M4 |
| gene_567988\|GeneMark.hmm\|394_aa\|+\|1638\|2822_278936 | GB_GCA_028825365.1 | 83.4 | Bacteroidota | Bacteroidia | Flavobacteriales | Flavobacteriaceae | SGZJ01 | SGZJ01 sp028825365 | 0.005 | 0.999 | 0.000 | Periplasm | M4 |
| gene_662964\|GeneMark.hmm\|368_aa\|-\|926\|2032_319858 | GB_GCA_905181635.1 | 96.2 | Bacteroidota | Bacteroidia | Flavobacteriales | BACL11 | UWMA-0277 | UWMA-0277 sp905181635 | 0.004 | 1.000 | 0.000 | Periplasm | M4 |
| gene_737601\|GeneMark.hmm\|318_aa\|-\|53\|1009_383613 | GB_GCA_913058275.1 | 93.2 | Pseudomonadota | Gammaproteobacteria | Pseudomonadales | Pseudohongiellaceae | UBA9145 | UBA9145 sp913058275 | 0.007 | 0.140 | 0.859 | Extracellular | M4 |
| gene_756391\|GeneMark.hmm\|382_aa\|-\|326\|1474_391550 | GB_GCA_016719225.1 | 55.8 | Bacteroidota | Bacteroidia | Flavobacteriales | PHOS-HE28 | PHOS-HE28 | PHOS-HE28 sp016719225 | 0.004 | 0.999 | 0.000 | Periplasm | M4 |
| gene_757048\|GeneMark.hmm\|377_aa\|-\|10004\|11137_370028 | GB_GCA_905182175.1 | 94.4 | Bacteroidota | Bacteroidia | Flavobacteriales | Vicingaceae | NORP294 | NORP294 sp905182175 | 0.004 | 0.999 | 0.001 | Periplasm | M4 |
| gene_841751\|GeneMark.hmm\|216_aa\|-\|2\|649_409020 | GB_GCA_949511925.1 | 91.6 | Bacteroidota | Bacteroidia | Flavobacteriales | BACL11 | UWMA-0277 | UWMA-0277 sp949511925 | 0.115 | 0.001 | 0.344 | Extracellular | M4 |
| gene_88267\|GeneMark.hmm\|304_aa\|-\|2971\|3885_46951 | GB_GCA_028823435.1 | 75.8 | Bacteroidota | Bacteroidia | Flavobacteriales | UA16 | UBA974 | UBA974 sp028823435 | 0.004 | 0.708 | 0.292 | Periplasm | M4 |
| gene_960196\|GeneMark.hmm\|303_aa\|-\|1023\|1934_528068 | GB_GCA_013204155.1 | 64.3 | Pseudomonadota | Gammaproteobacteria | Pseudomonadales | Pseudohongiellaceae | UBA9145 | UBA9145 sp013204155 | 0.007 | 0.006 | 0.992 | Extracellular | M4 |
| gene_98393\|GeneMark.hmm\|259_aa\|-\|3\|779_52196 | GB_GCA_004213995.1 | 86.9 | Bacteroidota | Bacteroidia | Flavobacteriales | Flavobacteriaceae | SGZJ01 | SGZJ01 sp004213995 | 0.076 | 0.972 | 0.000 | Periplasm | M4 |
| k141_1194293_39 | GB_GCA_905182495.1 | 97.3 | Bacteroidota | Bacteroidia | Flavobacteriales | Crocinitomicaceae | UBA952 | UBA952 sp905182495 | 0.004 | 0.986 | 0.014 | Periplasm | M4 |
| k141_1439800_50 | GB_GCA_002453265.1 | 100.0 | Bacteroidota | Bacteroidia | Flavobacteriales | UBA7430 | UBA6772 | UBA6772 sp002453265 | 0.004 | 1.000 | 0.000 | Periplasm | M4 |
| k141_2035086_31 | GB_GCA_949511925.1 | 97.9 | Bacteroidota | Bacteroidia | Flavobacteriales | BACL11 | UWMA-0277 | UWMA-0277 sp949511925 | 0.004 | 0.992 | 0.007 | Periplasm | M4 |
| k141_611722_3 | GB_GCA_028825365.1 | 99.4 | Bacteroidota | Bacteroidia | Flavobacteriales | Flavobacteriaceae | SGZJ01 | SGZJ01 sp028825365 | 0.004 | 0.998 | 0.002 | Periplasm | M4 |
| k141_628896_7 | GB_GCA_003537695.1 | 99.2 | Bacteroidota | Bacteroidia | Flavobacteriales | Flavobacteriaceae | Polaribacter | Polaribacter sp003537695 | 0.004 | 0.990 | 0.010 | Periplasm | M4 |
| NODE_146_length_44268_cov_7.716984_3 | GB_GCA_905182495.1 | 98.1 | Bacteroidota | Bacteroidia | Flavobacteriales | Crocinitomicaceae | UBA952 | UBA952 sp905182495 | 0.016 | 0.998 | 0.000 | Periplasm | M4 |
| NODE_1847_length_14967_cov_19.3284_5 | GB_GCA_003537695.1 | 99.6 | Bacteroidota | Bacteroidia | Flavobacteriales | Flavobacteriaceae | Polaribacter | Polaribacter sp003537695 | 0.039 | 0.946 | 0.000 | Periplasm | M4 |
| NODE_19_length_93834_cov_8.209972_2 | GB_GCA_949511925.1 | 97.3 | Bacteroidota | Bacteroidia | Flavobacteriales | BACL11 | UWMA-0277 | UWMA-0277 sp949511925 | 0.029 | 0.999 | 0.000 | Periplasm | M4 |
| NODE_39230_length_1107_cov_2.134981_1 | GB_GCA_943789485.1 | 92.2 | Pseudomonadota | Gammaproteobacteria | Pseudomonadales | Pseudohongiellaceae | UBA9145 | UBA9145 sp943789485 | 0.782 | 0.011 | 0.289 | Cytoplasm | M4 |
| NODE_8608_length_2183_cov_14.242_1 | GB_GCA_028825365.1 | 99.2 | Bacteroidota | Bacteroidia | Flavobacteriales | Flavobacteriaceae | SGZJ01 | SGZJ01 sp028825365 | 0.017 | 0.991 | 0.003 | Periplasm | M4 |

| | | | | | | | | | | | | | |
|---|---|---|---|---|---|---|---|---|---|---|---|---|---|
| OM-RGC.v1.002144010 | GB_GCA_002453265.1 | 99.8 | Bacteroidota | Bacteroidia | Flavobacteriales | UBA7430 | UBA6772 | UBA6772 sp002453265 | 0.004 | 1.000 | 0.000 | Periplasm | M4 |
| OM-RGC.v1.003293518 | GB_GCA_905478185.1 | 68.4 | Bacteroidota | Bacteroidia | Flavobacteriales | BACL11 | UWMA-0277 | UWMA-0277 sp905478185 | 0.004 | 1.000 | 0.000 | Periplasm | M4 |
| OM-RGC.v1.004123714 | GB_GCA_003537695.1 | 99.5 | Bacteroidota | Bacteroidia | Flavobacteriales | Flavobacteriaceae | Polaribacter | Polaribacter sp003537695 | 0.004 | 0.991 | 0.008 | Periplasm | M4 |
| OM-RGC.v1.004126835 | GB_GCA_913046365.1 | 83.9 | Bacteroidota | Bacteroidia | Flavobacteriales | Crocinitomicaceae | UBA952 | UBA952 sp002480215 | 0.004 | 0.994 | 0.006 | Periplasm | M4 |
| OM-RGC.v1.004206562 | GB_GCA_913051935.1 | 99.2 | Bacteroidota | Bacteroidia | Flavobacteriales | Crocinitomicaceae | UBA952 | UBA952 sp913051935 | 0.004 | 0.991 | 0.009 | Periplasm | M4 |
| OM-RGC.v1.004264467 | GB_GCA_913031405.1 | 99.2 | Bacteroidota | Bacteroidia | Flavobacteriales | Crocinitomicaceae | UBA952 | UBA952 sp913031405 | 0.004 | 0.985 | 0.015 | Periplasm | M4 |
| OM-RGC.v1.004351701 | GB_GCA_905181635.1 | 61.0 | Bacteroidota | Bacteroidia | Flavobacteriales | BACL11 | UWMA-0277 | UWMA-0277 sp905181635 | 0.004 | 1.000 | 0.000 | Periplasm | M4 |
| OM-RGC.v1.004831491 | GB_GCA_002714195.1 | 100.0 | Pseudomonadota | Gammaproteobacteria | Pseudomonadales | Pseudohongiellaceae | UBA9145 | UBA9145 sp002714195 | 0.020 | 0.691 | 0.283 | Periplasm | M4 |
| OM-RGC.v1.005368315 | GB_GCA_002167775.1 | 79.0 | Bacteroidota | Bacteroidia | Flavobacteriales | Crocinitomicaceae | UBA952 | UBA952 sp002167775 | 0.009 | 0.987 | 0.011 | Periplasm | M4 |
| OM-RGC.v1.005498887 | GB_GCA_949511925.1 | 71.3 | Bacteroidota | Bacteroidia | Flavobacteriales | BACL11 | UWMA-0277 | UWMA-0277 sp949511925 | 0.009 | 0.871 | 0.111 | Periplasm | M4 |
| OM-RGC.v1.005627526 | GB_GCA_905478185.1 | 89.7 | Bacteroidota | Bacteroidia | Flavobacteriales | BACL11 | UWMA-0277 | UWMA-0277 sp905478185 | 0.004 | 1.000 | 0.000 | Periplasm | M4 |
| OM-RGC.v1.005689078 | GB_GCA_905479165.1 | 67.5 | Bacteroidota | Bacteroidia | Flavobacteriales | BACL11 | UWMA-0277 | UWMA-0277 sp905479165 | 0.004 | 0.996 | 0.004 | Periplasm | M4 |
| OM-RGC.v1.006352937 | GB_GCA_028818225.1 | 88.9 | Pseudomonadota | Gammaproteobacteria | Pseudomonadales | Pseudohongiellaceae | UBA9145 | UBA9145 sp028818225 | 0.010 | 0.153 | 0.846 | Extracellular | M4 |
| OM-RGC.v1.006563570 | GB_GCA_949511925.1 | 70.5 | Bacteroidota | Bacteroidia | Flavobacteriales | BACL11 | UWMA-0277 | UWMA-0277 sp949511925 | 0.005 | 1.000 | 0.000 | Periplasm | M4 |
| OM-RGC.v1.006696243 | GB_GCA_002694855.1 | 100.0 | Pseudomonadota | Gammaproteobacteria | Pseudomonadales | Pseudohongiellaceae | UBA9145 | UBA9145 sp002694855 | 0.005 | 0.631 | 0.369 | Periplasm | M4 |
| OM-RGC.v1.006777490 | GB_GCA_002457215.1 | 99.7 | Pseudomonadota | Gammaproteobacteria | Pseudomonadales | Pseudohongiellaceae | UBA9145 | UBA9145 sp002457215 | 0.006 | 0.905 | 0.094 | Periplasm | M4 |
| OM-RGC.v1.006808752 | GB_GCA_028823435.1 | 75.8 | Bacteroidota | Bacteroidia | Flavobacteriales | UA16 | UBA974 | UBA974 sp028823435 | 0.004 | 0.911 | 0.089 | Periplasm | M4 |
| OM-RGC.v1.006891220 | GB_GCA_002700775.1 | 97.4 | Pseudomonadota | Gammaproteobacteria | Pseudomonadales | Pseudohongiellaceae | UBA9145 | UBA9145 sp002700775 | 0.006 | 0.006 | 0.994 | Extracellular | M4 |
| OM-RGC.v1.006939680 | GB_GCA_002477465.1 | 100.0 | Pseudomonadota | Gammaproteobacteria | Pseudomonadales | Pseudohongiellaceae | UBA9145 | UBA9145 sp002477465 | 0.007 | 0.030 | 0.969 | Extracellular | M4 |
| OM-RGC.v1.006974294 | GB_GCA_008081045.1 | 90.7 | Pseudomonadota | Gammaproteobacteria | Pseudomonadales | Pseudohongiellaceae | UBA9145 | UBA9145 sp008081045 | 0.010 | 0.001 | 0.998 | Extracellular | M4 |

| | | | | | | | | | | | | | |
|---|---|---|---|---|---|---|---|---|---|---|---|---|---|
| OM-RGC.v1.007141244 | RS_GCF_000153225.1 | 100.0 | Bacteroidota | Bacteroidia | Flavobacteriales | Flavobacteriaceae | Polaribacter | Polaribacter irgensii | 0.005 | 0.439 | 0.560 | Extracellular | M4 |
| OM-RGC.v1.007382880 | GB_GCA_913048385.1 | 100.0 | Pseudomonadota | Gammaproteobacteria | Pseudomonadales | Pseudohongiellaceae | UBA9145 | UBA9145 sp913048385 | 0.005 | 0.000 | 0.999 | Extracellular | M4 |
| OM-RGC.v1.007570743 | GB_GCA_905181635.1 | 87.6 | Bacteroidota | Bacteroidia | Flavobacteriales | BACL11 | UWMA-0277 | UWMA-0277 sp905181635 | 0.074 | 0.044 | 0.795 | Extracellular | M4 |
| OM-RGC.v1.008768647 | GB_GCA_913041775.1 | 100.0 | Bacteroidota | Bacteroidia | Flavobacteriales | UBA7430 | UBA6772 | UBA6772 sp913041775 | 0.038 | 0.938 | 0.014 | Periplasm | M4 |
| OM-RGC.v1.008889720 | GB_GCA_002726595.1 | 100.0 | Pseudomonadota | Gammaproteobacteria | Pseudomonadales | Pseudohongiellaceae | UBA9145 | UBA9145 sp002726595 | 0.433 | 0.006 | 0.076 | Cytoplasm | M4 |
| OM-RGC.v1.009318212 | GB_GCA_002471665.1 | 100.0 | Pseudomonadota | Gammaproteobacteria | Pseudomonadales | Ketobacteraceae | Ketobacter | Ketobacter sp002471665 | 0.015 | 0.785 | 0.215 | Periplasm | M4 |
| OM-RGC.v1.011593909 | GB_GCA_002714195.1 | 92.5 | Pseudomonadota | Gammaproteobacteria | Pseudomonadales | Pseudohongiellaceae | UBA9145 | UBA9145 sp002714195 | 0.123 | 0.955 | 0.000 | Periplasm | M4 |
| OM-RGC.v1.011914148 | GB_GCA_028825365.1 | 100.0 | Bacteroidota | Bacteroidia | Flavobacteriales | Flavobacteriaceae | SGZJ01 | SGZJ01 sp028825365 | 0.018 | 0.988 | 0.000 | Periplasm | M4 |
| OM-RGC.v1.012137754 | GB_GCA_011522905.1 | 97.4 | Bacteroidota | Bacteroidia | Flavobacteriales | Crocinitomicaceae | UBA952 | UBA952 sp011522905 | 0.755 | 0.632 | 0.000 | Cytoplasm | M4 |
| OM-RGC.v1.012406606 | GB_GCA_902597095.1 | 99.6 | Pseudomonadota | Gammaproteobacteria | Pseudomonadales | Pseudohongiellaceae | UBA9145 | UBA9145 sp902597095 | 0.246 | 0.001 | 0.046 | Cytoplasm | M4 |
| OM-RGC.v1.014562549 | GB_GCA_905181635.1 | 91.5 | Bacteroidota | Bacteroidia | Flavobacteriales | BACL11 | UWMA-0277 | UWMA-0277 sp905181635 | 0.173 | 0.000 | 0.219 | Extracellular | M4 |
| OM-RGC.v1.015463425 | GB_GCA_913061205.1 | 87.9 | Bacteroidota | Bacteroidia | Flavobacteriales | Flavobacteriaceae | GCA-002733185 | GCA-002733185 sp913061205 | 0.064 | 0.248 | 0.587 | Extracellular | M4 |
| gene_1116655\|GeneMark.hmm\|290_aa\|+\|2221\|3093_613124 | GB_GCA_015664885.1 | 99.7 | Gemmatimonadota | Gemmatimonadetes | Longimicrobiales | UBA6960 | UBA1138 | UBA1138 sp003447875 | 0.005 | 0.500 | 0.499 | Periplasm | M4_strict |
| gene_1148944\|GeneMark.hmm\|231_aa\|-\|3\|695_605446 | GB_GCA_018675825.1 | 95.2 | Bacteroidota | Bacteroidia | Flavobacteriales | BACL11 | GCA-002715885 | GCA-002715885 sp018675825 | 0.053 | 0.909 | 0.000 | Periplasm | M4_strict |
| gene_118676\|GeneMark.hmm\|365_aa\|+\|4995\|6092_59195 | GB_GCA_913047375.1 | 100.0 | Bacteroidota | Bacteroidia | Flavobacteriales | Flavobacteriaceae | Polaribacter | Polaribacter sp913047375 | 0.004 | 1.000 | 0.000 | Periplasm | M4_strict |
| gene_1248977\|GeneMark.hmm\|237_aa\|-\|2\|712_675408 | GB_GCA_027623555.1 | 69.5 | Pseudomonadota | Gammaproteobacteria | Pseudomonadales | Pseudohongiellaceae | UBA9145 | UBA9145 sp027623555 | 0.008 | 0.048 | 0.951 | Extracellular | M4_strict |
| gene_1266190\|GeneMark.hmm\|385_aa\|-\|11\|1168_610114 | GB_GCA_913020435.1 | 100.0 | Bacteroidota | Bacteroidia | Flavobacteriales | Salibacteraceae | UBA6770 | UBA6770 sp002453255 | 0.004 | 0.488 | 0.512 | Extracellular | M4_strict |
| gene_168645\|GeneMark.hmm\|293_aa\|+\|3616\|4497_88441 | GB_GCA_938082955.1 | 98.6 | Pseudomonadota | Gammaproteobacteria | Pseudomonadales | Pseudohongiellaceae | UBA9145 | UBA9145 sp938082955 | 0.010 | 0.114 | 0.885 | Extracellular | M4_strict |
| gene_187890\|GeneMark.hmm\|355_aa\|+\|10098\|11165_96614 | GB_GCA_018608625.1 | 98.9 | Bacteroidota | Bacteroidia | Flavobacteriales | Flavobacteriaceae | Hel1-33-131 | Hel1-33-131 sp018608625 | 0.005 | 0.996 | 0.000 | Periplasm | M4_strict |
| gene_250117\|GeneMark.hmm\|294_aa\|-\|951\|1835_138545 | GB_GCA_009692275.1 | 79.9 | Pseudomonadota | Gammaproteobacteria | Xanthomonadales | SZUA-36 | SHZL01 | SHZL01 sp009692275 | 0.012 | 0.000 | 0.998 | Extracellular | M4_strict |

| | | | | | | | | | | | | | |
|---|---|---|---|---|---|---|---|---|---|---|---|---|---|
| gene_25500\|GeneMark.hmm\|311_aa\|-\|1\|933_12901 | RS_GCF_003688405.1 | 84.1 | Bacteroidota | Bacteroidia | Flavobacteriales | Flavobacteriaceae | Ulvibacter | Ulvibacter antarcticus | 0.004 | 0.977 | 0.022 | Periplasm | M4_strict |
| gene_262650\|GeneMark.hmm\|283_aa\|-\|769\|1620_136792 | GB_GCA_002313925.1 | 97.5 | Gemmatimonadota | Gemmatimonadetes | Longimicrobiales | UBA6960 | UBA1142 | UBA1142 sp002313925 | 0.008 | 0.324 | 0.676 | Extracellular | M4_strict |
| gene_330287\|GeneMark.hmm\|320_aa\|+\|3\|965_160517 | GB_GCA_029212745.1 | 65.3 | Pseudomonadota | Gammaproteobacteria | Pseudomonadales | Alcanivoracaceae | JARGOL01 | JARGOL01 sp029212745 | 0.006 | 0.616 | 0.348 | Periplasm | M4_strict |
| gene_335482\|GeneMark.hmm\|292_aa\|+\|1349\|2227_187922 | RS_GCF_001444425.1 | 72.8 | Pseudomonadota | Gammaproteobacteria | Pseudomonadales | Pseudohongiellaceae | Pseudohongiella | Pseudohongiella spirulinae | 0.007 | 0.003 | 0.996 | Extracellular | M4_strict |
| gene_454339\|GeneMark.hmm\|272_aa\|-\|1\|816_227687 | RS_GCF_003052585.1 | 99.5 | Pseudomonadota | Gammaproteobacteria | Pseudomonadales | Pseudomonadaceae | Pseudomonas_E | Pseudomonas_E mangrovi | 0.005 | 0.949 | 0.051 | Periplasm | M4_strict |
| gene_584583\|GeneMark.hmm\|283_aa\|-\|21007\|21858_287309 | GB_GCA_002840055.1 | 99.3 | Pseudomonadota | Gammaproteobacteria | Pseudomonadales | Pseudomonadaceae | Halopseudomonas | Halopseudomonas sp002840055 | 0.086 | 0.658 | 0.294 | Periplasm | M4_strict |
| gene_636977\|GeneMark.hmm\|391_aa\|+\|2468\|3643_345038 | GB_GCA_913061205.1 | 95.7 | Bacteroidota | Bacteroidia | Flavobacteriales | Flavobacteriaceae | GCA-002733185 | GCA-002733185 sp913061205 | 0.005 | 0.960 | 0.039 | Periplasm | M4_strict |
| gene_745775\|GeneMark.hmm\|339_aa\|-\|796\|1815_389989 | GB_GCA_913048385.1 | 90.8 | Pseudomonadota | Gammaproteobacteria | Pseudomonadales | Pseudohongiellaceae | UBA9145 | UBA9145 sp913048385 | 0.053 | 0.002 | 0.867 | Extracellular | M4_strict |
| gene_780190\|GeneMark.hmm\|223_aa\|-\|2\|670_432297 | GB_GCA_029212745.1 | 75.3 | Pseudomonadota | Gammaproteobacteria | Pseudomonadales | Alcanivoracaceae | JARGOL01 | JARGOL01 sp029212745 | 0.902 | 0.002 | 0.004 | Cytoplasm | M4_strict |
| gene_814362\|GeneMark.hmm\|365_aa\|+\|1732\|2829_404646 | GB_GCA_905182835.1 | 100.0 | Bacteroidota | Bacteroidia | Flavobacteriales | Flavobacteriaceae | SGZJ01 | SGZJ01 sp905182835 | 0.004 | 1.000 | 0.000 | Periplasm | M4_strict |
| gene_833794\|GeneMark.hmm\|270_aa\|+\|2982\|3794_414625 | RS_GCF_950066645.1 | 63.9 | Pseudomonadota | Gammaproteobacteria | Pseudomonadales | DSM-6294 | Oleibacter | Oleibacter sp950066645 | 0.007 | 0.001 | 0.996 | Extracellular | M4_strict |
| gene_83814\|GeneMark.hmm\|261_aa\|-\|2\|784_43566 | GB_GCA_913060785.1 | 97.7 | Bacteroidota | Bacteroidia | Flavobacteriales | Flavobacteriaceae | Patiriisocius | Patiriisocius sp913060785 | 0.005 | 0.861 | 0.139 | Periplasm | M4_strict |
| gene_846731\|GeneMark.hmm\|380_aa\|+\|10057\|11199_434434 | GB_GCA_913045325.1 | 99.7 | Bacteroidota | Bacteroidia | Flavobacteriales | Flavobacteriaceae | Patiriisocius | Patiriisocius sp003450155 | 0.004 | 0.975 | 0.025 | Periplasm | M4_strict |
| gene_849006\|GeneMark.hmm\|214_aa\|-\|1649\|2293_450138 | GB_GCA_029212745.1 | 76.6 | Pseudomonadota | Gammaproteobacteria | Pseudomonadales | Alcanivoracaceae | JARGOL01 | JARGOL01 sp029212745 | 0.675 | 0.000 | 0.035 | Cytoplasm | M4_strict |
| gene_872362\|GeneMark.hmm\|375_aa\|-\|1113\|2240_436058 | GB_GCA_949519155.1 | 86.4 | Bacteroidota | Bacteroidia | Flavobacteriales | Flavobacteriaceae | Polaribacter | Polaribacter sp949519155 | 0.004 | 0.991 | 0.009 | Periplasm | M4_strict |
| gene_88095\|GeneMark.hmm\|313_aa\|-\|1574\|2515_46287 | RS_GCF_030070105.1 | 93.3 | Pseudomonadota | Gammaproteobacteria | Pseudomonadales | DSM-6294 | Bermanella | Bermanella sp030070105 | 0.006 | 0.999 | 0.000 | Periplasm | M4_strict |
| gene_918586\|GeneMark.hmm\|365_aa\|+\|376\|1473_458168 | GB_GCA_913061165.1 | 92.3 | Bacteroidota | Bacteroidia | Flavobacteriales | Flavobacteriaceae | Polaribacter | Polaribacter sp913061165 | 0.004 | 0.999 | 0.001 | Periplasm | M4_strict |
| gene_921526\|GeneMark.hmm\|320_aa\|+\|76476\|77438_447987 | GB_GCA_002840055.1 | 100.0 | Pseudomonadota | Gammaproteobacteria | Pseudomonadales | Pseudomonadaceae | Halopseudomonas | Halopseudomonas sp002840055 | 0.005 | 0.944 | 0.056 | Periplasm | M4_strict |
| gene_95915\|GeneMark.hmm\|364_aa\|-\|2550\|3644_47738 | GB_GCA_004213995.1 | 79.9 | Bacteroidota | Bacteroidia | Flavobacteriales | Flavobacteriaceae | SGZJ01 | SGZJ01 sp004213995 | 0.004 | 1.000 | 0.000 | Periplasm | M4_strict |
| k141_1064804_7 | GB_GCA_913048215.1 | 77.0 | Bacteroidota | Bacteroidia | Flavobacteriales | Flavobacteriaceae | Patiriisocius | Patiriisocius sp913048215 | 0.004 | 0.976 | 0.024 | Periplasm | M4_strict |

| | | | | | | | | | | | | | |
|---|---|---|---|---|---|---|---|---|---|---|---|---|---|
| k141_1133623_3 | GB_GCA_938082955.1 | 92.7 | Pseudomonadota | Gammaproteobacteria | Pseudomonadales | Pseudohongiellaceae | UBA9145 | UBA9145 sp938082955 | 0.010 | 0.002 | 0.996 | Extracellular | M4_strict |
| k141_1222342_44 | RS_GCF_000355905.1 | 77.0 | Bacteroidota | Bacteroidia | Flavobacteriales | Flavobacteriaceae | Psychroflexus | Psychroflexus gondwanensis | 0.004 | 0.983 | 0.016 | Periplasm | M4_strict |
| k141_1306186_31 | GB_GCA_913061205.1 | 93.0 | Bacteroidota | Bacteroidia | Flavobacteriales | Flavobacteriaceae | GCA-002733185 | GCA-002733185 sp913061205 | 0.005 | 0.975 | 0.025 | Periplasm | M4_strict |
| k141_2034477_44 | GB_GCA_913047375.1 | 100.0 | Bacteroidota | Bacteroidia | Flavobacteriales | Flavobacteriaceae | Polaribacter | Polaribacter sp913047375 | 0.004 | 1.000 | 0.000 | Periplasm | M4_strict |
| k141_225199_2 | GB_GCA_949519155.1 | 100.0 | Bacteroidota | Bacteroidia | Flavobacteriales | Flavobacteriaceae | Polaribacter | Polaribacter sp949519155 | 0.004 | 0.863 | 0.137 | Periplasm | M4_strict |
| k141_247466_2 | GB_GCA_913045325.1 | 99.5 | Bacteroidota | Bacteroidia | Flavobacteriales | Flavobacteriaceae | Patiriisocius | Patiriisocius sp003450155 | 0.004 | 0.995 | 0.005 | Periplasm | M4_strict |
| k141_663913_1 | GB_GCA_002336245.1 | 91.2 | Bacteroidota | Bacteroidia | Flavobacteriales | Salibacteraceae | UBA4419 | UBA4419 sp002336245 | 0.005 | 0.748 | 0.252 | Periplasm | M4_strict |
| k141_758567_6 | GB_GCA_937771065.1 | 88.7 | Bacteroidota | Bacteroidia | Flavobacteriales | Flavobacteriaceae | Polaribacter | Polaribacter sp937771065 | 0.004 | 0.999 | 0.000 | Periplasm | M4_strict |
| NODE_1272_length_18520_cov_12.9147_8 | GB_GCA_949519155.1 | 100.0 | Bacteroidota | Bacteroidia | Flavobacteriales | Flavobacteriaceae | Polaribacter | Polaribacter sp949519155 | 0.037 | 0.809 | 0.183 | Periplasm | M4_strict |
| NODE_13170_length_4426_cov_7.95516_1 | GB_GCA_913048115.1 | 82.2 | Bacteroidota | Bacteroidia | Flavobacteriales | Flavobacteriaceae | Patiriisocius | Patiriisocius sp913048115 | 0.049 | 0.988 | 0.000 | Periplasm | M4_strict |
| NODE_20603_length_3292_cov_18.7328_3 | GB_GCA_913061165.1 | 91.1 | Bacteroidota | Bacteroidia | Flavobacteriales | Flavobacteriaceae | Polaribacter | Polaribacter sp913061165 | 0.079 | 0.895 | 0.000 | Periplasm | M4_strict |
| NODE_2214_length_6448_cov_5.131550_2 | GB_GCA_913047375.1 | 100.0 | Bacteroidota | Bacteroidia | Flavobacteriales | Flavobacteriaceae | Polaribacter | Polaribacter sp913047375 | 0.054 | 0.992 | 0.000 | Periplasm | M4_strict |
| NODE_275041_length_587_cov_4.07331_1 | GB_GCA_018608625.1 | 95.0 | Bacteroidota | Bacteroidia | Flavobacteriales | Flavobacteriaceae | Hel1-33-131 | Hel1-33-131 sp018608625 | 0.041 | 0.959 | 0.000 | Periplasm | M4_strict |
| NODE_32616_length_1237_cov_3.987310_1 | GB_GCA_002713705.1 | 95.0 | Bacteroidota | Bacteroidia | Flavobacteriales | Flavobacteriaceae | GCA-002733185 | GCA-002733185 sp002713705 | 0.102 | 0.834 | 0.093 | Periplasm | M4_strict |
| NODE_43048_length_1030_cov_1.475897_1 | GB_GCA_938082955.1 | 91.7 | Pseudomonadota | Gammaproteobacteria | Pseudomonadales | Pseudohongiellaceae | UBA9145 | UBA9145 sp938082955 | 0.931 | 0.000 | 0.007 | Cytoplasm | M4_strict |
| NODE_7872_length_3614_cov_2.993538_2 | GB_GCA_018067045.1 | 95.2 | Pseudomonadota | Gammaproteobacteria | Pseudomonadales | Alcanivoracaceae | JARGPN01 | JARGPN01 sp018067045 | 0.092 | 0.515 | 0.127 | Periplasm | M4_strict |
| NODE_886_length_7822_cov_18.7847_8 | GB_GCA_913045325.1 | 99.6 | Bacteroidota | Bacteroidia | Flavobacteriales | Flavobacteriaceae | Patiriisocius | Patiriisocius sp003450155 | 0.043 | 0.999 | 0.000 | Periplasm | M4_strict |
| NODE_94302_length_1181_cov_3.31616_1 | RS_GCF_008641105.1 | 93.1 | Pseudomonadota | Gammaproteobacteria | Pseudomonadales | Pseudomonadaceae | Halopseudomonas | Halopseudomonas salina | 0.705 | 0.002 | 0.055 | Cytoplasm | M4_strict |
| OM-RGC.v1.001682938 | GB_GCA_002339135.1 | 64.3 | Bacteroidota | Bacteroidia | Flavobacteriales | UA16 | UBA974 | UBA974 sp002339135 | 0.007 | 1.000 | 0.000 | Periplasm | M4_strict |
| OM-RGC.v1.003540295 | GB_GCA_002707145.1 | 78.3 | Bacteroidota | Bacteroidia | Flavobacteriales | UBA7430 | GCA-2707145 | GCA-2707145 sp002707145 | 0.004 | 1.000 | 0.000 | Periplasm | M4_strict |

| | | | | | | | | | | | | | |
|---|---|---|---|---|---|---|---|---|---|---|---|---|---|
| OM-RGC.v1.003709582 | GB_GCA_913020435.1 | 100.0 | Bacteroidota | Bacteroidia | Flavobacteriales | Salibacteraceae | UBA6770 | UBA6770 sp002453255 | 0.004 | 0.488 | 0.512 | Extracellular | M4_strict |
| OM-RGC.v1.003793749 | GB_GCA_002707145.1 | 99.7 | Bacteroidota | Bacteroidia | Flavobacteriales | UBA7430 | GCA-2707145 | GCA-2707145 sp002707145 | 0.048 | 0.992 | 0.000 | Periplasm | M4_strict |
| OM-RGC.v1.004036536 | GB_GCA_913061205.1 | 81.2 | Bacteroidota | Bacteroidia | Flavobacteriales | Flavobacteriaceae | GCA-002733185 | GCA-002733185 sp913061205 | 0.004 | 0.996 | 0.004 | Periplasm | M4_strict |
| OM-RGC.v1.004037402 | GB_GCA_913061205.1 | 93.0 | Bacteroidota | Bacteroidia | Flavobacteriales | Flavobacteriaceae | GCA-002733185 | GCA-002733185 sp913061205 | 0.005 | 0.975 | 0.025 | Periplasm | M4_strict |
| OM-RGC.v1.004094379 | GB_GCA_002339135.1 | 99.7 | Bacteroidota | Bacteroidia | Flavobacteriales | UA16 | UBA974 | UBA974 sp002339135 | 0.006 | 0.933 | 0.066 | Periplasm | M4_strict |
| OM-RGC.v1.004249645 | GB_GCA_002707745.1 | 100.0 | Bacteroidota | Bacteroidia | Flavobacteriales | Flavobacteriaceae | Marinirhabdus | Marinirhabdus sp002707745 | 0.004 | 0.869 | 0.131 | Periplasm | M4_strict |
| OM-RGC.v1.004257814 | GB_GCA_002337605.1 | 100.0 | Bacteroidota | Bacteroidia | Flavobacteriales | Flavobacteriaceae | Marinirhabdus | Marinirhabdus sp002337605 | 0.005 | 0.995 | 0.005 | Periplasm | M4_strict |
| OM-RGC.v1.004276195 | GB_GCA_000981625.1 | 100.0 | Bacteroidota | Bacteroidia | Flavobacteriales | Flavobacteriaceae | Polaribacter | Polaribacter sp000981625 | 0.004 | 1.000 | 0.000 | Periplasm | M4_strict |
| OM-RGC.v1.004276271 | GB_GCA_913047375.1 | 100.0 | Bacteroidota | Bacteroidia | Flavobacteriales | Flavobacteriaceae | Polaribacter | Polaribacter sp913047375 | 0.004 | 1.000 | 0.000 | Periplasm | M4_strict |
| OM-RGC.v1.004277188 | RS_GCF_003335675.1 | 84.7 | Bacteroidota | Bacteroidia | Flavobacteriales | Flavobacteriaceae | Winogradskyella | Winogradskyella sp003335675 | 0.004 | 0.998 | 0.002 | Periplasm | M4_strict |
| OM-RGC.v1.004281317 | GB_GCA_943842625.1 | 92.3 | Bacteroidota | Bacteroidia | Flavobacteriales | Flavobacteriaceae | GCA-002733185 | GCA-002733185 sp943842625 | 0.004 | 0.989 | 0.011 | Periplasm | M4_strict |
| OM-RGC.v1.004281770 | RS_GCF_000952855.1 | 96.2 | Bacteroidota | Bacteroidia | Flavobacteriales | Flavobacteriaceae | Aequorivita | Aequorivita vladivostokensis | 0.004 | 0.970 | 0.030 | Periplasm | M4_strict |
| OM-RGC.v1.004300731 | GB_GCA_018656255.1 | 97.8 | Bacteroidota | Bacteroidia | Flavobacteriales | Flavobacteriaceae | Patiriisocius | Patiriisocius sp018656255 | 0.004 | 0.996 | 0.004 | Periplasm | M4_strict |
| OM-RGC.v1.004303738 | RS_GCF_000153485.2 | 100.0 | Bacteroidota | Bacteroidia | Flavobacteriales | Flavobacteriaceae | Psychroflexus | Psychroflexus torquis | 0.006 | 0.998 | 0.002 | Periplasm | M4_strict |
| OM-RGC.v1.004307492 | GB_GCA_913048115.1 | 100.0 | Bacteroidota | Bacteroidia | Flavobacteriales | Flavobacteriaceae | Patiriisocius | Patiriisocius sp913048115 | 0.004 | 0.998 | 0.002 | Periplasm | M4_strict |
| OM-RGC.v1.004333666 | RS_GCF_000265385.1 | 100.0 | Bacteroidota | Bacteroidia | Flavobacteriales | Flavobacteriaceae | Aequorivita | Aequorivita sublithincola | 0.004 | 0.917 | 0.083 | Periplasm | M4_strict |
| OM-RGC.v1.004335166 | GB_GCA_000170815.1 | 100.0 | Bacteroidota | Bacteroidia | Flavobacteriales | Flavobacteriaceae | Patiriisocius | Patiriisocius sp000170815 | 0.004 | 0.978 | 0.022 | Periplasm | M4_strict |
| OM-RGC.v1.005177300 | GB_GCA_002704685.1 | 93.7 | Marinisomatota | Marinisomatia | SCGC-AAA003-L08 | GCA-002707645 | GCA-002707645 | GCA-002707645 sp002704685 | 0.006 | 0.982 | 0.015 | Periplasm | M4_strict |
| OM-RGC.v1.005241351 | GB_GCA_002691285.1 | 85.6 | Marinisomatota | Marinisomatia | Marinisomatales | UBA1611 | GCA-002708715 | GCA-002708715 sp002691285 | 0.006 | 0.933 | 0.066 | Periplasm | M4_strict |
| OM-RGC.v1.005266853 | GB_GCA_002718285.1 | 100.0 | Marinisomatota | Marinisomatia | Marinisomatales | UBA1611 | UBA2134 | UBA2134 sp002718285 | 0.013 | 0.990 | 0.000 | Periplasm | M4_strict |

| | | | | | | | | | | | | | |
|---|---|---|---|---|---|---|---|---|---|---|---|---|---|
| OM-RGC.v1.005349672 | GB_GCA_002705205.1 | 59.8 | Bacteroidota | Bacteroidia | Flavobacteriales | BACL11 | GCA-2705205 | GCA-2705205 sp002705205 | 0.004 | 0.991 | 0.008 | Periplasm | M4_strict |
| OM-RGC.v1.005477654 | GB_GCA_002728175.1 | 82.5 | Bacteroidota | Bacteroidia | Flavobacteriales | BACL11 | GCA-2697505 | GCA-2697505 sp002728175 | 0.004 | 0.949 | 0.051 | Periplasm | M4_strict |
| OM-RGC.v1.005560072 | GB_GCA_002705205.1 | 100.0 | Bacteroidota | Bacteroidia | Flavobacteriales | BACL11 | GCA-2705205 | GCA-2705205 sp002705205 | 0.006 | 0.982 | 0.018 | Periplasm | M4_strict |
| OM-RGC.v1.005573091 | GB_GCA_002728175.1 | 65.4 | Bacteroidota | Bacteroidia | Flavobacteriales | BACL11 | GCA-2697505 | GCA-2697505 sp002728175 | 0.005 | 0.994 | 0.006 | Periplasm | M4_strict |
| OM-RGC.v1.005582688 | GB_GCA_902579065.1 | 71.1 | Bacteroidota | Bacteroidia | Flavobacteriales | BACL11 | GCA-002715885 | GCA-002715885 sp902579065 | 0.005 | 0.958 | 0.041 | Periplasm | M4_strict |
| OM-RGC.v1.005604552 | GB_GCA_002724895.1 | 97.6 | Bacteroidota | Bacteroidia | Flavobacteriales | BACL11 | GCA-2693335 | GCA-2693335 sp002724895 | 0.005 | 0.162 | 0.836 | Extracellular | M4_strict |
| OM-RGC.v1.005625285 | GB_GCA_902579065.1 | 72.6 | Bacteroidota | Bacteroidia | Flavobacteriales | BACL11 | GCA-002715885 | GCA-002715885 sp902579065 | 0.004 | 0.996 | 0.003 | Periplasm | M4_strict |
| OM-RGC.v1.005734613 | GB_GCA_002171125.1 | 69.3 | Marinisomatota | Marinisomatia | SCGC-AAA003-L08 | TMED161 | TMED161 | TMED161 sp002171125 | 0.004 | 0.998 | 0.002 | Periplasm | M4_strict |
| OM-RGC.v1.005791302 | GB_GCA_002724895.1 | 63.0 | Bacteroidota | Bacteroidia | Flavobacteriales | BACL11 | GCA-2693335 | GCA-2693335 sp002724895 | 0.012 | 0.791 | 0.206 | Periplasm | M4_strict |
| OM-RGC.v1.005869309 | GB_GCA_002171125.1 | 68.5 | Marinisomatota | Marinisomatia | SCGC-AAA003-L08 | TMED161 | TMED161 | TMED161 sp002171125 | 0.006 | 0.991 | 0.009 | Periplasm | M4_strict |
| OM-RGC.v1.005883376 | GB_GCA_018675825.1 | 66.3 | Bacteroidota | Bacteroidia | Flavobacteriales | BACL11 | GCA-002715885 | GCA-002715885 sp018675825 | 0.005 | 0.959 | 0.041 | Periplasm | M4_strict |
| OM-RGC.v1.005916762 | GB_GCA_004213995.1 | 87.9 | Bacteroidota | Bacteroidia | Flavobacteriales | Flavobacteriaceae | SGZJ01 | SGZJ01 sp004213995 | 0.040 | 0.994 | 0.000 | Periplasm | M4_strict |
| OM-RGC.v1.005927835 | GB_GCA_002707765.1 | 88.3 | Marinisomatota | Marinisomatia | SCGC-AAA003-L08 | GCA-002707645 | GCA-2717625 | GCA-2717625 sp002707765 | 0.004 | 0.999 | 0.000 | Periplasm | M4_strict |
| OM-RGC.v1.005943084 | RS_GCF_003688405.1 | 77.0 | Bacteroidota | Bacteroidia | Flavobacteriales | Flavobacteriaceae | Ulvibacter | Ulvibacter antarcticus | 0.004 | 0.998 | 0.002 | Periplasm | M4_strict |
| OM-RGC.v1.005991260 | GB_GCA_002171125.1 | 66.7 | Marinisomatota | Marinisomatia | SCGC-AAA003-L08 | TMED161 | TMED161 | TMED161 sp002171125 | 0.004 | 0.993 | 0.006 | Periplasm | M4_strict |
| OM-RGC.v1.006019564 | GB_GCA_002171125.1 | 93.9 | Marinisomatota | Marinisomatia | SCGC-AAA003-L08 | TMED161 | TMED161 | TMED161 sp002171125 | 0.005 | 0.974 | 0.026 | Periplasm | M4_strict |
| OM-RGC.v1.006233121 | GB_GCA_029253805.1 | 94.6 | Bacteroidota | Bacteroidia | Flavobacteriales | Flavobacteriaceae | Hel1-33-131 | Hel1-33-131 sp029253805 | 0.010 | 0.989 | 0.002 | Periplasm | M4_strict |
| OM-RGC.v1.006302096 | GB_GCA_002171125.1 | 67.2 | Marinisomatota | Marinisomatia | SCGC-AAA003-L08 | TMED161 | TMED161 | TMED161 sp002171125 | 0.192 | 0.997 | 0.000 | Periplasm | M4_strict |
| OM-RGC.v1.006655193 | GB_GCA_018675825.1 | 66.1 | Bacteroidota | Bacteroidia | Flavobacteriales | BACL11 | GCA-002715885 | GCA-002715885 sp018675825 | 0.067 | 0.206 | 0.235 | Extracellular | M4_strict |

| | | | | | | | | | | | | | |
|---|---|---|---|---|---|---|---|---|---|---|---|---|---|
| OM-RGC.v1.007043509 | GB_GCA_002715565.1 | 95.3 | Bacteroidota | Bacteroidia | Flavobacteriales | BACL11 | GCA-2715565 | GCA-2715565 sp002715565 | 0.058 | 0.350 | 0.620 | Extracellular | M4_strict |
| OM-RGC.v1.007091679 | GB_GCA_913052445.1 | 97.0 | Bacteroidota | Bacteroidia | Flavobacteriales | Flavobacteriaceae | Hel1-33-131 | Hel1-33-131 sp913052445 | 0.016 | 0.995 | 0.000 | Periplasm | M4_strict |
| OM-RGC.v1.007259368 | GB_GCA_004213555.1 | 99.7 | Planctomycetota | Phycisphaerae | Phycisphaerales | SM1A02 | UBA8653 | UBA8653 sp002862275 | 0.005 | 0.332 | 0.667 | Extracellular | M4_strict |
| OM-RGC.v1.007292470 | GB_GCA_002700775.1 | 100.0 | Pseudomonadota | Gammaproteobacteria | Pseudomonadales | Pseudohongiellaceae | UBA9145 | UBA9145 sp002700775 | 0.005 | 0.002 | 0.997 | Extracellular | M4_strict |
| OM-RGC.v1.007508830 | GB_GCA_002684935.1 | 90.8 | Pseudomonadota | Gammaproteobacteria | Pseudomonadales | Pseudohongiellaceae | UBA9145 | UBA9145 sp002684935 | 0.004 | 0.002 | 0.997 | Extracellular | M4_strict |
| OM-RGC.v1.007535214 | GB_GCA_002726595.1 | 100.0 | Pseudomonadota | Gammaproteobacteria | Pseudomonadales | Pseudohongiellaceae | UBA9145 | UBA9145 sp002726595 | 0.004 | 0.022 | 0.978 | Extracellular | M4_strict |
| OM-RGC.v1.007543902 | GB_GCA_002724925.1 | 94.9 | Pseudomonadota | Gammaproteobacteria | Pseudomonadales | DSM-6294 | Oleibacter | Oleibacter sp002724925 | 0.005 | 0.091 | 0.908 | Extracellular | M4_strict |
| OM-RGC.v1.007632674 | GB_GCA_015664885.1 | 99.7 | Gemmatimonadota | Gemmatimonadetes | Longimicrobiales | UBA6960 | UBA1138 | UBA1138 sp003447875 | 0.005 | 0.500 | 0.499 | Periplasm | M4_strict |
| OM-RGC.v1.007966038 | GB_GCA_902579065.1 | 94.8 | Bacteroidota | Bacteroidia | Flavobacteriales | BACL11 | GCA-002715885 | GCA-002715885 sp902579065 | 0.059 | 0.893 | 0.019 | Periplasm | M4_strict |
| OM-RGC.v1.008045018 | GB_GCA_028122735.1 | 89.8 | Bacteroidota | Bacteroidia | Flavobacteriales | BACL11 | GCA-002715885 | GCA-002715885 sp028122735 | 0.055 | 0.901 | 0.077 | Periplasm | M4_strict |
| OM-RGC.v1.008047701 | GB_GCA_004213995.1 | 98.9 | Bacteroidota | Bacteroidia | Flavobacteriales | Flavobacteriaceae | SGZJ01 | SGZJ01 sp004213995 | 0.004 | 1.000 | 0.000 | Periplasm | M4_strict |
| OM-RGC.v1.008095885 | GB_GCA_002313925.1 | 98.9 | Gemmatimonadota | Gemmatimonadetes | Longimicrobiales | UBA6960 | UBA1142 | UBA1142 sp002313925 | 0.008 | 0.105 | 0.893 | Extracellular | M4_strict |
| OM-RGC.v1.008179074 | GB_GCA_002347035.1 | 98.2 | Gemmatimonadota | Gemmatimonadetes | Longimicrobiales | UBA6960 | UBA2982 | UBA2982 sp002347035 | 0.004 | 0.002 | 0.998 | Extracellular | M4_strict |
| OM-RGC.v1.008394547 | RS_GCF_900156675.1 | 87.2 | Pseudomonadota | Gammaproteobacteria | Pseudomonadales | DSM-6294 | Oleibacter | Oleibacter maritimus | 0.006 | 0.005 | 0.993 | Extracellular | M4_strict |
| OM-RGC.v1.008525034 | GB_GCA_002298335.1 | 100.0 | Pseudomonadota | Gammaproteobacteria | Pseudomonadales | DSM-6294 | Oleibacter | Oleibacter sp002298335 | 0.007 | 0.160 | 0.838 | Extracellular | M4_strict |
| OM-RGC.v1.008836064 | RS_GCF_030070105.1 | 94.5 | Pseudomonadota | Gammaproteobacteria | Pseudomonadales | DSM-6294 | Bermanella | Bermanella sp030070105 | 0.175 | 0.986 | 0.000 | Periplasm | M4_strict |
| OM-RGC.v1.009489323 | GB_GCA_943842425.1 | 60.8 | Bacteroidota | Bacteroidia | Flavobacteriales | UA16 | UBA974 | UBA974 sp002292405 | 0.026 | 0.958 | 0.012 | Periplasm | M4_strict |
| OM-RGC.v1.009511645 | GB_GCA_938082955.1 | 95.4 | Pseudomonadota | Gammaproteobacteria | Pseudomonadales | Pseudohongiellaceae | UBA9145 | UBA9145 sp938082955 | 0.895 | 0.000 | 0.005 | Cytoplasm | M4_strict |
| OM-RGC.v1.009626043 | GB_GCA_002724895.1 | 70.4 | Bacteroidota | Bacteroidia | Flavobacteriales | BACL11 | GCA-2693335 | GCA-2693335 sp002724895 | 0.312 | 0.001 | 0.270 | Cytoplasm | M4_strict |
| OM-RGC.v1.010390359 | GB_GCA_002691285.1 | 77.7 | Marinisomatota | Marinisomatia | Marinisomatales | UBA1611 | GCA-002708715 | GCA-002708715 sp002691285 | 0.011 | 0.978 | 0.018 | Periplasm | M4_strict |

| | | | | | | | | | | | | | |
|---|---|---|---|---|---|---|---|---|---|---|---|---|---|
| OM-RGC.v1.010667201 | GB_GCA_002696345.1 | 99.2 | Myxococcota | UBA4151 | UBA4151 | UBA4151 | UBA4151 | UBA4151 sp002696345 | 0.224 | 0.003 | 0.046 | Cytoplasm | M4_strict |
| OM-RGC.v1.010738341 | GB_GCA_002704685.1 | 85.6 | Marinisomatota | Marinisomatia | SCGC-AAA003-L08 | GCA-002707645 | GCA-002707645 | GCA-002707645 sp002704685 | 0.005 | 0.999 | 0.000 | Periplasm | M4_strict |
| OM-RGC.v1.010756458 | GB_GCA_002724895.1 | 94.4 | Bacteroidota | Bacteroidia | Flavobacteriales | BACL11 | GCA-2693335 | GCA-2693335 sp002724895 | 0.064 | 0.010 | 0.444 | Extracellular | M4_strict |
| OM-RGC.v1.010961368 | GB_GCA_002691285.1 | 78.8 | Marinisomatota | Marinisomatia | Marinisomatales | UBA1611 | GCA-002708715 | GCA-002708715 sp002691285 | 0.004 | 0.270 | 0.729 | Extracellular | M4_strict |
| OM-RGC.v1.011554844 | GB_GCA_002691285.1 | 92.9 | Marinisomatota | Marinisomatia | Marinisomatales | UBA1611 | GCA-002708715 | GCA-002708715 sp002691285 | 0.011 | 0.911 | 0.085 | Periplasm | M4_strict |
| OM-RGC.v1.011570699 | GB_GCA_002709695.1 | 93.3 | Poribacteria | WGA-4E | WGA-4E | UBA9662 | UBA9662 | UBA9662 sp002709695 | 0.420 | 0.003 | 0.303 | Cytoplasm | M4_strict |
| OM-RGC.v1.012011538 | GB_GCA_002684315.1 | 97.0 | Planctomycetota | Phycisphaerae | Phycisphaerales | SM1A02 | UBA8087 | UBA8087 sp002684315 | 0.505 | 0.000 | 0.047 | Cytoplasm | M4_strict |
| OM-RGC.v1.012320556 | GB_GCA_028823435.1 | 95.3 | Bacteroidota | Bacteroidia | Flavobacteriales | UA16 | UBA974 | UBA974 sp028823435 | 0.005 | 0.917 | 0.083 | Periplasm | M4_strict |
| OM-RGC.v1.012634301 | GB_GCA_002171125.1 | 94.3 | Marinisomatota | Marinisomatia | SCGC-AAA003-L08 | TMED161 | TMED161 | TMED161 sp002171125 | 0.004 | 0.932 | 0.067 | Periplasm | M4_strict |
| OM-RGC.v1.013434308 | RS_GCF_900156675.1 | 98.2 | Pseudomonadota | Gammaproteobacteria | Pseudomonadales | DSM-6294 | Oleibacter | Oleibacter maritimus | 0.908 | 0.000 | 0.047 | Cytoplasm | M4_strict |
| gene_1122877\|GeneMark.hmm\|309_aa\|-\|39284\|40213_543468 | RS_GCF_020025155.1 | 75.2 | Pseudomonadota | Gammaproteobacteria | Pseudomonadales | Pseudomonadaceae | Halopseudomonas | Halopseudomonas nanhaiensis | 0.004 | 0.999 | 0.001 | Periplasm | M5 |
| gene_1131781\|GeneMark.hmm\|315_aa\|+\|2891\|3838_586482 | GB_GCA_002354735.1 | 82.1 | Pseudomonadota | Gammaproteobacteria | Pseudomonadales | Oleiphilaceae | UBA2679 | UBA2679 sp002354735 | 0.012 | 0.910 | 0.084 | Periplasm | M5 |
| gene_1245712\|GeneMark.hmm\|290_aa\|+\|141\|1013_641712 | GB_GCA_913061415.1 | 98.6 | Pseudomonadota | Gammaproteobacteria | Pseudomonadales | DSM-6294 | Oleispira | Oleispira sp913061415 | 0.133 | 0.734 | 0.093 | Periplasm | M5 |
| gene_1276145\|GeneMark.hmm\|291_aa\|+\|565\|1440_627515 | GB_GCA_000967895.1 | 99.3 | Pseudomonadota | Gammaproteobacteria | Pseudomonadales | DSM-6294 | Oleispira | Oleispira antarctica | 0.558 | 0.918 | 0.000 | Periplasm | M5 |
| gene_1345533\|GeneMark.hmm\|314_aa\|-\|2351\|3295_661293 | GB_GCA_016763285.1 | 100.0 | Pseudomonadota | Gammaproteobacteria | Pseudomonadales | Pseudomonadaceae | Halopseudomonas | Halopseudomonas sp016763285 | 0.004 | 0.996 | 0.004 | Periplasm | M5 |
| gene_261593\|GeneMark.hmm\|316_aa\|-\|192\|1142_130008 | GB_GCA_016763285.1 | 92.4 | Pseudomonadota | Gammaproteobacteria | Pseudomonadales | Pseudomonadaceae | Halopseudomonas | Halopseudomonas sp016763285 | 0.004 | 0.905 | 0.095 | Periplasm | M5 |
| gene_48860\|GeneMark.hmm\|289_aa\|+\|641\|1510_25799 | GB_GCA_002840835.1 | 98.6 | Pseudomonadota | Gammaproteobacteria | Burkholderiales | Burkholderiaceae | Hydrogenophaga | Hydrogenophaga sp002840835 | 0.005 | 0.994 | 0.005 | Periplasm | M5 |
| gene_895782\|GeneMark.hmm\|254_aa\|-\|365\|1129_447244 | RS_GCF_002377945.1 | 98.4 | Pseudomonadota | Gammaproteobacteria | Pseudomonadales | Moraxellaceae | Psychrobacter | Psychrobacter sp002377945 | 0.023 | 0.126 | 0.785 | Extracellular | M5 |
| gene_970408\|GeneMark.hmm\|301_aa\|-\|34\|939_542835 | RS_GCF_942548115.1 | 58.0 | Actinomycetota | Actinomycetes | Propionibacteriales | Nocardioidaceae | Aeromicrobium | Aeromicrobium sp001423335 | 0.034 | 0.251 | 0.719 | Extracellular | M5 |

| | | | | | | | | | | | | | |
|---|---|---|---|---|---|---|---|---|---|---|---|---|---|
| NODE_122804_length_985_cov_16.2419_1 | RS_GCF_014652895.1 | 100.0 | Pseudomonadota | Gammaproteobacteria | Pseudomonadales | Moraxellaceae | Psychrobacter | Psychrobacter glaciei | 0.021 | 0.082 | 0.833 | Extracellular | M5 |
| NODE_1564_length_5726_cov_15.0628_1 | RS_GCF_904846075.1 | 98.0 | Pseudomonadota | Gammaproteobacteria | Pseudomonadales | Moraxellaceae | Psychrobacter | Psychrobacter glacincola_C | 0.034 | 0.552 | 0.265 | Periplasm | M5 |
| NODE_18136_length_3583_cov_6.01162_4 | GB_GCA_009497895.1 | 96.5 | Pseudomonadota | Gammaproteobacteria | Pseudomonadales | Pseudomonadaceae | Halopseudomonas | Halopseudomonas pelagia_A | 0.672 | 0.110 | 0.094 | Cytoplasm | M5 |
| NODE_43566_length_1995_cov_7.65464_1 | GB_GCA_000967895.1 | 98.8 | Pseudomonadota | Gammaproteobacteria | Pseudomonadales | DSM-6294 | Oleispira | Oleispira antarctica | 0.675 | 0.468 | 0.000 | Cytoplasm | M5 |
| NODE_567_length_29609_cov_12.2345_1 | GB_GCA_011053055.1 | 100.0 | Pseudomonadota | Gammaproteobacteria | Pseudomonadales | Pseudomonadaceae | Halopseudomonas | Halopseudomonas sabulinigri_A | 0.768 | 0.182 | 0.000 | Cytoplasm | M5 |
| OM-RGC.v1.002390589 | RS_GCF_016863515.1 | 98.8 | Actinomycetota | Actinomycetes | Mycobacteriales | Micromonosporaceae | Micromonospora | Micromonospora gifhornensis | 0.004 | 1.000 | 0.000 | Periplasm | M5 |
| OM-RGC.v1.004916997 | RS_GCF_003671495.1 | 83.0 | Pseudomonadota | Gammaproteobacteria | Pseudomonadales | Ketobacteraceae | Ketobacter | Ketobacter sp003671495 | 0.040 | 0.282 | 0.694 | Extracellular | M5 |
| OM-RGC.v1.006047268 | RS_GCF_002903165.1 | 96.0 | Pseudomonadota | Gammaproteobacteria | Pseudomonadales | Pseudomonadaceae | Halopseudomonas | Halopseudomonas oceani | 0.006 | 0.993 | 0.007 | Periplasm | M5 |
| OM-RGC.v1.006912781 | RS_GCF_000244955.1 | 99.7 | Actinomycetota | Actinomycetes | Mycobacteriales | Pseudonocardiaceae | Saccharomonospora_A | Saccharomonospora_A marina | 0.007 | 0.449 | 0.550 | Extracellular | M5 |

**Supplementary Table S3.** Nucleotide and amino acid sequences of selected enzymes MetaG1-MetaG9. The additional carbohydrate binding module of MetaG4 was removed.

| Enzyme | Nucleotide (nt) and amino acid (aa) sequence |
|---|---|
| MetaG1 (nt) | ATGGACTCACCTTATCAACACGGACCAGATCCCACCTATAGCAGCATCGGCGGGGTGGGTCCGTATGCT<br>ACCGCACAGGTTTCCGTTCCGGCGTCTGGCACCCCGGGTTTCGGCAGCGGTACAATCTATTACCCGACC<br>ACGACTACGGAAGGTCGTTTTGGTGGCATTGCGATCAGCCCGGGCTACGGTGGCGCCGAATCCTCTATC<br>GGTTGGTATGGTCCGCGTCTGGCGTCACACGGCTTCGTGGTGATTACCATCGAAACCATCAGCCGTTAC<br>GAAGATCCGTCTGCTCGCGCAGATGAGCTCCAAGCGGCCTTGGACTGGTTGGTTGCTTCCAGCACCGTC<br>AAAGATCGTGTGGATGGTAGCCGTCTGGCGGTTATGGGTCATAGCATGGGTGGCGGTGGCGCGCTGGAG<br>GCAGCGAGAGATAATCCGTCGCTGAAGGCTGCGGTGCCATTGACCCCGTGGAATGCATACGAGAGCTTT<br>AGCACGATGACCGTCCCGACCCTGATTTTCGGCGCCGATGACGACAGCATCGCCCCTGCGAACTATCAT<br>GCGATTCCGTTTTATTGGAACATTCCGAGTTCAACGCGTAAAGCGTACCTGGAGCTGCGTTACGCAAAC<br>CACTTTACCCCAAATTTTCCGCACAGCGAAATTACTCGCTACAGCGTGTCCTGGTTAAAGCGCTGGGTT<br>GACCTGGACACCCGCTACGACGAGTTCATCTGTCCGGGTCCGTGGTGGAACTGGGAATTCACCCATTCC<br>TGGAACAACTGCCCGTAA |
| MetaG1 (aa) | MGSSHHHHHHSSGENLYFQGHMDSPYQHGPDPTYSSIGGVGPYATAQVSVPASGTPGFGSGTIYYPTTT<br>TEGRFGGIAISPGYGGAESSIGWYGPRLASHGFVVITIETISRYEDPSARADELQAALDWLVASSTVKD<br>RVDGSRLAVMGHSMGGGGALEAARDNPSLKAAVPLTPWNAYESFSTMTVPTLIFGADDDSIAPANYHAI<br>PFYWNIPSSTRKAYLELRYANHFTPNFPHSEITRYSVSWLKRWVDLDTRYDEFICPGPWWNWEFTHSWN<br>NCP |
| MetaG2 (nt) | ATGTGTGCAGCTCAAGCGCAGACATCACCACCCACTTCAGCCTCCTTGAACGCGACGGCTGGTCCGTTG<br>AGCGTTTCTACGAGCTCTGTTTCCAGCTGGGCGGCGCGTGGTTTTGGTGGCGGTACAATTTATTATCCG<br>AATGCGACTGGTCGTTACGGCGTTGTTGCGATTAGCCCGGGGTATACCGCTCGTCAGAGCAGCATTGCG<br>TGGCTGGGTCGCCGTCTCGCGACCCACGGTTTCGTGGTGATCACCATTGACACCAATTCGACCCTGGAC<br>CAGCCACCGAGCAGAGCCACGCAACTGATGGCAGCGCTGAATCATGTTGTAAATAACGCAAATGCGACG<br>GTGCGTAGCCGCGTGGACGCATCCAAGCTGGCGGTTGCAGGCCATTCTATGGGTGGAGGCGGCAGCCTG<br>ATCGCGGCGGAGAACAACCCGTCTTTGAAAGCTGCGTACCCGCTGACCCCTTGGAGCGTCTCCAAAAAC<br>TATAGCAGCGTCTGTGTTCCGACCATGATCATCGGCGCTGATGGTGATTCCATCGCCAGCGTGTCGACC<br>CACAGCCGTTTGTTTTATAACTCCTTATCCTCTAATGTGTCCAAGGCATACGGCGAACTGAACAACGCC<br>AGCCATTTCACCCCGAACACCACCAACACCCCCATCGGCCGCTACGCTGTGACCTGGATGAAACGTTTT<br>GTTGACAGCGATACCCGTTACAGTCCGTTCCTGTGCGGTGCTCCGCACGATAGCTACGCGACTCGCAGC<br>GTGTTCGACCGTTACGAGGATAACTGCGCATACTAA |
| MetaG2 (aa) | MGSSHHHHHHSSGENLYFQGHMCAAQAQTSPPTSASLNATAGPLSVSTSSVSSWAARGFGGGTIYYPNA<br>TGRYGVVAISPGYTARQSSIAWLGRRLATHGFVVITIDTNSTLDQPPSRATQLMAALNHVVNNANATVR<br>SRVDASKLAVAGHSMGGGGSLIAAENNPSLKAAYPLTPWSVSKNYSSVCVPTMIIGADGDSIASVSTHS<br>RLFYNSLSSNVSKAYGELNNASHFTPNTTNTPIGRYAVTWMKRFVDSDTRYSPFLCGAPHDSYATRSVF<br>DRYEDNCAY |
| MetaG3 (nt) | ATGCAAGAAAATCCCTATGAGAGGGGACCAGATCCAACGGAAGATTCGATCGAGGCCGTGCGTGGCCCG<br>TTCAGCGTAGCCGAGCAAAACGTGAGCTCCTTGACCCCGGGTTTCGGTGGCGGCACCATTTACTACCCG<br>ACGGATACCTCTGAGGGTACGTTCGGCGCGGTCGCGGTGGCTCCGGGTTATACCGCATCTGAAAGCAGC<br>ATGAGCTGGTATGGTCCGCGTATTGCTAGCCAGGGTTTTGTTGTTTTTACCATCGACACTAATACGCGT<br>TACGACCAACCGGGTAGCCGTGGCGATCAGCTGTTGGCAGCGCTGGACTACCTGGTTGAAGATGCTCCT<br>TTGGCGGTGCGCTCCCGCATTGACCCGGATAGACTGGGTGTTATGGGTCATTCCATGGGTGGAGGCGGC<br>GCGCTGGAAGCATCTGCAGATCGTCCGTCGCTTCAGGCAGCGATCCCGTTGACCCCGTGGAATCTGGAC<br>AAGACCTGGTCAGAGATTCGTGTTCCGACCTTCATCATTGGTGCTGAGAACGATAGCATCGCGAGCGTG<br>CGTACCCACGCGGAGCCGTTTTACGAGTCCCTGCCGGCTACCCTGGACAAGGCGTATCTGGAACTCAAC<br>GGCGCGACCCACTTTGCTCCGAACACCTCCAATACGACCATTGCCAAATACAGCATCAGCTGGCTGAAA<br>CGTTTCATCGACAACGACACTCGCTATGAACAATTTCTGTGCCCCGCGCCATCTGGCTTCGCCATCGAG<br>GAATATCGCGAAACCTGTCCGTACACCGGTTAA |

| | |
|---|---|
| MetaG3<br>(aa) | MGSSHHHHHHSSGENLYFQGHMQENPYERGPDPTEDSIEAVRGPFSVAEQNVSSLTPGFGGGTIYYPTD<br>TSEGTFGAVAVAPGYTASESSMSWYGPRIASQGFVVFTIDTNTRYDQPGSRGDQLLAALDYLVEDAPLA<br>VRSRIDPDRLGVMGHSMGGGGALEASADRPSLQAAIPLTPWNLDKTWSEIRVPTFIIGAENDSIASVRT<br>HAEPFYESLPATLDKAYLELNGATHFAPNTSNTTIAKYSISWLKRFIDNDTRYEQFLCPAPSGFAIEEY<br>RETCPYTG |
| MetaG4<br>(nt) | ATGGCTGATAATCCCTATGAAAGGGGACCCGACCCGACGCGTACCTCCGTGGCGACCGAGCGCGGTCCG<br>TTCGCCAATACCTCCGTGAGCGTGCCGACCGGTTACGGCTTCAATGGTGGTCGTATTTATTACCCGACC<br>GATACCTCGCAAGGCACCTTTGGTGCGATCGCCATCAGCCCGGGTTACACCGCGTTGTTTAGCGCAGAG<br>TTGGCTTGGATGGGCCCTTGGCTGGCCAGCCACGGCTTCGTGGTCATTGGCATTGAAACCAATTCACGC<br>AACGACTTTGATACCGCTCGTGGTACTCAACTGCTGGCTGCGTTGGACTACCTGACCCAGCAAAGCCCG<br>GTTCGTGATCGTGTTGACGCGAGCCGTCTCGCTGTGGCGGGTCATAGCATGGGTGGCGGCGGGGCATTA<br>AGCGCGGCGACGAGACGTCCGGCGTTGAAGGCGGCAGTAGGCATCACCCCGTTCTCTCCGTCGAGCAAC<br>CTGGCGAACGACCGCGTTCCAACGATGGTGATCAGTGGCCAGGCAGATACGGTTGTAACTCCGTCTTAT<br>GCACTTGACCTGTATAACTCCCTGCCGAGCACCACCGAGAGCGTCTACGTGGAAGTTGCGGGTGGTGAT<br>CACGGCTTCATGGTTGGTCGCTCCAATCCGGTGATGATTCGTACCATGCTGCCATTCCTGAAAATCTTT<br>GTTGACAACGACGCGCGTTATAGCCAGTTTCTGTGTCCGCTGATGGATAACTCCGGTGTTGTGACCTAC<br>CGCTCTACGTGCCCGCTGATTAACTAA |
| MetaG4<br>(aa) | MGSSHHHHHHSSGENLYFQGHMADNPYERGPDPTRTSVATERGPFANTSVSVPTGYGFNGGRIYYPTDT<br>SQGTFGAIAISPGYTALFSAELAWMGPWLASHGFVVIGIETNSRNDFDTARGTQLLAALDYLTQQSPVR<br>DRVDASRLAVAGHSMGGGGALSAATRRPALKAAVGITPFSPSSNLANDRVPTMVISGQADTVVTPSYAL<br>DLYNSLPSTTESVYVEVAGGDHGFMVGRSNPVMIRTMLPFLKIFVDNDARYSQFLCPLMDNSGVVTYRS<br>TCPLIN |
| MetaG4-CBM<br>(nt) | ATGGCTGATAATCCCTATGAAAGGGGACCCGACCCGACGCGTACCTCGGTGGCCACCGAGCGCGGTCCG<br>TTCGCAAATACCTCCGTGTCAGTCCCGACGGGTTATGGTTTCAACGGCGGCCGTATTTACTACCCGACC<br>GATACTTCTCAAGGTACGTTTGGAGCGATCGCCATCTCCCCTGGTTACACTGCGTTGTTTAGCGCTGAA<br>TTGGCGTGGATGGGTCCGTGGCTGGCGAGCCATGGTTTCGTGGTCATCGGCATTGAAACCAACAGCCGT<br>AACGACTTCGACACCGCGCGTGGTACGCAGTTGCTGGCTGCGCTGGATTATCTGACCCAGCAAAGCCCT<br>GTTCGTGATCGTGTGGACGCGTCCAGACTGGCGGTGGCGGGTCATAGCATGGGCGGTGGCGGCGCCTTA<br>AGCGCTGCGACCCGCCGTCCGGCACTTAAGGCTGCTGTGGGCATCACCCCGTTCAGCCCAAGCAGCAAC<br>TTGGCGAATGACCGTGTTCCGACCATGGTGATCTCTGGTCAGGCGGATACCGTTGTCACCCCGTCCTAT<br>GCATTGGACCTCTACAATTCTCTGCCGAGCACCACTGAGAGCGTGTATGTTGAGGTGGCGGGTGGGGAT<br>CATGGCTTCATGGTTGGTCGCTCCAACCCGGTTATGATCCGTACCATGCTGCCGTTCCTGAAGATCTTT<br>GTTGACAACGACGCACGCTATTCTCAGTTTCTGTGTCCGCTGATGGATAACAGCGGCGTTGTGACCTAT<br>CGTAGCACGTGCCCGCTGATTAACACCCCGACCACACCGCCGCCGACAAGTCCGCCGCCTACGAGCCCG<br>CCGCCAACGTCGCCGCCACCAACCACCCCGCCGCCGGGTAGCGCAACCCAAATTGTTGGCGTGCAATCC<br>AATCGTTGTATTGATGTGCCCAACTCGAGCCGCAACAACGGCACTCGCGTACAATTGTACGATTGCCAC<br>GGCCAGACCAATCAGGCATGGACCTACACCGCGAACAAACAGCTCCAAGTTTACGGCAACATGTGTCTG<br>GATGCGGCAGGTACGGGTAACGGTGCCGCGGTTCAGATCTACAGCTGCCACGGCGGCGCGAACCAACAG<br>TGGAATGTTAATAGCAATGGCACCATTAGCGGTGTCCAGTCTGGCCGCTGCCTGGACGTGTGGTCAACC<br>TCTAACGGTGCTCAGGTTCAGCTGTACGACTGCCACGGCGGTACTAATCAACGTTTTAATCTGGTTGCG<br>CGTTAA |
| MetaG4-CBM<br>(aa) | MGSSHHHHHHSSGENLYFQGHMADNPYERGPDPTRTSVATERGPFANTSVSVPTGYGFNGGRIYYPTDT<br>SQGTFGAIAISPGYTALFSAELAWMGPWLASHGFVVIGIETNSRNDFDTARGTQLLAALDYLTQQSPVR<br>DRVDASRLAVAGHSMGGGGALSAATRRPALKAAVGITPFSPSSNLANDRVPTMVISGQADTVVTPSYAL<br>DLYNSLPSTTESVYVEVAGGDHGFMVGRSNPVMIRTMLPFLKIFVDNDARYSQFLCPLMDNSGVVTYRS<br>TCPLINTPTTPPPTSPPPTSPPPTSPPPTTPPPGSATQIVGVQSNRCIDVPNSSRNNGTRVQLYDCHGQ<br>TNQAWTYTANKQLQVYGNMCLDAAGTGNGAAVQIYSCHGGANQQWNVNSNGTISGVQSGRCLDVWSTSN<br>GAQVQLYDCHGGTNQRFNLVAR |
| MetaG5<br>(nt) | ATGCCCTCAGTAAGTTATCTAGAAGCAGCTAGGGGAACCTATTCTGTTCGTACCAGCCGCGTTTCCTCG<br>TTCGTATCCGGTTTCGGCGGCGGCACGATCCACTACCCGACTGGTACCACCGGTACGATGGCAGCAATC<br>GTGGTCATCCCGGGTTTCGTCTCTGCGGAGAGCAGCATTGAATGGTGGGGTCCGAAACTCGCGAGCTAT<br>GGTTTCGTGGTGTTGACCATCGACACAAATTCCGGTTTCGATCAACCGCCTAGCCGCGCGCGTCAGATC<br>AACAACGCGCTGGACTACCTGGTTTCGCAAAACACCTCAAGACGTAGCGCTGTGCAGGGTATGATTGAT<br>ACCGATCGTCTGGGTGTTGTTGGTTGGTCCATGGGCGGCGGCGGCACCCTGCGCGTGGCCGAGGAGGGT |

| | |
|---|---|
| | CGTATTAAGGCTGCGATCCCGCTGGCTCCGTGGGACACTACGAACTTTCGTGATAATTATACCCCGACC<br>CTGATTTTTGCCTGCCAAAGCGACATCATCGCGCCGGTGTATCAGCATGCAAGCCCGTTTTACAACCAG<br>ATTCCGAACTCCACCGATAAGGCGTTTGTGGAACTGCGCGGCGGCAGCCACTACTGCGGTAATGGTGGC<br>GGCATTTATAACGATGTTTTGGGGCGTTTAGGCGTCTCTTGGATGAAACGTCATTTGGACCAAGATACC<br>CGTTACCAGCAGTTTCTGTGCGGTCCAAACCACGAAAGCGACAGCCAGGTTTCTGACTACCGCGGTAAT<br>TGTCAATAA |
| MetaG5 (aa) | MGSSHHHHHHSSGENLYFQGHMPSVSYLEAARGTYSVRTSRVSSFVSGFGGGTIHYPTGTTGTMAAIVV<br>IPGFVSAESSIEWWGPKLASYGFVVLTIDTNSGFDQPPSRARQINNALDYLVSQNTSRRSAVQGMIDTD<br>RLGVVGWSMGGGGTLRVAEEGRIKAAIPLAPWDTTNFRDNYTPTLIFACQSDIIAPVYQHASPFYNQIP<br>NSTDKAFVELRGGSHYCGNGGGIYNDVLGRLGVSWMKRHLDQDTRYQQFLCGPNHESDSQVSDYRGNCQ |
| MetaG6 (nt) | ATGGCTTCAAATCCCCCTCCCGACCCAGTAGATCCAGGACAACCGAGTGGCTTCGAGCGCGGTCCGAAT<br>CCGACCTTGTCCCTGGTCGAGGCGGATAGAGGCCCGTTTAGCGTTCGTAGCTCCCGCGTGAGTGGTCTC<br>GTTTCCGGCTTTGGTGGCGGTACGATTCATTACCCGACCAATACTACAGGCACCATGGCAGCCGTTGTT<br>GTCATCCCGGGTTATATTTCGGCGGAATCAAGCATTGAATGGTGGGGTCCAAAACTGGCGAGCCATGGT<br>TTCGTGGTGATGACCATCGACACGAATACCGGTTTCGACCAACCTCCGAGCCGCGCGCGTCAGATTAAC<br>AGCGCGCTGGACTACCTGGTCGATCAAAACACCGAGCGCAACGGCGACGTTGAGGGTATGATTGACACC<br>GATCGTTTGGGCGTGATCGGCTGGAGCATGGGTGGCGGCGGCACCTTACGTGTGGCTACCGAAGGTCGA<br>ATTAAAGCGGCAATTCCGCTGGCTCCGTGGGATACCAGCTCTTGGCAGTTTCGTAACGTGCAGGCGCCG<br>ACGATGATCATCGCCTGCGAATCTGATATCGTGGCCCCCGTTGGTAGCCATGCTTCGCCGTTCTACAAT<br>CGTCTTCCGGGTGACATCAACAAGGCGTTTGTTGAACTGGATGGTGGCAACCACTACTGCGGTAACGGT<br>GGCGCATCTTTTGGCCAGTATGATACTGTACTGAGCCGTTTGGGTGTTTCCTGGATGAAGCGCCACCTG<br>GACAATGATACCCGTTACAGCCAGTTCCTGTGTGGTCCGAACCACGAAAGCGACCGTGAGATCAGCGAG<br>TATCGTGGCAACTGCCCGTATTAA |
| MetaG6 (aa) | MGSSHHHHHHSSGENLYFQGHMASNPPPDPVDPGQPSGFERGPNPTLSLVEADRGPFSVRSSRVSGLVS<br>GFGGGTIHYPTNTTGTMAAVVVIPGYISAESSIEWWGPKLASHGFVVMTIDTNTGFDQPPSRARQINSA<br>LDYLVDQNTERNGDVEGMIDTDRLGVIGWSMGGGGTLRVATEGRIKAAIPLAPWDTSSWQFRNVQAPTM<br>IIACESDIVAPVGSHASPFYNRLPGDINKAFVELDGGNHYCGNGGASFGQYDTVLSRLGVSWMKRHLDN<br>DTRYSQFLCGPNHESDREISEYRGNCPY |
| MetaG7 (nt) | ATGCCCTCAGTAAGTTTTCTAGAAGCAGCTACAGGACCGTATAGCGTTGATACCGAGCGCGTGTCCGGC<br>CTGGTTTCCGGTTTCGGCGGTGGTACGATTCATTACCCGGAGGACACCACCGGTACGATGGCGGCGATC<br>GTGGTTATTCCGGGTTATGTTTCTGCTGAATCTAGCATTGAATGGTGGGGTCCGAAATTGGCTAGCCAC<br>GGCTTCGTGGTGATGACCATCGATACCAATAGCGGCTTCGATCAGCCTCCGAGCAGAGCTAATCAGATT<br>AACGCAGCGCTGGACTACCTTATCGACGAGAACACCGCATTCGGCTCGCCGGTTCAAGGTATGATTGAC<br>ACGGATCGTCTGGGTGTAGTCGGTTGGTCAATGGGTGGCGGCGGCACCCTGCGCGTGGCGACCGAAGGT<br>CGTATTAGCGCTGCGATCCCGTTGGCCCCATGGGACACTTCTAGCTTTCGTAATGTTGAGGCACCGACC<br>CTCATCATCGCCTGTGAGAGCGATATCATCGCCCCAGTTGGCTCCCACGCGAGCCCGTTTTATAACCGC<br>ATCCCGGAGGACGTGGACAAGGCGTTCGTCGAGATCGATGGGGGCAGTCACTACTGCGGCAACGGTGGT<br>GGGTTCAACAACGATGTGCTGAGCCGTTTTGGCGTCTCCTGGATGAAATTGCACCTGGATAATGACGCG<br>CGTTACGAACAATTTCTGTGCGGTCCGGATCATGAAGACGACAGCGACATTTCGGAATACCGTGGTAAC<br>TGCCCGTAA |
| MetaG7 (aa) | MGSSHHHHHHSSGENLYFQGHMPSVSFLEAATGPYSVDTERVSGLVSGFGGGTIHYPEDTTGTMAAIVV<br>IPGYVSAESSIEWWGPKLASHGFVVMTIDTNSGFDQPPSRANQINAALDYLIDENTAFGSPVQGMIDTD<br>RLGVVGWSMGGGGTLRVATEGRISAAIPLAPWDTSSFRNVEAPTLIIACESDIIAPVGSHASPFYNRIP<br>EDVDKAFVEIDGGSHYCGNGGGFNNDVLSRFGVSWMKLHLDNDARYEQFLCGPDHEDDSDISEYRGNCP |
| MetaG8 (nt) | ATGTCAAATTGTTATCAAAGGGGACCCAACCCAACCGTTTCGGCCTTGGAGGCGGACAGCGGCCCGTAC<br>TCAGTTCGTACCATTAACGTTTCCTCTTGGGTTTCTGGTTTCGGCGGTGGCACGATCCACTACCCGGTG<br>GGTACTGAGGGTACCATGGGCGCGATTGCTGTGATCCCTGGTTACGTTAGTTATGAACGTTCCATTAAA<br>TGGTGGGGTCCGCGTTTGGCGAGCTGGGGCTTCGTGGTGATCACCACTGATACCAATACCATCTACGAC<br>CAACCGGATAGCCGCGCGGACCAGCTGAGCGCGGCTCTGGACTATGTCATTAGCCAAAGCAATAGCTCC<br>CGTAGCCCGATTTACGGCATGGTTGATGCCAACCGTCTCGGCGCTATGGGTTGGAGCATGGGAGGCGGT<br>GGCACGCTGAAGCTGTCTACCGAACGTGAATTGAAGGCGGCGATTCCGCAGGCACCGTACTACGCAGGC<br>TTCAACCCGTTCGATGAGATCACAACCCCGACCCTGATTATCGCGTGTGAGCTGGACGTCGTGGCGCCA<br>GTGGCTCAGCATGCGTCCCCGTTCTATCGCGAGATCCCGGGCTCGACCGCAAAAGCCTTTCTGGAAATC |

AACGGTGGTGATCACTTTTGCGCCAACAGCGGCTACCCGGACGAGGACATCCTGGGTAAATATGGTATT
GCTTGGATGAAGCGCTTTATCGATGAGGATCGCCGTTATGACCAATTTCTTTGCGGTCCGAATCATGAA
GCAGACAGATCTATCAGCGAATATCGTGATACGTGCAACTATTAA

MetaG8 (aa)

MGSSHHHHHHSSGENLYFQGHMSNCYQRGPNPTVSALEADSGPYSVRTINVSSWVSGFGGGTIHYPVGT
EGTMGAIAVIPGYVSYERSIKWWGPRLASWGFVVITTDTNTIYDQPDSRADQLSAALDYVISQSNSSRS
PIYGMVDANRLGAMGWSMGGGGTLKLSTERELKAAIPQAPYYAGFNPFDEITTPTLIIACELDVVAPVA
QHASPFYREIPGSTAKAFLEINGGDHFCANSGYPDEDILGKYGIAWMKRFIDEDRRYDQFLCGPNHEAD
RSISEYRDTCNY

MetaG9 (nt)

ATGATAACGGCTACAGAACTAACAAAAACTAGGGACAACGGCCCTTTTTCCGTGAAGAGCAAGCACGTG
TCTCGCCAGAGCGCTAATGGCTTTGGTGGCGGCACCATCCACTACCCGACGGATGCGGGTAGCTGCGGT
CTGCTGGGTGGTATTGCCGTTGTCCCGGGTTACGTTTCGTACGAGAGCTCCATTAAATGGTGGGGTCCG
CGCCTGGCGAGCTGGGGCTTCGTGGTGATTACCATTAACACCAGTTCGATCTACGACAATCCGGACAGC
CGTGCAGAACAGCTGTCGGCGGCACTGGATCATCTGATTGCGGACAAGACCGTTGGTCACATGATCGAC
CCGAATCGTCTTGGTGCCATCGGGTGGTCTATGGGTGGCGGCGGCGCGTTACGTTTGGCTACCGAACGT
AGCACCGTCCAGGCAATTATCGCGCAAACCCCGTATCACGACACGAGCTACGGCGCTATGGATACTCCG
GCATTGTTTATCGCGTGTGAAAATGATCGTATCGCTCCGAACAAAAAGTACACCAATATCTTTTATCAA
AAAGCCGATGGTCCAAAAATGAAAGTTGAAATTAACAACGGCTCTCACTTCTGCGCCAGCCATCGTTTC
AACGAGAAACTGCTGAGCAAGCCGGCGATCGCGTGGATGCAACGTTATATCAACGGTGACACTCGCTTC
GACAAGTTCCTGTGTGGAAACGAGTCCTACATTGATGATCCGCGCATCTCCGCGTATGACTATGAGGAT
TGCTTGTAA

MetaG9 (aa)

MGSSHHHHHHSSGENLYFQGHMITATELTKTRDNGPFSVKSKHVSRQSANGFGGGTIHYPTDAGSCGLL
GGIAVVPGYVSYESSIKWWGPRLASWGFVVITINTSSIYDNPDSRAEQLSAALDHLIADKTVGHMIDPN
RLGAIGWSMGGGGALRLATERSTVQAIIAQTPYHDTSYGAMDTPALFIACENDRIAPNKKYTNIFYQKA
DGPKMKVEINNGSHFCASHRFNEKLLSKPAIAWMQRYINGDTRFDKFLCGNESYIDDPRISAYDYEDCL

PHL7 (nt)

ATGGCGAACCCGTACGAGCGCGGGCCCGATCCCACCGAGTCGAGCATCGAGGCCGTCCGCGGGCCGTTC
GCCGTGGCCCAGACGACGGTGTCGAGGCTCCAGGCCGACGGCTTCGGCGGCGGGACCATCTACTACCCG
ACCGACACGAGCCAGGGCACCTTCGGTGCGGTGGCGATCTCGCCGGGGTTCACGGCGGGCCAGGAGAGC
ATCGCCTGGCTCGGCCCCCGCATCGCGTCGCAGGGCTTCGTGGTGATCACGATCGACACGATCACGCGC
CTCGACCAGCCCGACAGCCGGGGTCGCCAGCTGCAGGCCGCGCTCGACCACCTGCGCACCAACAGCGTC
GTGCGCAACCGGATCGACCCGAACCGGATGGCGGTCATGGGCCACTCGATGGGCGGCGGCGGGGCGCTG
TCCGCCGCGGCGAACAACACGAGCCTCAAAGCCGCCATCCCGCTGCAGGGCTGGCACACCCGGAAGAAC
TGGTCGAGCGTGCGGACGCCGACCCTGGTGGTCGGGGCCCAGCTCGACACCATCGCGCCGGTGAGCTCG
CACTCGGAGGCCTTCTACAACAGCCTGCCGAGCGACCTCGACAAGGCGTACATGGAGCTCCGCGGGGCC
AGCCACCTCGTGTCGAACACGTATGACACGACGCTGGCGAAGTACAGCATCGCCTGGCTCAAGCGGTTC
GTCGACGACGACCTCCGCTACGAGCAGTTCCTGTGCCCGGCGCCGGACGACTTCGCGATCTCCGAGTAC
CGCTCCACCTGCCCGTTC

PHL7 (aa)

MANPYERGPDPTESSIEAVRGPFAVAQTTVSRLQADGFGGGTIYYPTDTSQGTFGAVAISPGFTAGQES
IAWLGPRIASQGFVVITIDTITRLDQPDSRGRQLQAALDHLRTNSVVRNRIDPNRMAVMGHSMGGGGAL
SAAANNTSLKAAIPLQGWHTRKNWSSVRTPTLVVGAQLDTIAPVSSHSEAFYNSLPSDLDKAYMELRGA
SHLVSNTYDTTLAKYSIAWLKRFVDDDLRYEQFLCPAPDDFAISEYRSTCPFLEHHHHHH

LCC$^{ICCG}$ (nt)

ATGAGCAACCCGTACCAGCGTGGCCCGAATCCGACCCGCAGCGCACTGACCGCAGATGGCCCGTTTAGC
GTGGCAACCTACACCGTCTCACGCCTGTCAGTCTCGGGTTTTGGCGGTGGCGTGATTTATTACCCGACC
GGCACGTCTCTGACGTTCGGTGGCATCGCGATGAGTCCGGGTTATACCGCAGATGCTAGCTCTCTGGCA
TGGCTGGGTCGTCGCCTGGCTTCCCATGGCTTTGTGGTTCTGGTGATTAACACGAATTCACGTTTCGAT
GGCCCGGACAGCCGCGCCTCTCAGCTGAGTGCCGCCCTGAACTACCTGCGTACCAGTTCCCCGAGCGCC
GTTCGCGCACGTCTGGATGCAAATCGTCTGGCGGTTGCCGGTCATTCTATGGGTGGCGGTGGCACCCTG
CGTATTGCAGAACAAAACCCGAGCCTGAAAGCGGCTGTCCCGCTGACCCCGTGGCACACCGATAAAACG
TTTAATACCAGTGTCCCGGTGCTGATTGTTGGCGCAGAAGCTGACACCGTGGCGCCGGTTTCGCAGCAT
GCCATCCCGTTTTATCAAAACCTGCCGAGCACCACGCCGAAAGTTTACGTCGAACTGTGCAACGCATCG
CACATTGCTCCGAATAGCAACAATGCGGCCATTTCCGTTTATACGATCTCATGGATGAAACTGTGGGTC
GATAATGACACCCGTTACCGCCAGTTCCTGTGTAATGTGAACGACCCGGCTCTGTGCGACTTCCGCACC
AATAATCGCCACTGCCAA

| | |
|---|---|
| LCC$^{ICCG}$ (aa) | MSPGYTADASSLAWLGRRLASHGFVVLVINTNSRFDGPDSRASQLSAALNYLRTSSPSAVRARLDANRL AVAGHSMGGGGTLRIAEQNPSLKAAVPLTPWHTDKTFNTSVPVLIVGAEADTVAPVSQHAIPFYQNLPS TTPKVYVELCNASHIAPNSNNAAISVYTISWMKLWVDNDTRYRQFLCNVNDPALCDFRTNNRHCQLEHH HHHH |

MGSSHHHHHHSSGENLYFQGH = N-terminal His-tag + TEV cleavage site (from plasmid pET28a-TEV).

LEHHHHHH = C-terminal His-tag.

TAA or TGA = Stop codon.

**Supplementary Table S4. Known and partially characterized MHETases.** Names of MHETases used in this work as references according to the National Center for Biotechnology Information (NCBI) database. These sequences were used for the construction of the HMM for the search of putative MHETases.

| Sequence no. | NCBI entry | Organism | Reference |
|---|---|---|---|
| 1 | A0A0K8P8E7.1 | *Ideonella sakaiensis* 201-F6 | [1] |
| 2 | WP_083293388.1 | *Hydrogenophaga sp.* PML113 | [2] |
| 3 | WP_080747404.1 | *Comamonas thiooxydans* | [2] |

**Supplementary Table S5. Known and partially characterized TPA degrading enzymes and their homologues.** Names of the TPA-degrading enzymes used in this work as references according to the National Center for Biotechnology Information (NCBI) database. These sequences were used for the construction of the HMM to search for TPA-degrading enzymes. Some of these sequences are homologous due to the lack of enzymes for HMM generation.

| Name of TPA degrading enzyme* | NCBI entry | Organism | Reference |
|---|---|---|---|
| Terephthalate 1,2-dioxygenase, reductase component (TphA1) | BAE47088.1 | *Comamonas sp. E6* | [3] |
| | WP_034369195.1 | *Delftia tsuruhatensis sp.* | [4] |
| | WP_054022750.1 | *Ideonella sakaiensis* 201-F6 | [1] |
| | ABE33246.1 | *Paraburkholderia xenovorans LB400* | [5] |
| | WP_011599112.1 | *Rhodococcus jostii RHA1* | [6] |
| | AAR90190.1 | *Rhodococcus sp. DK17* | [7] |
| | AAX18944.1 | *Comamonas testosteroni* | [8] |
| Terephthalate 1,2-dioxygenase, terminal oxygenase component subunit alpha (TphA2) | BAE47085.1 | *Comamonas sp. E6* | [3] |
| | WP_047219860.1 | *Delftia tsuruhatensis sp.* | [4] |
| | WP_054022747.1 | *Ideonella sakaiensis* 201-F6 | [1] |
| | ABE33243.1 | *Paraburkholderia xenovorans LB400* | [5] |
| | WP_011599110.1 | *Rhodococcus jostii RHA1* | [6] |
| | AAR90187.1 | *Rhodococcus sp. DK17* | [7] |
| | AAX18941.1 | *Comamonas testosteroni* | [8] |
| | BAE47086.1 | *Comamonas sp. E6* | [3] |
| Terephthalate 1,2-dioxygenase, terminal oxygenase component subunit beta (TphA3) | WP_034369201.1 | *Delftia tsuruhatensis sp.* | [4] |
| | WP_054022748.1 | *Ideonella sakaiensis* 201-F6 | [1] |
| | ABE33244.1 | *Paraburkholderia xenovorans LB400* | [5] |
| | WP_005256013.1 | *Rhodococcus jostii RHA1* | [6] |
| | AAR90188.1 | *Rhodococcus sp. DK17* | [7] |
| | AAX18942.1 | *Comamonas testosteroni* | [8] |
| 1,2-dihydroxy-3,5-cyclohexadiene-1,4-dicarboxylate dehydrogenase (TphB) | BAE47087.1 | *Comamonas sp. E6* | [3] |
| | WP_047219859.1 | *Delftia tsuruhatensis sp.* | [4] |
| | WP_082368716.1 | *Ideonella sakaiensis* 201-F6 | [1] |
| | ABE33245.1 | *Paraburkholderia xenovorans LB400* | [5] |
| | WP_011599111.1 | *Rhodococcus jostii RHA1* | [6] |

| Name of TPA degrading enzyme* | NCBI entry | Organism | Reference |
|---|---|---|---|
| | AAR90189.1 | *Rhodococcus sp. DK17* | [7] |
| | AAX18943.1 | *Comamonas testosteroni* | [8] |

*The enzymes named TphA are part of the terephthalate 1,2-dioxygenase complex.

**Supplementary Table S6. Multifactorial ANOVA with Tukey's a posteriori test for Fig. 3.**

| Motifs | N | Average | Groups | | | |
|---|---|---|---|---|---|---|
| M0 | 109 | 19,4841 | a | | | |
| M4 strict | 108 | 5,6846 | | b | | |
| M4 | 109 | 4,2613 | | b | c | |
| M5 | 104 | 3,1902 | | | c | d |
| M3 strict | 108 | 2,1245 | | | c | d |
| M3 | 102 | 0,9050 | | | | d |

| Zone | N | Average | Groups | | | |
|---|---|---|---|---|---|---|
| Antarctic Deep | 54 | 11,0805 | a | | | |
| Antarctic Surface | 127 | 8,1637 | | b | | |
| Arctic Surface | 88 | 4,4565 | | | c | |
| Arctic Deep | 134 | 3,8040 | | | c | |
| Temperate | 237 | 2,2033 | | | | d |

| HMM Motifs and Zones | N | Average | Groups | | | | | | | | |
|---|---|---|---|---|---|---|---|---|---|---|---|
| M0 Antarctic Deep | 9 | 37,8810 | a | | | | | | | | |
| M0 Antarctic Surface | 22 | 20,7154 | | b | | | | | | | |
| M0 Arctic Deep | 23 | 16,9341 | | b | c | | | | | | |
| M0 Arctic Surface | 15 | 15,6330 | | b | c | d | | | | | |
| M5 Antarctic Deep | 9 | 10,8133 | | | c | d | e | f | g | | |
| M4 strict Antarctic Surface | 21 | 10,5625 | | | | d | e | | | | |
| M4 Antarctic Surface | 22 | 9,0805 | | | | | e | f | | | |
| M4 Antarctic Deep | 9 | 6,9907 | | | | | e | f | g | h | i |
| M4 strict Arctic Surface | 15 | 6,7062 | | | | | e | f | g | h | i |
| M0 Temperate | 40 | 6,2568 | | | | | e | f | g | h | i |
| M3 strict Antarctic Deep | 9 | 4,9232 | | | | | e | f | g | h | i |
| M4 strict Antarctic Deep | 9 | 4,5274 | | | | | e | f | g | h | i |

| | | | | | | |
|---|---|---|---|---|---|---|
| M5 Antarctic Surface | 20 | 4,2094 | f | g | h | i |
| M4 strict Temperate | 40 | 3,9195 | | g | h | i |
| M4 strict Arctic Deep | 23 | 2,7072 | | | h | i |
| M3 Antarctic Surface | 21 | 2,5436 | | | h | i |
| M4 Arctic Surface | 15 | 2,1872 | | | h | i |
| M3 strict Antarctic Surface | 21 | 1,8707 | | | h | i |
| M3 strict Arctic Surface | 15 | 1,7138 | | | h | i |
| M4 Temperate | 40 | 1,5752 | | | | i |
| M4 Arctic Deep | 23 | 1,4727 | | | h | i |
| M3 Antarctic Deep | 9 | 1,3474 | | | h | i |
| M3 strict Arctic Deep | 23 | 1,1171 | | | | i |
| M3 strict Temperate | 40 | 0,9978 | | | | i |
| M5 Arctic Deep | 23 | 0,4399 | | | | i |
| M5 Arctic Surface | 15 | 0,2835 | | | | i |
| M3 Temperate | 40 | 0,2653 | | | | |
| M3 Arctic Surface | 13 | 0,2153 | | | h | i |
| M5 Temperate | 37 | 0,2050 | | | | |
| M3 Arctic Deep | 19 | 0,1533 | | | | i |

**Supplementary Table S7. Multifactorial ANOVA with Tukey's a posteriori test for Supplementary Fig. S4.**

| Enzyme | N | Average | Groups | | |
|---|---|---|---|---|---|
| PETase | 131 | 35,4228 | a | | |
| TphA1 | 109 | 7,1139 | | b | |
| MHETase | 109 | 6,7573 | | b | |
| TphA2 | 103 | 1,6663 | | | c |
| TphA3 | 83 | 0,6416 | | | c |
| TphB | 92 | 0,4527 | | | c |

| Zone | N | Average | Groups | | |
|---|---|---|---|---|---|
| Antarctic Deep | 53 | 13,4788 | a | | |
| Antarctic Surface | 132 | 10,2421 | a | b | |
| Arctic Deep | 138 | 8,3471 | | b | |
| Arctic Surface | 90 | 7,7918 | | b | |
| Temperate | 214 | 3,5190 | | | c |

| Enzyme and Zone | N | Average | Groups | | | | | | | |
|---|---|---|---|---|---|---|---|---|---|---|
| PETase Antarctic Deep | 9 | 66,4831 | a | | | | | | | |
| PETase Antarctic Surface | 44 | 47,9187 | | b | | | | | | |
| PETase Arctic Surface | 15 | 26,7102 | | | c | | | | | |
| PETase Arctic Deep | 23 | 22,7976 | | | c | | | | | |
| PETase Temperate | 40 | 13,2043 | | | | d | | | | |
| MHETase Arctic Deep | 23 | 11,8516 | | | | d | e | | | |
| TphA1 Arctic Surface | 15 | 10,1846 | | | | d | e | f | | |
| TphA1 Arctic Deep | 23 | 8,9962 | | | | d | e | f | | |
| MHETase Antarctic Deep | 9 | 8,8076 | | | | d | e | f | g | h |
| TphA1 Antarctic Surface | 22 | 8,6629 | | | | d | e | f | | h |
| MHETase Arctic Surface | 15 | 6,9139 | | | | d | e | f | g | h |
| TphA1 Antarctic Deep | 9 | 4,1657 | | | | d | e | f | g | h |

| | | | | | | |
|---|---|---|---|---|---|---|
| TphA2 Arctic Deep | 23 | 4,1268 | e | f | g | h |
| MHETase Temperate | 40 | 3,7289 | | f | g | h |
| TphA1 Temperate | 40 | 3,5600 | | f | g | h |
| MHETase Antarctic Surface | 22 | 2,4844 | | f | g | h |
| TphA2 Arctic Surface | 15 | 2,0224 | e | f | g | h |
| TphA3 Arctic Deep | 23 | 1,2892 | | f | g | h |
| TphB Arctic Deep | 23 | 1,0210 | | f | g | h |
| TphA2 Antarctic Surface | 17 | 0,9607 | | f | g | h |
| TphB Antarctic Surface | 13 | 0,7187 | | f | g | h |
| TphA3 Antarctic Surface | 14 | 0,7073 | | f | g | h |
| TphA2 Antarctic Deep | 9 | 0,6746 | e | f | g | h |
| TphA3 Antarctic Deep | 8 | 0,6356 | e | f | g | h |
| TphA2 Temperate | 39 | 0,5468 | | | g | |
| TphA3 Arctic Surface | 15 | 0,5186 | | f | g | h |
| TphB Arctic Surface | 15 | 0,4008 | | f | g | h |
| TphB Antarctic Deep | 9 | 0,1059 | | f | g | h |
| TphA3 Temperate | 23 | 0,0573 | | | g | h |
| TphB Temperate | 32 | 0,0169 | | | g | |

**Supplementary Table S8. Composition of the reconstructed 112 MAGs.** Com = Completeness. Con = Contamination.

| MAGs ID | origin | classification | PETase | MHETase | TphA1 | TphA2 | TphA3 | TphB | PET seq ID | MHET seq ID | TphA1 seq ID | TphA2 seq ID | TphA3 seq ID | TphB seq ID | Com | Con | Size |
|---|---|---|---|---|---|---|---|---|---|---|---|---|---|---|---|---|---|
| TARA_ANE_MAG_00022 | TARA_Delmont_v1 | Bacteria; Proteobacteria; Gammaproteobacteria; Pseudomonadales; Pseudohongiellaceae; UBA9145; UBA9145 sp002457215 | 1 | 1 | FALSE | FALSE | FALSE | FALSE | TARA_ANE_MAG_00022_000000000327_1 | TARA_ANE_MAG_00022_000000000226_9 | FALSE | FALSE | FALSE | FALSE | 72.21 | 1.50 | 2656647 |
| TARA_ANE_MAG_00036 | TARA_Delmont_v3 | Bacteria; Bacteroidota; Bacteroidia; Flavobacteriales; BACL11; ; | 1 | 0 | FALSE | FALSE | FALSE | FALSE | TARA_ANE_MAG_00036_000000000077_5 | FALSE | FALSE | FALSE | FALSE | FALSE | 77.78 | 1.08 | 1200065 |
| TARA_ANE_MAG_00037 | TARA_Delmont_v4 | Bacteria; Proteobacteria; Gammaproteobacteria; Pseudomonadales; Saccharospirillaceae_A; Oleibacter; Oleibacter sp002331105 | 4 | 0 | FALSE | FALSE | FALSE | FALSE | TARA_ANE_MAG_00037_000000000033_15;TARA_ANE_MAG_00037_000000000033_16;TARA_ANE_MAG_00037_000000000033_17;TARA_ANE_MAG_00037_000000000054_5 | FALSE | FALSE | FALSE | FALSE | FALSE | 93.58 | 2.42 | 3266448 |
| TARA_ANE_MAG_00038 | TARA_Delmont_v5 | Bacteria; Verrucomicrobiota; Verrucomicrobiae; Verrucomicrobiales; Akkermansiaceae; Roseibacillus; Roseibacillus sp002420305 | 1 | 0 | FALSE | FALSE | FALSE | FALSE | TARA_ANE_MAG_00038_000000000088_4 | FALSE | FALSE | FALSE | FALSE | FALSE | 85.36 | 3.20 | 3563869 |
| TARA_ANE_MAG_00040 | TARA_Delmont_v6 | Bacteria; Bacteroidota; Bacteroidia; Flavobacteriales; BACL11; GCA-2693335; GCA-2693335 sp002724895 | 1 | 0 | FALSE | FALSE | FALSE | FALSE | TARA_ANE_MAG_00040_000000000009_19 | FALSE | FALSE | FALSE | FALSE | FALSE | 92.83 | 1.90 | 1640092 |
| TARA_ANE_MAG_00041 | TARA_Delmont_v7 | Bacteria; Bacteroidota; Bacteroidia; Flavobacteriales; BACL11; GCA-2715565; GCA-2715565 sp002715565 | 1 | 0 | FALSE | FALSE | FALSE | FALSE | TARA_ANE_MAG_00041_000000000148_6 | FALSE | FALSE | FALSE | FALSE | FALSE | 86.83 | 4.21 | 1515503 |

| | | | | | | | | | | | | | | | | | |
|---|---|---|---|---|---|---|---|---|---|---|---|---|---|---|---|---|---|
| TARA_ANE_MAG_00043 | TARA_Delmont_v8 | Bacteria; Bacteroidota; Bacteroidia; Flavobacteriales; BACL11; BACL11; | 1 | 0 | FALSE | FALSE | FALSE | FALSE | TARA_ANE_MAG_00043_000000000099_9 | FALSE | FALSE | FALSE | FALSE | FALSE | 84.61 | 2.96 | 1590729 |
| TARA_ANE_MAG_00061 | TARA_Delmont_v9 | Bacteria; Marinisomatota; Marinisomatia; SCGC-AAA003-L08; GCA-002707645; GCA-002718585; | 1 | 0 | FALSE | FALSE | FALSE | FALSE | TARA_ANE_MAG_00061_000000000124_4 | FALSE | FALSE | FALSE | FALSE | FALSE | 71.83 | 0.00 | 967588 |
| TARA_ANW_MAG_00004 | TARA_Delmont_v11 | Bacteria; Proteobacteria; Alphaproteobacteria; Sphingomonadales; Emcibacteraceae; Emcibacter; Emcibacter sp002380195 | 0 | 1 | FALSE | FALSE | FALSE | FALSE | FALSE | TARA_ANW_MAG_00004_000000000013_18 | FALSE | FALSE | FALSE | FALSE | 99.50 | 0.87 | 2825991 |
| TARA_ANW_MAG_00006 | TARA_Delmont_v12 | Bacteria; Proteobacteria; Gammaproteobacteria; Pseudomonadales; Ketobacteraceae; Ketobacter; Ketobacter sp002471665 | 3 | 0 | FALSE | FALSE | FALSE | FALSE | TARA_ANW_MAG_00006_000000000001_4;TARA_ANW_MAG_00006_000000000012_83;TARA_ANW_MAG_00006_000000000102_59 | FALSE | FALSE | FALSE | FALSE | FALSE | 98.70 | 4.43 | 5485499 |
| TARA_ANW_MAG_00009 | TARA_Delmont_v13 | Bacteria; Proteobacteria; Gammaproteobacteria; Pseudomonadales; Halieaceae; Halioglobus; Halioglobus sp002699145 | 0 | 1 | FALSE | FALSE | FALSE | FALSE | FALSE | TARA_ANW_MAG_00009_000000000065_239 | FALSE | FALSE | FALSE | FALSE | 98.97 | 1.59 | 5050254 |
| TARA_ANW_MAG_00030 | TARA_Delmont_v15 | Bacteria; Marinisomatota; Marinisomatia; SCGC-AAA003-L08; TMED6; ; | 1 | 0 | FALSE | FALSE | FALSE | FALSE | TARA_ANW_MAG_00030_000000000062_9 | FALSE | FALSE | FALSE | FALSE | FALSE | 90.01 | 2.00 | 1869178 |
| TARA_ASE_MAG_00006 | TARA_Delmont_v17 | Bacteria; Proteobacteria; Gammaproteobacteria; Pseudomonadales; Pseudomonadaceae; Pseudomonas_D; Pseudomonas_D aestusnigri | 1 | 0 | FALSE | FALSE | FALSE | FALSE | TARA_ASE_MAG_00006_000000000445_8 | FALSE | FALSE | FALSE | FALSE | FALSE | 76.46 | 1.16 | 2981946 |
| TARA_ASW_MAG_00004 | TARA_Delmont_v20 | Bacteria; Proteobacteria; Gammaproteobacteria; | 3 | 0 | FALSE | FALSE | FALSE | FALSE | TARA_ASW_MAG_00004_000000000138_ | FALSE | FALSE | FALSE | FALSE | FALSE | 92.72 | 1.91 | 3202824 |

| | | | | | | | | | | | | | | | | | |
|---|---|---|---|---|---|---|---|---|---|---|---|---|---|---|---|---|---|
| | | Pseudomonadales; Saccharospirillaceae_A; Oleibacter; Oleibacter sp002331105 | | | | | | | 2;TARA_ASW_MAG_00004_000000000138_3;TARA_ASW_MAG_00004_000000000138_4 | | | | | | | | |
| TARA_ASW_MAG_00012 | TARA_Delmont_v21 | Bacteria; Bacteroidota; Bacteroidia; Flavobacteriales; Flavobacteriaceae; Maribacter; Maribacter dokdonensis | 1 | 0 | FALSE | FALSE | FALSE | FALSE | TARA_ASW_MAG_00012_000000000042_26 | FALSE | FALSE | FALSE | FALSE | FALSE | 99.01 | 1.60 | 4451478 |
| TARA_ASW_MAG_00018 | TARA_Delmont_v22 | Bacteria; Bacteroidota; Bacteroidia; Flavobacteriales; Crocinitomicaceae; UBA952; UBA952 sp002480215 | 1 | 0 | FALSE | FALSE | FALSE | FALSE | TARA_ASW_MAG_00018_000000000097_2 | FALSE | FALSE | FALSE | FALSE | FALSE | 88.56 | 0.81 | 2217978 |
| TARA_ASW_MAG_00029 | TARA_Delmont_v23 | Bacteria; Proteobacteria; Gammaproteobacteria; Pseudomonadales; Pseudomonadaceae; Pseudomonas_D; Pseudomonas_D pachastrellae | 2 | 0 | FALSE | FALSE | FALSE | FALSE | TARA_ASW_MAG_00029_000000000024_47;TARA_ASW_MAG_00029_000000000037_8 | FALSE | FALSE | FALSE | FALSE | FALSE | 77.76 | 0.00 | 3864562 |
| TARA_ION_MAG_00067 | TARA_Delmont_v25 | Bacteria; Proteobacteria; Gammaproteobacteria; Pseudomonadales; Saccharospirillaceae_A; Oleibacter; Oleibacter sp002724925 | 3 | 0 | FALSE | FALSE | FALSE | FALSE | TARA_ION_MAG_00067_000000000050_3;TARA_ION_MAG_00067_000000000050_4;TARA_ION_MAG_00067_000000000050_5 | FALSE | FALSE | FALSE | FALSE | FALSE | 68.45 | 0.86 | 2941447 |
| TARA_IOS_MAG_00016 | TARA_Delmont_v26 | Bacteria; Marinisomatota; Marinisomatia; SCGC-AAA003-L08; GCA-2689995; GCA-2689995; | 1 | 0 | FALSE | FALSE | FALSE | FALSE | TARA_IOS_MAG_00016_000000000086_2 | FALSE | FALSE | FALSE | FALSE | FALSE | 92.11 | 0.53 | 1769221 |
| TARA_IOS_MAG_00026 | TARA_Delmont_v27 | Bacteria; Proteobacteria; Gammaproteobacteria; Pseudomonadales; Pseudohongiellaceae; UBA9145; | 1 | 2 | FALSE | FALSE | FALSE | FALSE | TARA_IOS_MAG_00026_000000000130_1 | TARA_IOS_MAG_00026_000000000225_2;TARA_IOS_MAG_00026_000000000342_4 | FALSE | FALSE | FALSE | FALSE | 58.97 | 2.23 | 1599775 |
| TARA_IOS_MAG_00041 | TARA_Delmont_v28 | Bacteria; Verrucomicrobiota; Verrucomicrobiae; | 1 | 0 | FALSE | FALSE | FALSE | FALSE | TARA_IOS_MAG_00041_00 | FALSE | FALSE | FALSE | FALSE | FALSE | 76.16 | 2.04 | 3169777 |

| | | | | | | | | | | | | | | | | | |
|---|---|---|---|---|---|---|---|---|---|---|---|---|---|---|---|---|---|
| | | Verrucomicrobiales; Akkermansiaceae; Roseibacillus; Roseibacillus sp002686085 | | | | | | | 0000000505_10 | | | | | | | | |
| TARA_MED_MAG_00002 | TARA_Delmont_v30 | Bacteria; Proteobacteria; Gammaproteobacteria; UBA6522; UBA6522; TMED78; TMED78 sp002171615 | 0 | 1 | FALSE | FALSE | FALSE | FALSE | FALSE | TARA_MED_MAG_00002_000000000023_16 | FALSE | FALSE | FALSE | FALSE | 79.71 | 0.21 | 1431420 |
| TARA_MED_MAG_00027 | TARA_Delmont_v33 | Bacteria; Marinisomatota; Marinisomatia; SCGC-AAA003-L08; D37C17; D37C17; | 1 | 0 | FALSE | FALSE | FALSE | FALSE | TARA_MED_MAG_00027_000000000069_2 | FALSE | FALSE | FALSE | FALSE | FALSE | 83.52 | 4.40 | 1825475 |
| TARA_MED_MAG_00058 | TARA_Delmont_v34 | Bacteria; Proteobacteria; Gammaproteobacteria; Pseudomonadales; Saccharospirillaceae_A; Oleibacter; Oleibacter sp002724925 | 4 | 0 | FALSE | FALSE | FALSE | FALSE | TARA_MED_MAG_00058_000000000072_6;TARA_MED_MAG_00058_000000000113_3;TARA_MED_MAG_00058_000000000113_4;TARA_MED_MAG_00058_000000000357_11 | FALSE | FALSE | FALSE | FALSE | FALSE | 83.88 | 2.02 | 3389560 |
| TARA_MED_MAG_00068 | TARA_Delmont_v35 | Bacteria; Proteobacteria; Gammaproteobacteria; Pseudomonadales; Pseudohongiellaceae; UBA9145; UBA9145 sp002477465 | 0 | 1 | FALSE | FALSE | FALSE | FALSE | FALSE | TARA_MED_MAG_00068_000000000097_3 | FALSE | FALSE | FALSE | FALSE | 74.94 | 1.06 | 2571986 |
| TARA_MED_MAG_00090 | TARA_Delmont_v36 | Bacteria; Bacteroidota; Bacteroidia; Flavobacteriales; Crocinitomicaceae; UBA952; UBA952 sp002167775 | 1 | 0 | FALSE | FALSE | FALSE | FALSE | TARA_MED_MAG_00090_000000000033_2 | FALSE | FALSE | FALSE | FALSE | FALSE | 87.24 | 2.16 | 2141649 |
| TARA_PON_MAG_00016 | TARA_Delmont_v37 | Bacteria; Bacteroidota; Bacteroidia; Flavobacteriales; BACL11; ; | 1 | 0 | FALSE | FALSE | FALSE | FALSE | TARA_PON_MAG_00016_000000000033_2 | FALSE | FALSE | FALSE | FALSE | FALSE | 87.72 | 0.00 | 1472662 |
| TARA_PON_MAG_00029 | TARA_Delmont_v38 | Bacteria; Verrucomicrobiota; Verrucomicrobiae; Verrucomicrobiales; Akkermansiaceae; | 1 | 0 | FALSE | FALSE | FALSE | FALSE | TARA_PON_MAG_00029_000000000005_2 | FALSE | FALSE | FALSE | FALSE | FALSE | 84.25 | 0.91 | 3024430 |

| | | | | | | | | | | | | | | | | | |
|---|---|---|---|---|---|---|---|---|---|---|---|---|---|---|---|---|---|
| | | SW10; SW10 sp002729295 | | | | | | | | | | | | | | | |
| TARA_PON_MAG_00036 | TARA_Delmont_v39 | Bacteria; Bacteroidota; Bacteroidia; Flavobacteriales; BACL11; GCA-2711125; | 1 | 0 | FALSE | FALSE | FALSE | FALSE | TARA_PON_MAG_00036_000000000125_3 | FALSE | FALSE | FALSE | FALSE | FALSE | 74.19 | 0.81 | 1543517 |
| TARA_PON_MAG_00066 | TARA_Delmont_v41 | Bacteria; Acidobacteriota; Vicinamibacteria; Vicinamibacterales; UBA8438; UBA8438; UBA8438 sp002726085 | 1 | 4 | FALSE | FALSE | FALSE | FALSE | TARA_PON_MAG_00066_000000000256_4 | TARA_PON_MAG_00066_000000000252_4;TARA_PON_MAG_00066_000000000385_3;TARA_PON_MAG_00066_000000000480_19;TARA_PON_MAG_00066_000000000541_2 | FALSE | FALSE | FALSE | FALSE | 51.78 | 0.51 | 3199981 |
| TARA_PSE_MAG_00024 | TARA_Delmont_v44 | Bacteria; Proteobacteria; Alphaproteobacteria; Rhodobacterales; Rhodobacteraceae; Salipiger; Salipiger profundus | 0 | 1 | FALSE | FALSE | FALSE | FALSE | FALSE | TARA_PSE_MAG_00024_000000000001_20 | FALSE | FALSE | FALSE | FALSE | 97.31 | 0.75 | 4253894 |
| TARA_PSE_MAG_00048 | TARA_Delmont_v45 | Bacteria; Verrucomicrobiota; Verrucomicrobiae; Pedosphaerales; UBA1100; UBA1100; UBA1100 sp003509725 | 1 | 0 | FALSE | FALSE | FALSE | FALSE | TARA_PSE_MAG_00048_000000000330_4 | FALSE | FALSE | FALSE | FALSE | FALSE | 74.01 | 1.27 | 2529742 |
| TARA_PSE_MAG_00057 | TARA_Delmont_v46 | Bacteria; Marinisomatota; Marinisomatia; SCGC-AAA003-L08; GCA-002707645; GCA-002718585; | 1 | 0 | FALSE | FALSE | FALSE | FALSE | TARA_PSE_MAG_00057_000000000061_5 | FALSE | FALSE | FALSE | FALSE | FALSE | 71.98 | 2.75 | 1223860 |
| TARA_PSE_MAG_00120 | TARA_Delmont_v49 | Bacteria; Proteobacteria; Gammaproteobacteria; Pseudomonadales; Pseudomonadaceae; Pseudomonas_D; Pseudomonas_D sp002683975 | 2 | 0 | FALSE | FALSE | FALSE | FALSE | TARA_PSE_MAG_00120_000000000041_3;TARA_PSE_MAG_00120_000000000340_2 | | FALSE | FALSE | FALSE | FALSE | 75.00 | 0.00 | 2768210 |
| TARA_PSE_MAG_00130 | TARA_Delmont_v50 | Bacteria; Proteobacteria; Gammaproteobacteria; Pseudomonadales; Saccharospirillaceae_A; | 1 | 0 | FALSE | FALSE | FALSE | FALSE | TARA_PSE_MAG_00130_000000000313_1 | FALSE | FALSE | FALSE | FALSE | FALSE | 63.79 | 0.00 | 2657260 |

| | | | | | | | | | | | | | | | | | |
|---|---|---|---|---|---|---|---|---|---|---|---|---|---|---|---|---|---|
| | | Bermanella; Bermanella sp002683575 | | | | | | | | | | | | | | | |
| TARA_PSE_MAG_00133 | TARA_Delmont_v51 | Bacteria; Proteobacteria; Gammaproteobacteria; Pseudomonadales; Saccharospirillaceae_A; Thalassolituus; Thalassolituus oleivorans | 1 | 0 | FALSE | FALSE | FALSE | FALSE | TARA_PSE_MAG_00133_000000000310_4 | FALSE | FALSE | FALSE | FALSE | FALSE | 51.03 | 0.00 | 2637354 |
| TARA_PSW_MAG_00040 | TARA_Delmont_v54 | Bacteria; Proteobacteria; Alphaproteobacteria; Defluviicoccales; Defluviicoccaceae; UBA830; UBA830 sp002471575 | 0 | 0 | FALSE | TRUE | FALSE | FALSE | FALSE | FALSE | FALSE | TARA_PSW_MAG_00040_000000000006_3 | FALSE | FALSE | 96.42 | 1.99 | 3100748 |
| TARA_PSW_MAG_00047 | TARA_Delmont_v55 | Bacteria; Bacteroidota; Bacteroidia; Flavobacteriales; Crocinitomicaceae; UBA952; | 1 | 0 | FALSE | FALSE | FALSE | FALSE | TARA_PSW_MAG_00047_000000000026_3 | FALSE | FALSE | FALSE | FALSE | FALSE | 90.05 | 1.35 | 2170786 |
| TARA_PSW_MAG_00060 | TARA_Delmont_v56 | Bacteria; Proteobacteria; Gammaproteobacteria; Pseudomonadales; Pseudohongiellaceae; UBA9145; UBA9145 sp002719575 | 1 | 0 | FALSE | FALSE | FALSE | FALSE | TARA_PSW_MAG_00060_000000000118_2 | FALSE | FALSE | FALSE | FALSE | FALSE | 82.68 | 2.00 | 2442486 |
| TARA_PSW_MAG_00062 | TARA_Delmont_v57 | Bacteria; Proteobacteria; Gammaproteobacteria; Pseudomonadales; Pseudomonadaceae; Pseudomonas_D; Pseudomonas_D pachastrellae | 2 | 0 | FALSE | FALSE | FALSE | FALSE | TARA_PSW_MAG_00062_000000000118_3;TARA_PSW_MAG_00062_000000000548_6 | FALSE | FALSE | FALSE | FALSE | FALSE | 79.61 | 1.99 | 3467416 |
| TARA_PSW_MAG_00064 | TARA_Delmont_v58 | Bacteria; Myxococcota; UBA727; UBA4151; UBA4151; UBA4151; UBA4151 sp002696345 | 1 | 0 | FALSE | FALSE | FALSE | FALSE | TARA_PSW_MAG_00064_000000000356_3 | FALSE | FALSE | FALSE | FALSE | FALSE | 74.10 | 1.97 | 3020418 |
| TARA_PSW_MAG_00070 | TARA_Delmont_v59 | Bacteria; Verrucomicrobiota; Verrucomicrobiae; Verrucomicrobiales; Akkermansiaceae; Roseibacillus; Roseibacillus sp002715825 | 1 | 0 | FALSE | FALSE | FALSE | FALSE | TARA_PSW_MAG_00070_000000000272_6 | FALSE | FALSE | FALSE | FALSE | FALSE | 69.05 | 0.68 | 2545416 |

| | | | | | | | | | | | | | | | | | |
|---|---|---|---|---|---|---|---|---|---|---|---|---|---|---|---|---|---|
| TARA_PSW_MAG_00080 | TARA_Delmont_v60 | Bacteria; Verrucomicrobiota; Verrucomicrobiae; Verrucomicrobiales; Akkermansiaceae; Roseibacillus; Roseibacillus sp002709915 | 1 | 0 | FALSE | FALSE | FALSE | FALSE | TARA_PSW_MAG_00080_000000000270_8 | FALSE | FALSE | FALSE | FALSE | FALSE | 68.94 | 0.68 | 2324340 |
| TARA_PSW_MAG_00088 | TARA_Delmont_v61 | Bacteria; Marinisomatota; Marinisomatia; SCGC-AAA003-L08; TMED6; ; | 1 | 0 | FALSE | FALSE | FALSE | FALSE | TARA_PSW_MAG_00088_000000000018_4 | FALSE | FALSE | FALSE | FALSE | FALSE | 55.29 | 0.47 | 1188083 |
| TARA_RED_MAG_00006 | TARA_Delmont_v63 | Bacteria; Proteobacteria; Alphaproteobacteria; UBA8366; UBA8366; UBA6544; UBA6544 sp002724555 | 0 | 0 | FALSE | TRUE | FALSE | FALSE | FALSE | FALSE | FALSE | TARA_RED_MAG_00006_000000000085_34 | FALSE | FALSE | 96.13 | 1.96 | 3754020 |
| TARA_RED_MAG_00008 | TARA_Delmont_v64 | Bacteria; Acidobacteriota; Vicinamibacteria; Vicinamibacterales; UBA8438; ; | 0 | 1 | FALSE | FALSE | FALSE | FALSE | FALSE | TARA_RED_MAG_00008_000000000060_126 | FALSE | FALSE | FALSE | FALSE | 94.82 | 6.03 | 3017963 |
| TARA_RED_MAG_00022 | TARA_Delmont_v65 | Bacteria; Proteobacteria; Gammaproteobacteria; Pseudomonadales; Saccharospirillaceae_A; Oleibacter; Oleibacter sp002724925 | 4 | 0 | FALSE | FALSE | FALSE | FALSE | TARA_RED_MAG_00022_000000000003_9;TARA_RED_MAG_00022_000000000003_10;TARA_RED_MAG_00022_000000000003_11;TARA_RED_MAG_00022_000000000011_30 | FALSE | FALSE | FALSE | FALSE | FALSE | 97.64 | 0.89 | 3655333 |
| TARA_RED_MAG_00030 | TARA_Delmont_v66 | Bacteria; Acidobacteriota; Vicinamibacteria; Vicinamibacterales; UBA8438; UBA8438; | 1 | 1 | FALSE | FALSE | FALSE | FALSE | TARA_RED_MAG_00030_000000000136_2 | TARA_RED_MAG_00030_000000000054_11 | FALSE | FALSE | FALSE | FALSE | 88.89 | 3.69 | 4023354 |
| TARA_RED_MAG_00041 | TARA_Delmont_v67 | Bacteria; Chloroflexota; UBA11872; UBA11872; UBA11872; UBA11872; UBA11872 sp002433065 | 0 | 1 | FALSE | FALSE | FALSE | FALSE | FALSE | TARA_RED_MAG_00041_000000000028_21 | FALSE | FALSE | FALSE | FALSE | 83.95 | 0.00 | 2113769 |
| TARA_RED_MAG_00046 | TARA_Delmont_v68 | Bacteria; Bacteroidota; Bacteroidia; Flavobacteriales; BACL11; GCA-2711125; | 1 | 0 | FALSE | FALSE | FALSE | FALSE | TARA_RED_MAG_00046_000000000029_7 | FALSE | FALSE | FALSE | FALSE | FALSE | 89.42 | 1.61 | 1522052 |

| | | | | | | | | | | | | | | | | | |
|---|---|---|---|---|---|---|---|---|---|---|---|---|---|---|---|---|---|
| TARA_RED_MAG_00081 | TARA_Delmont_v70 | Bacteria; Acidobacteriota; Vicinamibacteria; Vicinamibacterales; UBA8438; UBA8438; | 1 | 0 | FALSE | FALSE | FALSE | FALSE | TARA_RED_MAG_00081_000000000175_2 | FALSE | FALSE | FALSE | FALSE | FALSE | 80.74 | 1.92 | 4159930 |
| TARA_RED_MAG_00093 | TARA_Delmont_v71 | Bacteria; Planctomycetota; Planctomycetes; Pirellulales; Pirellulaceae; UBA9264; UBA9264 sp002724245 | 1 | 0 | FALSE | FALSE | FALSE | FALSE | TARA_RED_MAG_00093_000000000211_6 | FALSE | FALSE | FALSE | FALSE | FALSE | 91.49 | 1.18 | 6139921 |
| TARA_SOC_MAG_00002 | TARA_Delmont_v72 | Bacteria; Proteobacteria; Gammaproteobacteria; Enterobacterales; Alteromonadaceae; Alteromonas_A; Alteromonas_A sp002335925 | 0 | 1 | FALSE | FALSE | FALSE | FALSE | FALSE | TARA_SOC_MAG_00002_000000000072_23 | FALSE | FALSE | FALSE | FALSE | 97.15 | 1.26 | 4429622 |
| TARA_SOC_MAG_00003 | TARA_Delmont_v73 | Bacteria; Bacteroidota; Bacteroidia; Flavobacteriales; UA16; UBA974; UBA974 sp002339135 | 1 | 0 | FALSE | FALSE | FALSE | FALSE | TARA_SOC_MAG_00003_000000000045_1 | FALSE | FALSE | FALSE | FALSE | FALSE | 92.83 | 0.02 | 1857767 |
| TARA_SOC_MAG_00004 | TARA_Delmont_v74 | Bacteria; Bacteroidota; Bacteroidia; Flavobacteriales; Salibacteraceae; UBA6770; UBA6770 sp002453255 | 1 | 0 | FALSE | FALSE | FALSE | FALSE | TARA_SOC_MAG_00004_000000000078_1 | FALSE | FALSE | FALSE | FALSE | FALSE | 93.28 | 0.11 | 2708830 |
| TARA_SOC_MAG_00007 | TARA_Delmont_v75 | Bacteria; Proteobacteria; Gammaproteobacteria; Pseudomonadales; Pseudohongiellaceae; ; | 2 | 1 | FALSE | FALSE | FALSE | FALSE | TARA_SOC_MAG_00007_000000000017_2;TARA_SOC_MAG_00007_000000000193_2 | TARA_SOC_MAG_00007_000000000068_3 | FALSE | FALSE | FALSE | FALSE | 78.13 | 1.73 | 2918716 |
| TARA_SOC_MAG_00011 | TARA_Delmont_v76 | Bacteria; Bacteroidota; Bacteroidia; Flavobacteriales; Flavobacteriaceae; Polaribacter; Polaribacter sp000981625 | 2 | 0 | FALSE | FALSE | FALSE | FALSE | TARA_SOC_MAG_00011_000000000041_18;TARA_SOC_MAG_00011_000000000111_15 | FALSE | FALSE | FALSE | FALSE | FALSE | 84.20 | 0.64 | 2520155 |
| Genome_008 | PM-RGC | Bacteria; Proteobacteria; Gammaproteobacteria; UBA10353; UBA7415; Arenicella; | 0 | 1 | FALSE | FALSE | FALSE | FALSE | FALSE | Genome_008_k141_513483_2 | FALSE | FALSE | FALSE | FALSE | 81.95 | 2.00 | 2695447 |

| | | | | | | | | | | | | | | | | | |
|---|---|---|---|---|---|---|---|---|---|---|---|---|---|---|---|---|---|
| Genome_014 | PM-RGC | Bacteria; Proteobacteria; Alphaproteobacteria; Micavibrionales; Micavibrionaceae; ; | 1 | 0 | FALSE | FALSE | FALSE | FALSE | Genome_014_k141_334157_53 | FALSE | FALSE | FALSE | FALSE | FALSE | 94.14 | 0.43 | 2202676 |
| Genome_026 | PM-RGC | Bacteria; Proteobacteria; Alphaproteobacteria; Rhodospirillales; UBA2165; GCA-2712665; GCA-2712665 sp002938035 | 1 | 0 | FALSE | FALSE | FALSE | FALSE | Genome_026_k141_469205_1 | FALSE | FALSE | FALSE | FALSE | FALSE | 91.17 | 4.66 | 3290634 |
| Genome_040 | PM-RGC | Bacteria; Proteobacteria; Gammaproteobacteria; Burkholderiales; Burkholderiaceae; RS62; RS62 sp000496475 | 1 | 0 | FALSE | FALSE | FALSE | FALSE | Genome_040_k141_193472_3 | FALSE | FALSE | FALSE | FALSE | FALSE | 97.29 | 1.55 | 2529036 |
| Genome_055 | PM-RGC | Bacteria; Proteobacteria; Gammaproteobacteria; Pseudomonadales; Pseudohongiellaceae; UBA9145; UBA9145 sp002731775 | 1 | 1 | FALSE | FALSE | FALSE | FALSE | Genome_055_k141_78115_114 | Genome_055_k141_135728_16 | FALSE | FALSE | FALSE | FALSE | 92.96 | 0.37 | 3566335 |
| Genome_069 | PM-RGC | Bacteria; Proteobacteria; Gammaproteobacteria; Enterobacterales; Alteromonadaceae; Pseudoalteromonas; Pseudoalteromonas marina | 1 | 0 | FALSE | FALSE | FALSE | FALSE | Genome_069_k141_13446_4 | FALSE | FALSE | FALSE | FALSE | FALSE | 96.52 | 3.33 | 4118936 |
| Genome_076 | PM-RGC | Bacteria; Proteobacteria; Gammaproteobacteria; Pseudomonadales; Pseudohongiellaceae; UBA5109; UBA5109 sp002414385 | 0 | 1 | FALSE | FALSE | FALSE | FALSE | FALSE | Genome_076_k141_518704_14 | FALSE | FALSE | FALSE | FALSE | 92.84 | 1.39 | 2493334 |
| Genome_077 | PM-RGC | Bacteria; Proteobacteria; Gammaproteobacteria; Pseudomonadales; Pseudohongiellaceae; Pseudohongiella; | 2 | 1 | FALSE | FALSE | FALSE | FALSE | Genome_077_k141_830733_542;Genome_077_k141_633034_649 | Genome_077_k141_830733_29 | FALSE | FALSE | FALSE | FALSE | 96.67 | 1.30 | 3653317 |
| Genome_078 | PM-RGC | Bacteria; Proteobacteria; Gammaproteobacteria; | 0 | 2 | FALSE | FALSE | FALSE | FALSE | FALSE | Genome_078_k141_816444_83;Genome_0 | FALSE | FALSE | FALSE | FALSE | 94.73 | 1.67 | 2660278 |

| | | | | | | | | | | | | | | | | | |
|---|---|---|---|---|---|---|---|---|---|---|---|---|---|---|---|---|---|
| | | Pseudomonadales; Pseudohongiellaceae; UBA5109; | | | | | | | | 78_k141_591403_59 | | | | | | | |
| Genome_081 | PM-RGC | Bacteria; Proteobacteria; Gammaproteobacteria; Pseudomonadales; Porticoccaceae; BRH-c0; | 0 | 3 | FALSE | FALSE | FALSE | FALSE | FALSE | Genome_081_k141_475952_79;Genome_081_k141_403167_7;Genome_081_k141_403167_10 | FALSE | FALSE | FALSE | FALSE | 99.38 | 1.11 | 3257516 |
| Genome_085 | PM-RGC | Bacteria; Proteobacteria; Gammaproteobacteria; Pseudomonadales; Pseudomonadaceae; Pseudomonas_D; | 7 | 0 | FALSE | FALSE | FALSE | FALSE | Genome_085_k141_1008732_40;Genome_085_k141_1008732_41;Genome_085_k141_146371_1;Genome_085_k141_515476_12;Genome_085_k141_515476_13;Genome_085_k141_515476_14;Genome_085_k141_593680_5 | FALSE | FALSE | FALSE | FALSE | FALSE | 88.39 | 5.20 | 3063467 |
| Genome_086 | PM-RGC | Bacteria; Proteobacteria; Gammaproteobacteria; Pseudomonadales; Spongiibacteraceae; Zhongshania; Zhongshania aliphaticivorans | 0 | 1 | FALSE | FALSE | FALSE | FALSE | FALSE | Genome_086_k141_680256_69 | FALSE | FALSE | FALSE | FALSE | 97.84 | 1.94 | 3926890 |
| Genome_088 | PM-RGC | Bacteria; Proteobacteria; Gammaproteobacteria; Enterobacterales; Alteromonadaceae; Idiomarina; Idiomarina sp004025325 | 0 | 1 | FALSE | FALSE | FALSE | FALSE | FALSE | Genome_088_k141_319900_1 | FALSE | FALSE | FALSE | FALSE | 98.25 | 1.88 | 2369573 |
| Genome_092 | PM-RGC | Bacteria; Proteobacteria; Gammaproteobacteria; Nevskiales; ; ; | 1 | 0 | FALSE | FALSE | FALSE | FALSE | Genome_092_k141_477014_12 | FALSE | FALSE | FALSE | FALSE | FALSE | 96.82 | 0.82 | 2875534 |
| Genome_096 | PM-RGC | Bacteria; Proteobacteria; Gammaproteobacteria; Pseudomonadales; Pseudohongiellaceae; UBA9145; UBA9145 sp002731775 | 1 | 1 | FALSE | FALSE | FALSE | FALSE | Genome_096_k141_705644_76 | Genome_096_k141_792116_77 | FALSE | FALSE | FALSE | FALSE | 91.98 | 1.91 | 3437796 |

| | | | | | | | | | | | | | | | | | |
|---|---|---|---|---|---|---|---|---|---|---|---|---|---|---|---|---|---|
| Genome_099 | PM-RGC | Bacteria; Proteobacteria; Gammaproteobacteria; Pseudomonadales; Pseudohongiellaceae; UBA9145; | 3 | 2 | FALSE | FALSE | FALSE | FALSE | Genome_099_k141_507008_110;Genome_099_k141_580178_43;Genome_099_k141_408748_2 | Genome_099_k141_507008_8;Genome_099_k141_748089_17 | FALSE | FALSE | FALSE | FALSE | 92.50 | 1.23 | 3274457 |
| Genome_101 | PM-RGC | Bacteria; Proteobacteria; Gammaproteobacteria; Enterobacterales; Alteromonadaceae; Pseudoalteromonas; Pseudoalteromonas aliena | 1 | 0 | FALSE | FALSE | FALSE | FALSE | Genome_101_k141_444081_20 | FALSE | FALSE | FALSE | FALSE | FALSE | 97.52 | 2.61 | 4398842 |
| Genome_102 | PM-RGC | Bacteria; Proteobacteria; Gammaproteobacteria; Pseudomonadales; HTCC2089; UBA11889; UBA11889 sp002313255 | 1 | 0 | FALSE | FALSE | FALSE | FALSE | Genome_102_k141_782857_4 | FALSE | FALSE | FALSE | FALSE | FALSE | 86.02 | 2.45 | 2826849 |
| Genome_103 | PM-RGC | Bacteria; Proteobacteria; Gammaproteobacteria; Pseudomonadales; Pseudohongiellaceae; OM182; OM182 sp001438145 | 1 | 0 | FALSE | FALSE | FALSE | FALSE | Genome_103_k141_82266_5 | FALSE | FALSE | FALSE | FALSE | FALSE | 95.08 | 4.03 | 2561645 |
| Genome_105 | PM-RGC | Bacteria; Proteobacteria; Gammaproteobacteria; Pseudomonadales; Oleiphilaceae; PZPK01; | 1 | 0 | FALSE | FALSE | FALSE | FALSE | Genome_105_k141_1085022_39 | FALSE | FALSE | FALSE | FALSE | FALSE | 97.37 | 0.72 | 3466696 |
| Genome_106 | PM-RGC | Bacteria; Proteobacteria; Gammaproteobacteria; Pseudomonadales; Pseudohongiellaceae; UBA9145; | 2 | 1 | FALSE | FALSE | FALSE | FALSE | Genome_106_k141_572073_49;Genome_106_k141_895536_7 | Genome_106_k141_628886_10 | FALSE | FALSE | FALSE | FALSE | 84.36 | 2.04 | 3291670 |
| Genome_111 | PM-RGC | Bacteria; Proteobacteria; Gammaproteobacteria; Enterobacterales; Alteromonadaceae; Alteromonas_A; | 0 | 1 | FALSE | FALSE | FALSE | FALSE | FALSE | Genome_111_k141_466671_6 | FALSE | FALSE | FALSE | FALSE | 91.26 | 3.98 | 4584948 |
| Genome_114 | PM-RGC | Bacteria; Proteobacteria; Gammaproteobacteria; Enterobacterales; Alteromonadaceae; | 1 | 0 | FALSE | FALSE | FALSE | FALSE | Genome_114_k141_744438_17 | FALSE | FALSE | FALSE | FALSE | FALSE | 95.89 | 2.03 | 4053032 |

| | | | | | | | | | | | | | | | | | |
|---|---|---|---|---|---|---|---|---|---|---|---|---|---|---|---|---|---|
| | | Pseudoalteromonas; Pseudoalteromonas marina | | | | | | | | | | | | | | | |
| Genome_116 | PM-RGC | Bacteria; Proteobacteria; Gammaproteobacteria; Pseudomonadales; Pseudohongiellaceae; UBA9145; | 3 | 2 | FALSE | FALSE | FALSE | FALSE | Genome_116_k141_754012_45;Genome_116_k141_1542_44;Genome_116_k141_924706_17 | Genome_116_k141_912763_7;Genome_116_k141_307028_49 | FALSE | FALSE | FALSE | FALSE | 92.59 | 1.90 | 3358426 |
| Genome_122 | PM-RGC | Bacteria; Proteobacteria; Gammaproteobacteria; Immundisolibacterales; Immundisolibacteraceae; Immundisolibacter; | 0 | 0 | FALSE | TRUE | FALSE | FALSE | FALSE | FALSE | FALSE | Genome_122_k141_746842_47 | FALSE | FALSE | 96.95 | 1.22 | 3245373 |
| Genome_133 | PM-RGC | Bacteria; Bacteroidota; Bacteroidia; Flavobacteriales; Flavobacteriaceae; HC6-5; | 1 | 0 | FALSE | FALSE | FALSE | FALSE | Genome_133_k141_6248_21 | FALSE | FALSE | FALSE | FALSE | FALSE | 88.24 | 2.56 | 1857360 |
| Genome_136 | PM-RGC | Bacteria; Bacteroidota; Bacteroidia; Flavobacteriales; Flavobacteriaceae; HC6-5; HC6-5 sp003450155 | 1 | 0 | FALSE | FALSE | FALSE | FALSE | Genome_136_k141_123361_5 | FALSE | FALSE | FALSE | FALSE | FALSE | 93.23 | 1.27 | 2039446 |
| Genome_152 | PM-RGC | Bacteria; Bacteroidota; Bacteroidia; Flavobacteriales; Vicingaceae; ; | 1 | 0 | FALSE | FALSE | FALSE | FALSE | Genome_152_k141_529945_32 | FALSE | FALSE | FALSE | FALSE | FALSE | 99.46 | 0.65 | 3547026 |
| Genome_154 | PM-RGC | Bacteria; Bacteroidota; Bacteroidia; Flavobacteriales; Crocinitomicaceae; UBA4466; | 1 | 0 | FALSE | FALSE | FALSE | FALSE | Genome_154_k141_13852_65 | FALSE | FALSE | FALSE | FALSE | FALSE | 100.0 | 0.00 | 4063552 |
| Genome_160 | PM-RGC | Bacteria; Bacteroidota; Bacteroidia; Flavobacteriales; Crocinitomicaceae; UBA4466; | 1 | 0 | FALSE | FALSE | FALSE | FALSE | Genome_160_k141_358717_5 | FALSE | FALSE | FALSE | FALSE | FALSE | 97.30 | 3.69 | 2828968 |
| Genome_166 | PM-RGC | Bacteria; Bacteroidota; Bacteroidia; Flavobacteriales; Crocinitomicaceae; UBA4466; | 1 | 0 | FALSE | FALSE | FALSE | FALSE | Genome_166_k141_935554_26 | FALSE | FALSE | FALSE | FALSE | FALSE | 80.54 | 0.00 | 3480720 |
| Genome_178 | PM-RGC | Bacteria; Actinobacteriota; Acidimicrobiia; | 1 | 0 | FALSE | FALSE | FALSE | FALSE | Genome_178_k141_330579_40 | FALSE | FALSE | FALSE | FALSE | FALSE | 99.15 | 1.28 | 4570169 |

| | | | | | | | | | | | | | | | | | |
|---|---|---|---|---|---|---|---|---|---|---|---|---|---|---|---|---|---|
| | | Acidimicrobiales; UBA8139; ; | | | | | | | | | | | | | | | |
| Genome_204 | PM-RGC | Bacteria; Planctomycetota; Planctomycetes; Pirellulales; Pirellulaceae; Mariniblastus; | 1 | 0 | FALSE | FALSE | FALSE | FALSE | Genome_204_k141_417965_4 | FALSE | FALSE | FALSE | FALSE | FALSE | 85.41 | 3.47 | 4377633 |
| Genome_212 | PM-RGC | Bacteria; Verrucomicrobiota; Verrucomicrobiae; Verrucomicrobiales; Akkermansiaceae; ; | 1 | 0 | FALSE | FALSE | FALSE | FALSE | Genome_212_k141_663810_5 | FALSE | FALSE | FALSE | FALSE | FALSE | 89.93 | 2.35 | 4847897 |
| Genome_214 | PM-RGC | Bacteria; Verrucomicrobiota; Verrucomicrobiae; Verrucomicrobiales; Akkermansiaceae; SW10; SW10 sp000383735 | 1 | 0 | FALSE | FALSE | FALSE | FALSE | Genome_214_k141_101183_20 | FALSE | FALSE | FALSE | FALSE | FALSE | 99.29 | 0.00 | 4189413 |
| Ant_ICE_2016_bin.26 | Chile_Bay | Bacteria; Proteobacteria; Gammaproteobacteria; Burkholderiales; Burkholderiaceae; Rhodoferax; | 1 | 0 | FALSE | FALSE | FALSE | FALSE | NODE_2181_length_10778_cov_8.024900_5 | FALSE | FALSE | FALSE | FALSE | FALSE | 78.47 | 0.56 | 3445311 |
| Ant_ICE_2016_bin.3 | Chile_Bay | Bacteria; Proteobacteria; Alphaproteobacteria; Sphingomonadales; Sphingomonadaceae; Sphingomonas; | 1 | 1 | FALSE | FALSE | FALSE | FALSE | NODE_19222_length_2634_cov_2.400155_2 | NODE_63191_length_1194_cov_2.189640_1 | FALSE | FALSE | FALSE | FALSE | 50.55 | 6.01 | 2371488 |
| Ant_P3_2_2016_bin.13 | Chile_Bay | Bacteria; Bacteroidota; Bacteroidia; Flavobacteriales; BACL11; ; | 1 | 0 | FALSE | FALSE | FALSE | FALSE | NODE_19_length_93834_cov_8.209972_2 | FALSE | FALSE | FALSE | FALSE | FALSE | 95.43 | 7.26 | 1956410 |
| Ant_P3_2_2016_bin.7 | Chile_Bay | Bacteria; Bacteroidota; Bacteroidia; Flavobacteriales; Flavobacteriaceae; Polaribacter; | 1 | 0 | FALSE | FALSE | FALSE | FALSE | NODE_230_length_30031_cov_4.477449_16 | FALSE | FALSE | FALSE | FALSE | FALSE | 51.05 | 2.63 | 1253845 |
| Ant_P3_2_2018_bin.1 | Chile_Bay | Bacteria; Proteobacteria; Alphaproteobacteria; Rhodobacterales; Rhodobacteraceae; Yoonia; | 1 | 0 | FALSE | FALSE | FALSE | FALSE | NODE_1914_length_7160_cov_3.307952_7 | FALSE | FALSE | FALSE | FALSE | FALSE | 78.86 | 2.91 | 2377493 |
| Ant_P3_2_2018_bin.11 | Chile_Bay | Bacteria; Bacteroidota; Bacteroidia; | 1 | 0 | FALSE | FALSE | FALSE | FALSE | NODE_10_length_132181_c | FALSE | FALSE | FALSE | FALSE | FALSE | 97.12 | 0.11 | 2443073 |

| | | | | | | | | | | | | | | | | | |
|---|---|---|---|---|---|---|---|---|---|---|---|---|---|---|---|---|---|
| | | Flavobacteriales; Flavobacteriaceae; Polaribacter; | | | | | | | ov_16.874120_120 | | | | | | | | |
| Ant_P3_2_2018_bin.2 | Chile_Bay | Bacteria; Bacteroidota; Bacteroidia; Flavobacteriales; Flavobacteriaceae; HC6-5; HC6-5 sp003450155 | 1 | 0 | FALSE | FALSE | FALSE | FALSE | NODE_5578_length_3362_cov_5.721500_1 | FALSE | FALSE | FALSE | FALSE | FALSE | 87.87 | 1.57 | 1863293 |
| Ant_P3_2_2018_bin.4 | Chile_Bay | Bacteria; Proteobacteria; Alphaproteobacteria; Rhodobacterales; Rhodobacteraceae; Ascidiaceihabitans; | 1 | 0 | FALSE | FALSE | FALSE | FALSE | NODE_50_length_76774_cov_9.642279_35 | FALSE | FALSE | FALSE | FALSE | FALSE | 98.92 | 0.64 | 2906169 |
| Ant_P3_2_2019_bin.9 | Chile_Bay | Bacteria; Bacteroidota; Bacteroidia; Flavobacteriales; Flavobacteriaceae; HC6-5; HC6-5 sp003450155 | 1 | 0 | FALSE | FALSE | FALSE | FALSE | NODE_5500_length_4454_cov_8.765401_1 | FALSE | FALSE | FALSE | FALSE | FALSE | 80.17 | 0.00 | 1778277 |
| Ant_P3_30_2019_bin.16 | Chile_Bay | Bacteria; Bacteroidota; Bacteroidia; Flavobacteriales; Flavobacteriaceae; HC6-5; HC6-5 sp003450155 | 1 | 0 | FALSE | FALSE | FALSE | FALSE | NODE_389_length_32729_cov_10.819490_7 | FALSE | FALSE | FALSE | FALSE | FALSE | 92.72 | 0.00 | 1887449 |
| Ant_P3_30_2019_bin.22 | Chile_Bay | Bacteria; Bacteroidota; Bacteroidia; Flavobacteriales; Flavobacteriaceae; Polaribacter; | 1 | 0 | FALSE | FALSE | FALSE | FALSE | NODE_1743_length_11732_cov_34.379464_3 | FALSE | FALSE | FALSE | FALSE | FALSE | 87.81 | 3.45 | 2825975 |
| Ant_P3_30_2019_bin.28 | Chile_Bay | Bacteria; Proteobacteria; Gammaproteobacteria; Pseudomonadales; Alcanivoracaceae; PGZG01; | 1 | 0 | FALSE | FALSE | FALSE | FALSE | NODE_7872_length_3614_cov_2.993538_2 | FALSE | FALSE | FALSE | FALSE | FALSE | 70.64 | 3.13 | 2286974 |
| Ant_P3_30_2019_bin.3 | Chile_Bay | Bacteria; Proteobacteria; Gammaproteobacteria; Enterobacterales; Alteromonadaceae; Pseudoalteromonas; Pseudoalteromonas distincta | 1 | 0 | FALSE | FALSE | FALSE | FALSE | NODE_17894_length_1941_cov_5.053022_1 | FALSE | FALSE | FALSE | FALSE | FALSE | 55.17 | 6.90 | 2271457 |
| Ant_P3_30_2019_bin.25 | Chile_Bay | Bacteria; Proteobacteria; Gammaproteobacteria; | 0 | 1 | FALSE | FALSE | FALSE | FALSE | FALSE | NODE_274_length_38879_cov_5.258448_12 | FALSE | FALSE | FALSE | FALSE | 56.66 | 3.45 | 3489082 |

| | | | | | | | | | | | | | | | | | |
|---|---|---|---|---|---|---|---|---|---|---|---|---|---|---|---|---|---|
| | | Pseudomonadales; Pseudohongiellaceae; ; | | | | | | | | | | | | | | | |
| TARA_PSE_MAG_00104 | TARA_Delmont_v48 | Bacteria; Proteobacteria; Gammaproteobacteria; Enterobacterales; Alteromonadaceae; Marisediminitalea; Marisediminitalea aggregata | 0 | 1 | TRUE | FALSE | FALSE | FALSE | FALSE | TARA_PSE_MAG_00104_000000000039_34 | TARA_PSE_MAG_00104_000000000078_97 | FALSE | FALSE | FALSE | 94.55 | 0.74 | 4696093 |
| TARA_ANW_MAG_00065 | TARA_Delmont_v16 | Bacteria; Myxococcota; XYA12-FULL-58-9; XYA12-FULL-58-9; XYA12-FULL-58-9; ; | 0 | 0 | TRUE | FALSE | FALSE | FALSE | FALSE | FALSE | TARA_ANW_MAG_00065_000000000245_2 | FALSE | FALSE | FALSE | 88.61 | 5.97 | 5058444 |
| Ant_ICE_2016_bin.5 | Chile_Bay | Bacteria; Proteobacteria; Gammaproteobacteria; Burkholderiales; Burkholderiaceae; Hydrogenophaga; | 0 | 0 | TRUE | FALSE | TRUE | TRUE | FALSE | FALSE | NODE_2866_length_9146_cov_4.844902_4 | FALSE | NODE_2866_length_9146_cov_4.844902_2 | NODE_2866_length_9146_cov_4.844902_3 | 80.77 | 2.41 | 3417611 |
| Genome_042 | PM-RGC | Bacteria; Proteobacteria; Gammaproteobacteria; Burkholderiales; Burkholderiaceae; Hydrogenophaga; Hydrogenophaga sp002840835 | 1 | 1 | TRUE | TRUE | TRUE | TRUE | Genome_042_k141_390349_8 | Genome_042_k141_827382_98 | Genome_042_k141_547018_6 | Genome_042_k141_547018_4 | Genome_042_k141_547018_5 | Genome_042_k141_547018_13 | 98.60 | 0.40 | 5665796 |
| Genome_041 | PM-RGC | Bacteria; Proteobacteria; Gammaproteobacteria; Burkholderiales; Burkholderiaceae; Rubrivivax; | 1 | 3 | TRUE | FALSE | FALSE | FALSE | Genome_041_k141_933210_5 | Genome_041_k141_20164_5;Genome_041_k141_507163_8;Genome_041_k141_514530_2 | Genome_041_k141_590147_4 | | FALSE | FALSE | 89.46 | 1.36 | 4478983 |
| Genome_020 | PM-RGC | Bacteria; Proteobacteria; Alphaproteobacteria; Rhodobacterales; Rhodobacteraceae; Planktomarina; | 0 | 0 | TRUE | FALSE | FALSE | FALSE | FALSE | FALSE | Genome_020_k141_758178_22 | | FALSE | FALSE | 91.00 | 2.84 | 2372087 |

### Supplementary Table S9. Composition of Genome_042 MAG.

| Target | Context Protein | Relative Position | Definition |
|---|---|---|---|
| k141_827382_98 | k141_827382_93 | -5 | 4-hydroxybenzoyl-CoA thioesterase [EC:3.1.2.23] |
| k141_827382_98 | k141_827382_94 | -4 | branched-chain amino acid transport system substrate-binding protein |
| k141_827382_98 | k141_827382_95 | -3 | branched-chain amino acid transport system permease protein |
| k141_827382_98 | k141_827382_96 | -2 | branched-chain amino acid transport system ATP-binding protein |
| k141_827382_98 | k141_827382_97 | -1 | branched-chain amino acid transport system ATP-binding protein |
| k141_827382_98 | k141_827382_98 | 0 | mono(ethylene terephthalate) hydrolase [EC:3.1.1.102] |
| k141_827382_98 | k141_827382_99 | 1 | 3-(methylsulfanyl)propanoyl-CoA dehydrogenase [EC:1.3.99.41] |
| k141_827382_98 | k141_827382_100 | 2 | MarR family transcriptional regulator, lower aerobic nicotinate degradation pathway regulator |
| k141_827382_98 | k141_827382_101 | 3 | apolipoprotein N-acyltransferase [EC:2.3.1.269] |
| k141_827382_98 | k141_827382_104 | 4 | insecticidal toxin complex protein TccC |
| k141_827382_98 | k141_827382_109 | 5 | ribonuclease T1 [EC:4.6.1.24] |
| k141_547018_6 | k141_547018_1 | -5 | two-component system, OmpR family, copper resistance phosphate regulon response regulator CusR |
| k141_547018_6 | k141_547018_2 | -4 | transposase, IS5 family |
| k141_547018_6 | k141_547018_3 | -3 | IclR family transcriptional regulator, pca regulon regulatory protein |
| k141_547018_6 | k141_547018_4 | -2 | terephthalate 1,2-dioxygenase oxygenase component alpha subunit [EC:1.14.12.15] |
| k141_547018_6 | k141_547018_5 | -1 | terephthalate 1,2-dioxygenase oxygenase component beta subunit [EC:1.14.12.15] |
| k141_547018_6 | k141_547018_6 | 0 | CDP-4-dehydro-6-deoxyglucose reductase, E3 [EC:1.17.1.1] |
| k141_547018_6 | k141_547018_7 | 1 | 4-methoxybenzoate monooxygenase (O-demethylating) [EC:1.14.99.15] |
| k141_547018_6 | k141_547018_8 | 2 | ferredoxin, 2Fe-2S |
| k141_547018_6 | k141_547018_9 | 3 | 3-phenylpropionate/trans-cinnamate dioxygenase ferredoxin reductase component [EC:1.18.1.3] |
| k141_547018_6 | k141_547018_10 | 4 | 3-oxoacyl-[acyl-carrier protein] reductase [EC:1.1.1.100] |
| k141_547018_6 | k141_547018_11 | 5 | succinate-semialdehyde dehydrogenase / glutarate-semialdehyde dehydrogenase [EC:1.2.1.16 1.2.1.79 1.2.1.20] |
| k141_547018_4 | k141_547018_1 | -3 | two-component system, OmpR family, copper resistance phosphate regulon response regulator CusR |
| k141_547018_4 | k141_547018_2 | -2 | transposase, IS5 family |
| k141_547018_4 | k141_547018_3 | -1 | IclR family transcriptional regulator, pca regulon regulatory protein |

| | | | |
|---|---|---|---|
| k141_547018_4 | k141_547018_4 | 0 | terephthalate 1,2-dioxygenase oxygenase component alpha subunit [EC:1.14.12.15] |
| k141_547018_4 | k141_547018_5 | 1 | terephthalate 1,2-dioxygenase oxygenase component beta subunit [EC:1.14.12.15] |
| k141_547018_4 | k141_547018_6 | 2 | CDP-4-dehydro-6-deoxyglucose reductase, E3 [EC:1.17.1.1] |
| k141_547018_4 | k141_547018_7 | 3 | 4-methoxybenzoate monooxygenase (O-demethylating) [EC:1.14.99.15] |
| k141_547018_4 | k141_547018_8 | 4 | ferredoxin, 2Fe-2S |
| k141_547018_4 | k141_547018_9 | 5 | 3-phenylpropionate/trans-cinnamate dioxygenase ferredoxin reductase component [EC:1.18.1.3] |
| k141_547018_5 | k141_547018_1 | -4 | two-component system, OmpR family, copper resistance phosphate regulon response regulator CusR |
| k141_547018_5 | k141_547018_2 | -3 | transposase, IS5 family |
| k141_547018_5 | k141_547018_3 | -2 | IclR family transcriptional regulator, pca regulon regulatory protein |
| k141_547018_5 | k141_547018_4 | -1 | terephthalate 1,2-dioxygenase oxygenase component alpha subunit [EC:1.14.12.15] |
| k141_547018_5 | k141_547018_5 | 0 | terephthalate 1,2-dioxygenase oxygenase component beta subunit [EC:1.14.12.15] |
| k141_547018_5 | k141_547018_6 | 1 | CDP-4-dehydro-6-deoxyglucose reductase, E3 [EC:1.17.1.1] |
| k141_547018_5 | k141_547018_7 | 2 | 4-methoxybenzoate monooxygenase (O-demethylating) [EC:1.14.99.15] |
| k141_547018_5 | k141_547018_8 | 3 | ferredoxin, 2Fe-2S |
| k141_547018_5 | k141_547018_9 | 4 | 3-phenylpropionate/trans-cinnamate dioxygenase ferredoxin reductase component [EC:1.18.1.3] |
| k141_547018_5 | k141_547018_10 | 5 | 3-oxoacyl-[acyl-carrier protein] reductase [EC:1.1.1.100] |
| k141_547018_13 | k141_547018_8 | -5 | ferredoxin, 2Fe-2S |
| k141_547018_13 | k141_547018_9 | -4 | 3-phenylpropionate/trans-cinnamate dioxygenase ferredoxin reductase component [EC:1.18.1.3] |
| k141_547018_13 | k141_547018_10 | -3 | 3-oxoacyl-[acyl-carrier protein] reductase [EC:1.1.1.100] |
| k141_547018_13 | k141_547018_11 | -2 | succinate-semialdehyde dehydrogenase / glutarate-semialdehyde dehydrogenase [EC:1.2.1.16 1.2.1.79 1.2.1.20] |
| k141_547018_13 | k141_547018_12 | -1 | TRAP-type transport system periplasmic protein |
| k141_547018_13 | k141_547018_13 | 0 | 1,2-dihydroxy-3,5-cyclohexadiene-1,4-dicarboxylate dehydrogenase [EC:1.3.1.53] |
| k141_547018_13 | k141_547018_14 | 1 | isovaleryl-CoA dehydrogenase [EC:1.3.8.4] |
| k141_547018_13 | k141_547018_15 | 2 | TetR/AcrR family transcriptional regulator, cholesterol catabolism regulator |
| k141_547018_13 | k141_547018_16 | 3 | pivalyl-CoA mutase small subunit [EC:5.4.99.-] |
| k141_547018_13 | k141_547018_17 | 4 | pivalyl-CoA mutase large subunit [EC:5.4.99.-] |
| k141_547018_13 | k141_547018_18 | 5 | GTPase [EC:3.6.5.-] |

### SUPPLEMENTARY REFERENCES

1. Yoshida S, Hiraga K, Takehana T, Taniguchi I, Yamaji H, Maeda Y, et al. A bacterium that degrades and assimilates poly(ethylene terephthalate). Science. 2016;351:1196–9.

2. Knott BC, Erickson E, Allen MD, Gado JE, Graham R, Kearns FL, et al. Characterization and engineering of a two-enzyme system for plastics depolymerization. Proc Natl Acad Sci U S A. 2020;117:25476–85.

3. Sasoh M, Masai E, Ishibashi S, Hara H, Kamimura N, Miyauchi K, et al. Characterization of the terephthalate degradation genes of Comamonas sp. strain E6. Appl Environ Microbiol. 2006;72:1825–32.

4. Shigematsu T, Yumihara K, Ueda Y, Numaguchi M, Morimura S, Kida K. Delftia tsuruhatensis sp. nov., a terephthalate-assimilating bacterium isolated from activated sludge. Int J Syst Evol Microbiol. 2003;53:1479–83.

5. Chain PSG, Denef VJ, Konstantinidis KT, Vergez LM, Agulló L, Reyes VL, et al. Burkholderia xenovorans LB400 harbors a multi-replicon, 9.73-Mbp genome shaped for versatility. Proc Natl Acad Sci U S A. 2006;103:15280–7.

6. Hara H, Stewart GR, Mohn WW. Involvement of a novel ABC transporter and monoalkyl phthalate ester hydrolase in phthalate ester catabolism by Rhodococcus jostii RHA1. Appl Environ Microbiol. 2010;76:1516–23.

7. Choi KY, Kim D, Sul WJ, Chae J-C, Zylstra GJ, Kim YM, et al. Molecular and biochemical analysis of phthalate and terephthalate degradation by Rhodococcus sp. strain DK17. FEMS Microbiol Lett. 2005;252:207–13.

8. Wang YZ, Zhou Y, Zylstra GJ. Molecular Analysis of Isophthalate and Terephthalate Degradation by Comamonas testosteroni YZW-D. Environ Health Perspect. 1995;103:9
